# Context-dependent regulatory networks connect Alzheimer’s disease genetics to microglial inflammatory responses

**DOI:** 10.64898/2026.08.24.746572

**Authors:** Ting-Ting Fu, Margareta Kurkela, Jiayong Tu, Jiayi Zhang, Na Sun, Lindsay A. Farrer, TCW Julia, Lei Hou

## Abstract

Inflammation is central to Alzheimer’s disease (AD) pathogenesis. Microglia, the resident innate immune cells of the brain, exhibit diverse inflammatory states and are enriched for AD-associated genetic variants within active *cis*-regulatory elements (CREs). However, the interplay among genetic variants, transcription factor (TF)–CRE– gene programs, and microglial responses across inflammatory and disease contexts remain poorly understood. Here, we develop context-dependent epigenomic networks (*cEpiNets*), integrating bulk and single-nucleus assay for transposase-accessible chromatin using sequencing (ATAC-seq) to reconstruct regulatory programs across inflammatory, genetic perturbation, and disease contexts. Leveraging TF footprinting and graph embedding, *cEpiNets* identifies shared and context-specific programs and predicts regulatory circuits in unseen biological contexts. In a *SORL1*-marked inflammatory microglial state that expands during AD progression, *cEpiNets* annotates AD risk variants at the *SORL1* locus and identifies variants associated with cellular state abundance across donors. Cross-context analysis further identifies ZBTB14, whose inflammation-associated program connects AD risk variant–harboring CREs to target genes and widespread TF remodeling in AD. Donor-level ZBTB14 footprint activity is negatively associated with AD pathology, while combined IFNγ/TNFα stimulation represses *ZBTB14* and activates a subset of inferred targets. Collectively, *cEpiNets* bridges genetic variation, regulatory programs, and disease-associated cellular phenotypes to facilitate mechanistic interpretation of complex disease genetics.

## Introduction

Inflammation is increasingly recognized as a central driver of Alzheimer’s disease (AD) pathogenesis, actively influencing disease initiation and progression^1^. Microglia integrate diverse inflammatory cues elicited by cytokines, proteinopathy (e.g., amyloid-β and tau), vascular dysfunction, and other disease-associated insults. Rather than eliciting a uniform activation program, these perturbations engage distinct yet partially overlapping transcription factors (TFs) that orchestrate specific microglial responses. For example, interferon signaling primarily engages STAT and IRF family TFs^2,3^, inflammatory cytokines and stress signals induce AP-1 (FOS/JUN) programs^4^, while C/EBP and lineage-determining ETS factors such as PU.1 cooperate with these signal-dependent TFs to shape microglial inflammatory and disease-associated states^5–7^, recognized by recent single-cell transcriptomic studies^8,9^. Although controlled *in vitro* experiments provide mechanistic insight into downstream alterations for each response, how these distinct regulatory responses induce heterogeneous microglial dysfunctions observed in human AD remains largely unknown.

Human genetics provides an independent line of evidence implicating microglia as central mediators of AD susceptibility. Genome-wide association studies (GWASs) have consistently shown that AD risk variants are highly enriched within regulatory elements active in human microglia^10,11^, and functional studies have demonstrated that perturbation of established AD risk genes, such as *TREM2*^12^*, APOE*^13^*, INPP5D*^14^*, PLCG2*^15^, and *SORL1*^16^, reshapes microglial activation, inflammatory signaling, phagocytosis, or disease-associated cellular states. These observations suggest that inflammatory signaling and inherited genetic variation may converge on shared transcriptional regulatory mechanisms that ultimately determine microglial phenotypes during AD. However, given that genetic effects of disease-associated variants may depend on disease, cellular, or inflammatory states^17,18^, how inflammatory perturbations interact with AD genetic risk to remodel microglial regulatory programs remains poorly understood. Epigenomic profiling, such as assay for transposase-accessible chromatin using sequencing (ATAC-seq), provides a powerful data source both to dissect TF-CRE programs across conditions and annotate non-coding genetic variates associated with diseases. Recent single-nucleus ATAC-seq (snATAC-seq) studies have revealed widespread remodeling of chromatin accessibility associated with loss of cell identity during AD, referred to as epigenomic erosion^10,19^. However, compared with the rich diversity of microglial states consistently resolved by single-nucleus transcriptomic studies^8,9^, current single-nucleus epigenomic analyses typically revealed much less microglia clusters^8,20^, leaving many AD-associated transcriptional states without clear regulatory counterparts. Furthermore, perturbation models^21–24^ and human AD have largely been studied in parallel, with few efforts to systematically integrate perturbation-derived regulatory programs with human AD microglial states, which reflect both inflammatory signals and genetic risk.

To bridge these gaps, we present *<u>c</u>ontext-dependent <u>Epi</u>genomic <u>Net</u>work<u>s</u>* (*cEpiNets*), a framework that systematically models TF–CRE–gene regulatory circuits across diverse cellular contexts, using microglia as a proof-of-concept. We hypothesize that microglia exhibit numerous transcriptional states in AD, reflecting dynamic TF–CRE–gene regulation under various genetic and environmental perturbations, and these dynamics can be effectively modeled by a collection of context-dependent epigenomic networks.

We curated available ATAC-seq datasets, comprising bulk ATAC-seq of human microglia-like cells across 21 contexts^24–30^, i.e. different perturbation conditions, and snATAC-seq profiling of human microglia from AD (n = 12) and control donors (n = 8)^31^ (**Fig. 1a** and **Table S1**). Most of these contexts are relevant to AD such as perturbation conditions related to AD risk genes (e.g., *SORL1* and *APOE4*), and non-genetic stimuli such as interferon (IFN)-***β*** stimulation, enabling us to capture the microglial regulation during AD pathogenesis. Following uniform preprocessing and quality control for each context pair (see **Methods**), we calculated TF footprint scores to quantify binding profiles for 878 TFs and identified globally differential footprinting TFs (dfTFs) for each perturbation (**Fig. 1b**). Based on differential footprinting analysis, we linked 556 non-redundant dfTFs to over 191,292 differential footprinting CREs (dfCREs) across contexts and clustered these dfCREs into 100 modules (**Fig. 1c**). This strategy overcomes the sparsity inherent in snATAC-seq data (with only 87,548 CREs detected and poorly quantified in microglia) and the limited knowledge of TF binding landscape due to lack of TF ChIP-seq data for low-abundance cell types such as microglia. Furthermore, we developed a novel and flexible embedding-based framework to model CRE-gene linking (predicting ∼4 million pairs) based on *trans*-regulatory networks (**Fig. 1d**). This method outperforms existing correlation-based and genetic-based strategies in predicting chromatin interactions^32^. All the networks are available for download and interactive visualization at https://cepinets.thresh-infra.xyz/. Using *cEpiNets*, we dissect the regulatory circuits underlying AD-associated microglial inflammatory states (**Fig. 1e**) and annotated context-dependent *cis*- and *trans*-regulation programs for the AD risk gene *SORL1* and their association with AD-associated microglial inflammatory states (**Fig. 1f).** We also identified epigenomic regulatory circuits of inflammatory TFs, such as ZBTB14, that may drive dysregulated TFs under AD-related contexts (**Fig. 1g**). Overall, we established *cEpiNets* as a generalizable computational framework applicable to other cell types and diseases.

**Fig. 1.**
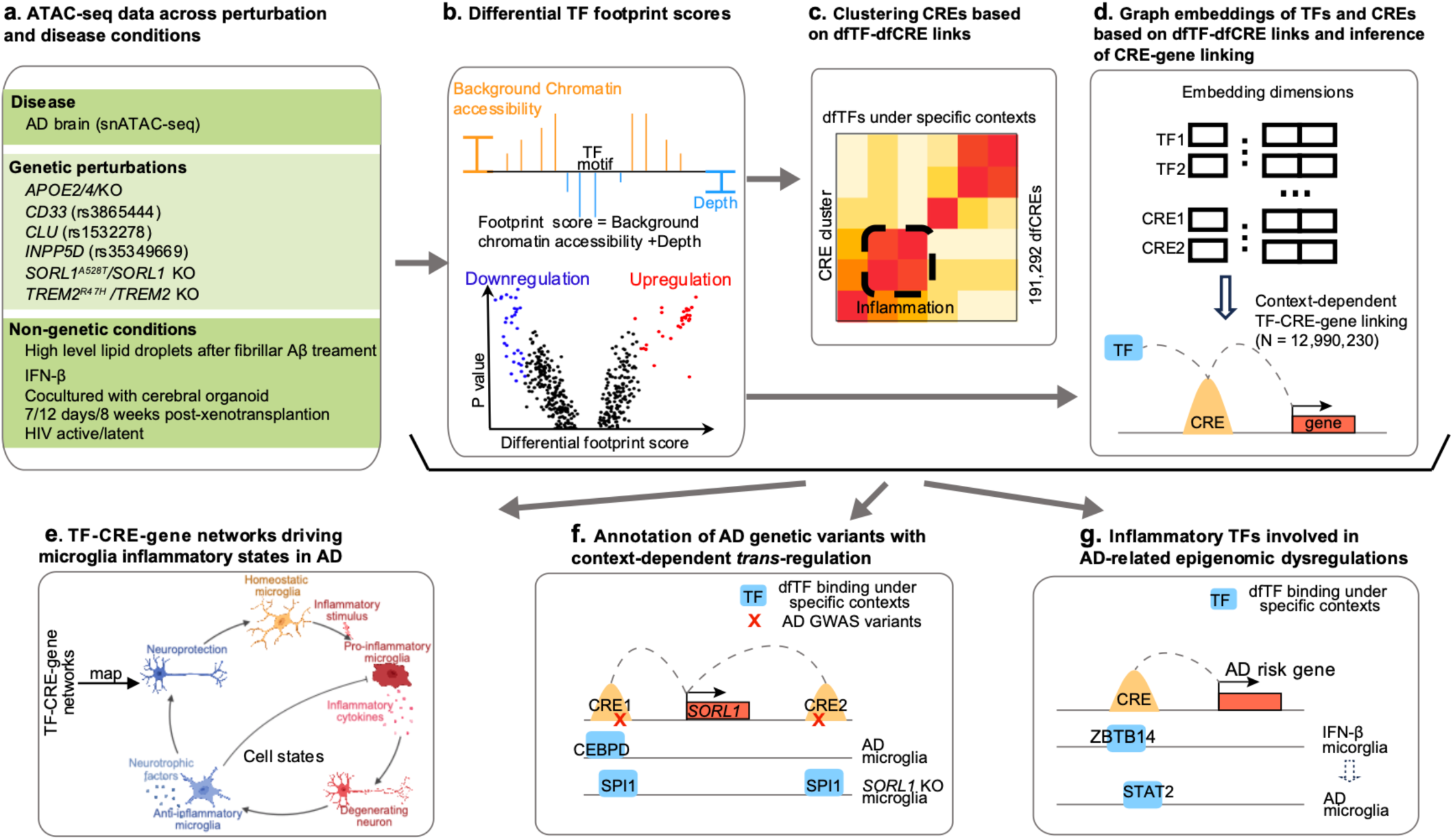
Overview of context-dependent epigenetic networks (*cEpiNets*) for microglia. **a.** Publicly available snATAC-seq datasets of human AD microglia and ATAC-seq of microglia-like cells under various perturbations and contexts were curated (**Table S1**). **b.** TF footprint scores and differential TF footprint scores between contexts and paired controls. **c.** dfCREs linked to dfTFs based on differential TF footprint scores under each context, and then clustered based on these links. **d.** Context-dependent TF-CRE-gene regulatory programs inferred based on graph embeddings. **e.** Critical regulatory circuits driving transcriptional programs of AD-associated inflammatory states. **f.** Annotation of *cis*- and *trans*-regulation programs for the AD risk variants at the *SORL1* locus and their association with microglial inflammatory states. **g.** Inflammation-associated regulatory programs modulated by ZBTB14 maybe involved in AD-related epigenomic dysregulation.

## Results

### An atlas of the context-dependent Epigenomic Networks (*cEpiNets*) for microglia

We constructed *cEpiNets* for microglia by integrating publicly available snATAC-seq data from AD microglia^31^ and bulk ATAC-seq data from human microglia-like cells under various contexts (**Fig. 1**). These contexts, the majority of which are associated with AD, encompass genetic variants (e.g., *APOE4*) as well as non-genetic conditions (e.g., IFN-***β*** stimulation and HIV infection) (**Table S1**). In total, we annotate 191,292 CREs targeted by 556 non-redundant TFs across 22 context pairs, i.e. perturbations vs controls. Comparison of our CREs with another independent AD snATAC-seq dataset^9^, which detected 87,525 CREs in microglia, revealed that our atlas captures 92.5% (n = 80,937) of their microglial CREs. Furthermore, 42% of our CREs were *cEpiNets*-specific in microglia, 24% overlapped with their microglial CREs, and 34% overlapped with their CREs in other brain cell types (**Fig. 2a**). Notably, the *cEpiNets*-specific CREs were significantly enriched for AD-associated functions including *chronic inflammatory response*, *interferon-mediated regulation*, *neurotransmission*, *the neuropilin signaling pathway*, and *lipid transport across blood-brain barrier* (Two-sided Binomial test, multiple testing correction by the Benjamini–Hochberg (BH) procedure, adjusted *P* < 0.05) (**Data S1**). This indicates that our networks capture a significantly broader repertoire of context-specific CREs relevant to microglial function than typically resolved by snATAC-seq alone.

**Fig. 2.**
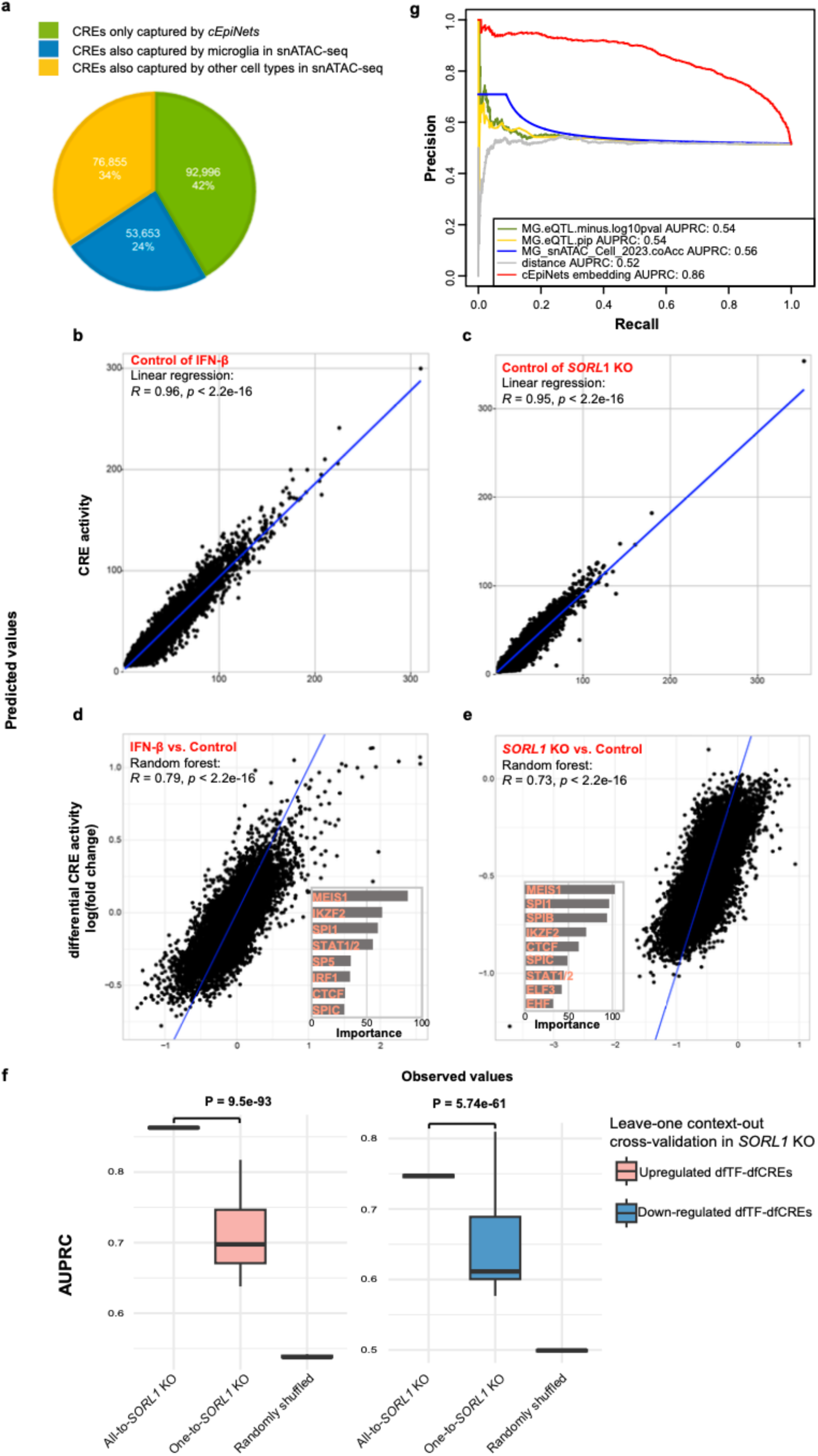
*cEpiNets* predicts CRE activity and differential activity based on footprinting and predicts CRE-genes linking with footprint-derived embedding. **a.** Mapping dfCREs from *cEpiNets* with those from AD snATAC-seq dataset (Xiong *et. al.*, Cell, 2023). **b-e.** Examples of prediction for CRE activity with multiple linear regression (**b,c**) and its differential signal with random forest regression (**d, e**) in contexts of IFN-*β* stimulation (**b, d**) and *SORL1* KO (**c, e**). Each dot represents a CRE, x-axis and y-axis represent the observed values and predicted values, respectively. The importance from the random forest model is descending sorted and the top 10 are shown in **d** and **e**. **f.** Embedding-based prediction for dfTF-dfCRE linkings in the target context of *SORL1* KO. The pink boxplots on the left are AUPRCs for predictions of dfTF-dfCREs based on upregulated binding and the blue boxplots on the right are for down-regulated ones. For upregulated or down-regulated dfTF-dfCREs, AUPRCs for predictions based on embeddings learned from all the context but the target (All-to-*SORL1* KO), embeddings from any other specific context (One-to-*SORL1* KO), or randomly shuffled embedded features (Randomly shuffled) were shown, respectively. **g.** Benchmarking embedding-based prediction of CRE-gene ABC scores with other methods on the hold-out chromosome, chr1 as an example. AUPRC is used for the assessment of prediction quality. The red line represents the embedding method used in *cEpiNets*. MG.eQTL.minus.log10pvalue: CRE-gene linking scored by max -log_10_(*P*) of eQTL of SNPs located in the CRE and targeting the gene in microglia; MG.eQTL.pip: scored by max eQTL posterior inclusion probability; MG_snATAC_Cell_2023.coAcc: scored by co-accessibility based on snATACseq data; distance: based on genomic distance between CRE and gene.

To map the dynamics of TF binding profiles across contexts, we utilized the TOBIAS pipeline^33^ to calculate TF footprint scores. We quantified binding profiles for 878 TFs in each context and identified the TFs with globally differential footprinting profiles (dfTFs) and their targeted differential footprinting CREs (dfCREs) relative to the paired controls (see **Methods**). We found that TF footprint scores could significantly predict both CRE activity in the same sample with multiple linear regression (Two-sided *t*-test, *P* < 0.05) (**Fig. S1-11**) and differential activity between paired conditions using a random forest model (Two-sided t-test, *P* < 0.05) (**Fig. S12-23**), with IFN-***β*** and *SORL1* KO shown as two examples (**Fig. 2b-e**, see **Methods**). We also identified top TFs contributing to the differential CRE activity based on variable importance derived from the random forest model. These included MEIS1, SPI1 (PU.1), SPIB, SPIC, STAT1/2, and CTCF both in IFN-***β*** stimulation and *SORL1* KO contexts, as well as IRF1 specifically under IFN-***β*** stimulation (**Fig. 2d and 2e**). Notably, SPI1, a master regulator of microglia with genetic links to AD^5,34^, was identified as a key driver. All of these highly important TFs but MEIS1 were also identified as dfTFs under *SORL1* KO or IFN-***β*** context (**Fig. 2a**), supporting the footprinting-based approach.

We subsequently linked dfTFs to dfCREs across contexts, establishing context-dependent epigenomic networks containing over 4 million non-redundant links (see **Methods, available from** https://huggingface.co/datasets/Thresh514/cEpiNets). We then projected TFs and CREs into an embedding space using *node2vec*^35^ to represent context-dependent dfTF-dfCRE linking and TF-TF protein interactions. To test whether embeddings learned capture biologically meaningful regulations, we first test whether they could predict dfTF-dfCRE linking in unseen contexts. We used a leave-one context-out cross-validation method (see **Methods**). We treated dfTF-dfCREs linking of one target context as true signals each time and used embedding learned for TFs and CREs from the rest contexts as features to predict the truth. For instance, in the target context of *SORL1* KO, the logistic regression model achieved mean Area Under the Precision-Recall Curves (AUPRCs) of 0.86 for upregulated dfTF-dfCREs and 0.75 for down-regulated ones, respectively (All-to-*SORL1* KO in **Fig. 2f**). The performance is significantly better than those either using the embeddings learned from one of the rest contexts (0.71 for upregulated dfTF-dfCREs and 0.65 for down-regulated dfTF-dfCREs) or using randomly shuffled embedded features of the rest contexts (0.54 for upregulated dfTF-dfCREs and 0.5 for down-regulated dfTF-dfCREs) (Two-sided Student’s t-test, *P* < 0.01) (**Fig. 2f**). The logistic regression models achieved mean AUPRCs of 0.81 for upregulated dfTF-dfCREs and 0.76 for down-regulated ones across all hold-out contexts, higher than those of 0.51 for upregulated dfTF-dfCREs and 0.52 for down-regulated ones predicted by randomly shuffled embedded features (**Fig. S24-S25**). These results supported that embedding learned across multiple contexts potentially captured dynamic regulations of TFs and CREs, thus being able to predict their links under new contexts.

Leveraging these embeddings of TF and CRE from networks, we further predicted the target genes for each dfCRE based on the premise that functionally linked CREs and gene promoters may share similar TF/cofactor regulatory environments, mediated through either direct TF binding or protein–protein interactions^36^. We took the difference between embedded vectors of CRE-gene pairs as features and used the generalized linear model to predict CRE-gene linking scores derived from microglia Hi-C data^11^, which profiles long-range enhancer–promoter contacts (see **Methods**). Compared to other methods including eQTLs^37^, genomic distance, or co-accessibility based on snATAC-seq^8^, our embedding-based prediction *cEpiNets* achieved superior performance with an AUPRC of 0.86 (**Fig. 2g**), validating the dfTF-CRE-gene regulatory networks used in subsequent analyses.

### TF regulation is context-dependent yet interconnected across contexts

We first compared dfTF identified in AD to those from other contexts. Upregulated dfTFs in AD microglia included those related to inflammatory response (e.g., FOS and JUN families), innate immune regulation (e.g., SPI1), C/EBP family members associated with myeloid activation (e.g., CEBPB, CEBPE, and CEBPG), and ETS-family (e.g., EHF) (**Fig. 3a**). The observed upregulation of SPI1 aligns with previous findings^31^. Intriguingly, several upregulated AD dfTF families, such as FOS, JUN, and ETS (e.g., EHF) families, appeared frequently across other contexts (**Fig. 3b**, **Fig. S26-S36**), suggesting their pivotal roles in microglial function and various stimulus responses. However, the directionality of their binding activity varied by contexts (**Fig. 3a**). For instance, FOS and JUN family members, early transcription factors that mediate inflammatory and stress-responsive gene expression, exhibited increased binding in AD, IFN-***β*** stimulation, and *APOE4* allele contexts, but reduced binding in *SORL1* KO, consistent with impaired endolysosomal trafficking and attenuated acute inflammatory activation associated with *SORL1* deficiency^38,39^. Conversely, the ETS family (e.g., SPI1, SPIB, SPIC, EHF) was upregulated in AD and downregulated under IFN-***β*** stimulation and *SORL1* KO (**Fig. 3a**), reflecting distinct engagement of ETS-dependent microglial programs during chronic disease remodeling compared to other contexts. CTCF, a key organizer of three-dimensional chromatin architecture, exhibited reduced binding in AD, consistent with previous evidence of diminished CTCF occupancy in aged and AD brains^20,40^, and showed upregulated binding in *APOE4* allele and no significant change under IFN-***β*** stimulation nor *SORL1* KO (**Fig. 3a**). These distinct patterns showed each context might only recapitulate a specific aspect of AD dysregulation, suggesting the need for a more systematic view of regulatory dynamics.

**Fig. 3.**
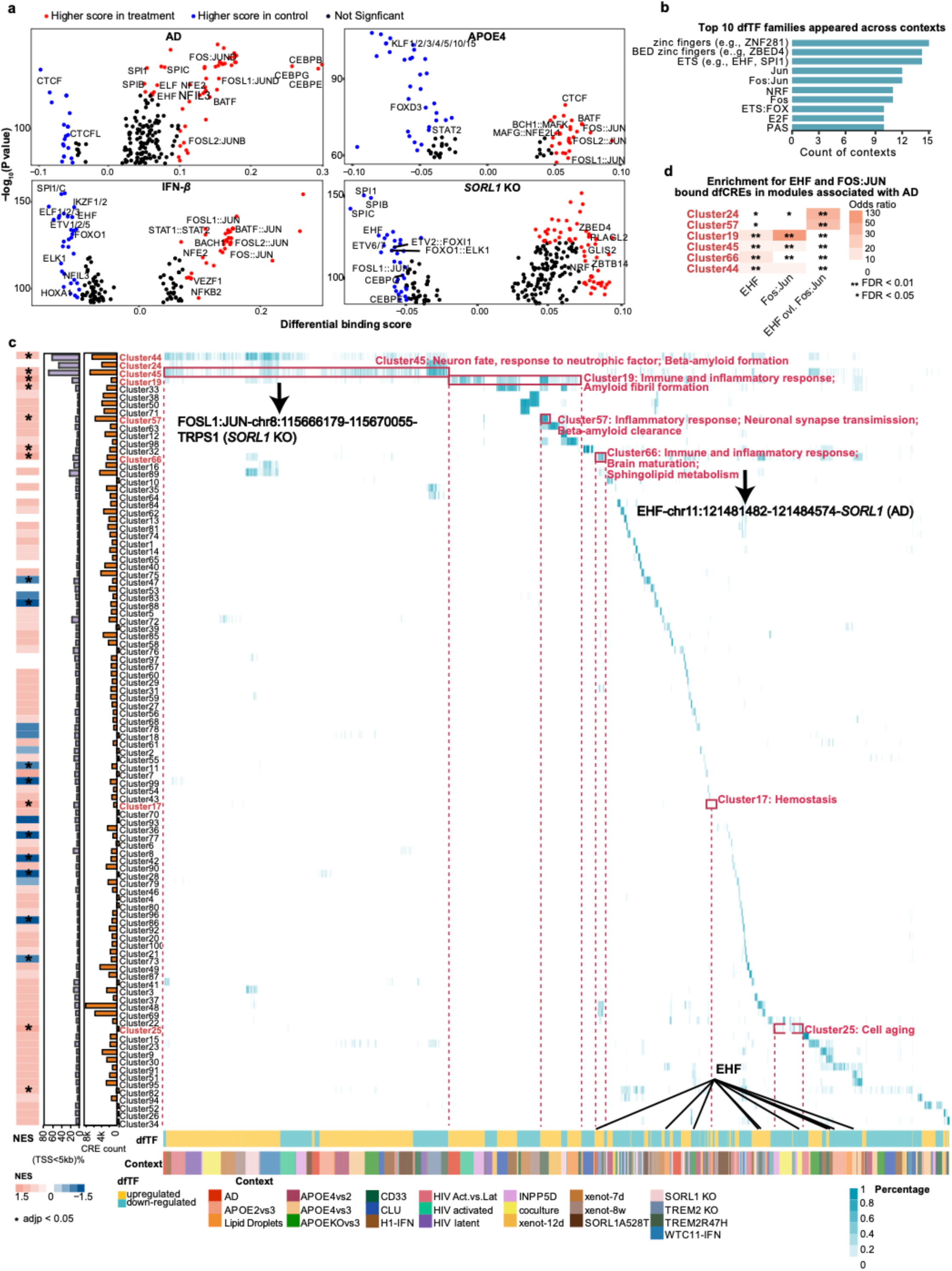
dfTFs and dfCRE modules across contexts. **a.** Differential TF footprinting across contexts. The volcano plots show the overall footprinting of dfTFs in AD microglia, microglia with IFN-***β*** stimulation, *APOE4* allele and *SORL1* KO, respectively. Each dot represents a dfTF. Red indicates a significantly higher binding score after treatment while blue indicates a significantly lower binding score. Black dots are non-significant dfTFs. **b.** Top 10 dfTFs families appeared across contexts. **c.** CRE modules across TF regulation and contexts. The heatmap shows the differential binding activity by dfTFs across contexts (by column) for each dfCRE module of *cEpiNets* (by row). The left three panels in order are statistics of NES from GSEA enrichment analysis for AD-associated CRE modules, percentage of CREs located with TSS < 5kb, and the CRE counts in each dfCRE module. The dfCRE modules with asterisk are enriched in chromatin accessibility positively associated with AD. The representative biological functions for enriched modules are highlighted. Two representative circuits from their corresponding CRE modules, in the format of TF-CRE-gene (context) are shown below the black arrow. A dfTF EHF, as an example, was shown across contexts. **d.** The heatmap shows the enrichment of TFs EHF, FOS:JUN, and co-bound dfCREs in some of the modules associated with AD (asterisk marked in **c**).

To dissect the complicated epigenomic signatures of AD and connect them to various TF-directed programs we identified from other contexts, we clustered dfCREs of *cEpiNets* into 100 modules based on context-dependent dfTF-CRE networks using the Jaccard index as the distance metric (**Fig. 3c, Data S2**; see **Methods**). Integrating with CRE differential signals in AD microglia^10^ and performing enrichment analysis by GSEA^41^, we identified modules significantly enriched for AD-associated epigenomic signals (multiple hypothesis correction, adjusted *P* < 0.05), including Clusters 44, 45, 19, 57, 66, 17, and 25 (**Fig. 3c**). Functionally, AD epigenomic enriched modules were implicated in AD-related processes such as *neurotrophic factor response* and *beta-amyloid formation* (Cluster 44); *immune/inflammatory response* and *amyloid fibril formation* (Cluster 45); *sphingolipid metabolism* (Cluster 66); *cell hemostasis* (Cluster 17); and *cell aging* (Cluster 25) (**Fig. 3c**, **Data S3**).

Clusters 44 and 45 contained the largest number of dfCREs and were predominantly promoter-like, with over 70% of elements located within 5 kb of a transcription start site (TSS) (**Fig. 3c**, **Data S4, S5**). These promoter-like dfCRE modules showed consistently differential binding activity across a broad range of contexts, including AD microglia, AD risk variants (*SORL1*, *TREM2*, and *INPP5D*), HIV infection, and xenotransplantation (**Fig. 3c**). In contrast, the remaining dfCRE modules were largely enhancer-like (50–500 kb from TSS) and exhibited context-specific activity (**Fig. 3c**). Collectively, these results demonstrate that *cEpiNets* captures both context-shared (promoter-like) and specific (enhancer-like) epigenomic regulation underlying AD-associated biological processes.

We further examined dfTF regulation within specific dfCRE modules. We observed that ETS (e.g., EHF) and FOS/JUN families, upregulated in AD, not only appeared frequently in other contexts (**Fig. 3b**), but also showed significant enriched binding (Two-sided Fisher’s exact test *P* < 0.05) in dfCRE modules associated with AD epigenomic signals (e.g., Cluster 19 and 24) (**Fig. 3d**, **Fig. S37**). Moreover, EHF and FOS/JUN families exhibited extensive co-occupancy at the same dfCREs within these modules across contexts. For example, in AD microglia, dfCREs co-bound by EHF and FOS/JUN were enriched in Cluster 19, 45, 44, and 66 (Two-sided Fisher’s exact test, *P* < 0.01) (**Fig. 3d**). In *SORL1* KO, both TF families were downregulated compared with the control, and their co-bound dfCREs also showed enrichment in those AD-associated modules (**Table S2**). Given the reported anticooperative binding between EHF and the JUN–FOS AP-1 complex^42^, and that AP-1 regulates gene expression in response to various stimuli including inflammation^43^, our observation of shared binding sites of them enriched in AD-associated modules suggests that interactions among TFs are critical for shaping the context-dependent TF regulation in response to inflammatory stimulation in microglia.

### Epigenomic regulatory programs of *SORL1* drive inflammatory microglial states in AD

Microglia shows diverse transcriptional states in AD. Sun *et al.* reported that microglial states MG2, MG8, and MG10 represented inflammatory status, yet only MG8 showed a significantly increased fraction in AD^8^. Importantly, *SORL1* is both a marker gene in MG8 and an AD risk gene. *SORL1* KO impairs Aβ uptake both in a human microglia-like cell model and a mouse AD brain xenotransplant model^27^. We therefore sought to understand how *SORL1* modulates MG8, through epigenomic networks, utilizing the *SORL1* KO context we collected in *cEpiNets*.

We integrated context-dependent TF-CRE networks and CRE–gene linking derived from *cEpiNets*, and identified candidate TF-CRE-gene circuits that are enriched in marker genes of microglial transcriptional states defined by Sun *et al*.^8^ using GSEA (multiple testing correction by the BH procedure, adjusted *P* < 0.05, see **Methods**). We first tested whether AD and *SORL1* KO share dysregulated TF-directed programs associated with MG8. We found that several dfTFs programs positively enriched in MG8 were shared between AD microglia and *SORL1* KO contexts, including CEBP (e.g., DBP, HLF, and NFIL3), ETS (e.g., EHF, ELK1, and ETV2), and FOX (e.g., FOXO1 and FOXI1) families (**Fig. 4a**), implicating that *SORL1* KO and AD pathogenesis may share these dysregulation in modulating microglial inflammatory states, e.g., MG8. To mitigate potential bias from cell state definitions, we validated our findings using the Human Microglia Atlas (*HuMicA*)^9^, which consolidates multiple microglia snRNA-seq and scRNA-seq datasets. *HuMicA* clustered the integrated object into nine clusters with annotation of 0 to 8 and calculated upregulated gene markers for each of them. Based on the enrichment of markers from *HuMicA* and those from MG8, *HuMicA* Cluster 0, 2, 4, and 8 were significantly enriched in the MG8 state (FDR < 0.05). Consistently, dfTFs from the AD and *SORL1* KO contexts (e.g., ATF, CEBP, FOS, JUN families, and MITF) were enriched in these *HuMicA* clusters, confirming their important roles in modulating the MG8 state (**Fig. S38-S39**).

**Fig. 4.**
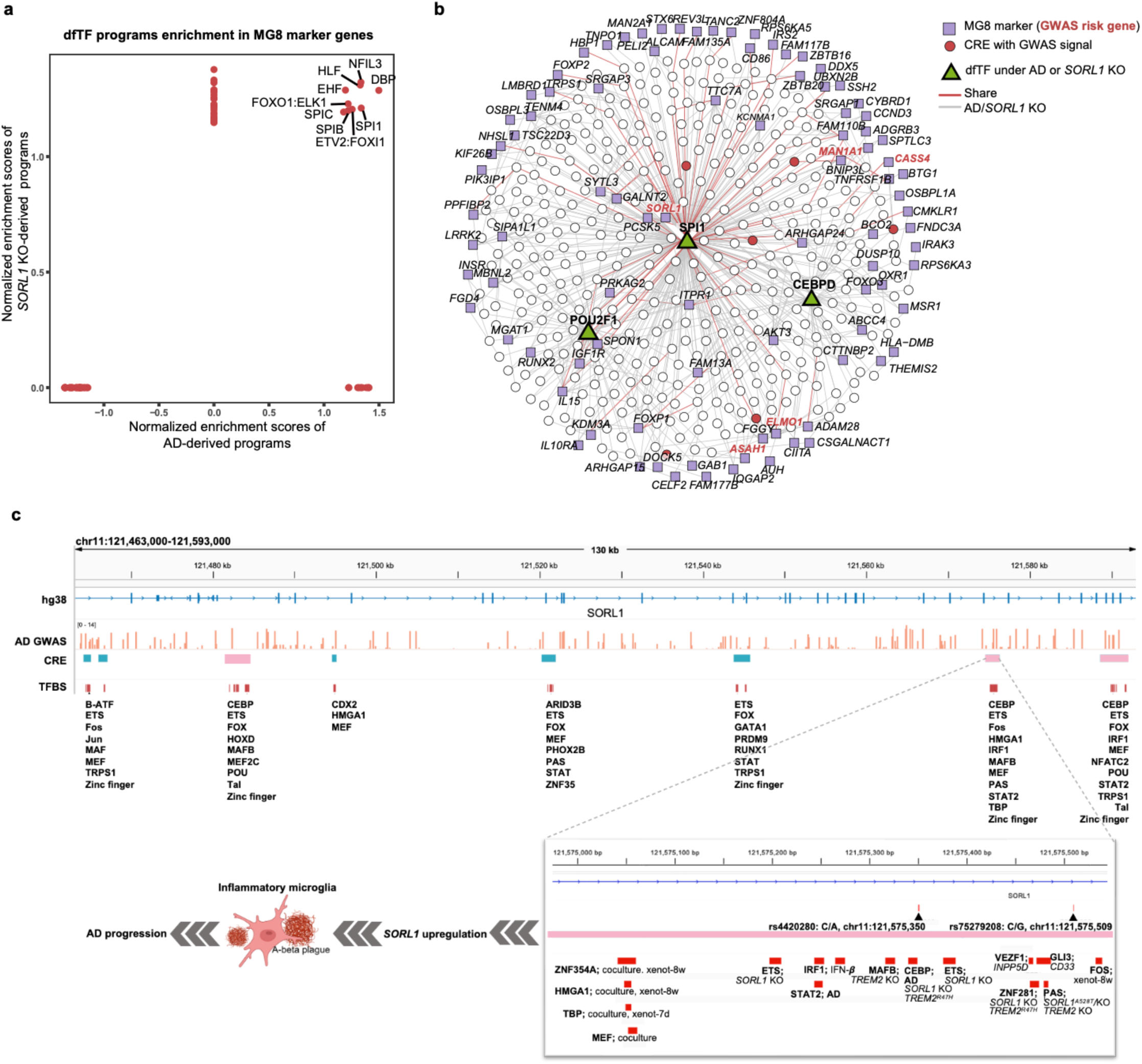
*SORL1-*related Epigenomic regulatory programs drive inflammatory microglial states in AD. **a.** Comparison of enrichment of TF-directed regulatory circuits derived from AD and *SORL1* KO microglia in marker genes of microglial MG8. X-axis and Y-axis represent normalized enrichment scores (NES) of circuits derived from AD microglia and *SORL1* KO context, respectively. **b.** MG8 enriched regulatory circuits from contexts of AD or *SORL1* KO. Green triangles represent TFs consistent with the previous report^8^. Purple squares represent MG8 markers. Gene symbols in red bold fonts represent marker genes are also known AD risk genes. Circles represent dfCREs, and the red ones are associated with AD risk variants. Red lines represent shared links between AD and *SORL1* KO and grey lines mean specific in one of two contexts. **c.** AD GWAS variants and related *trans* circuits *of* dfTF-dfCREs across contexts at the *SORL1* locus. dfCREs annotated by *cEpiNets* harboring AD GWAS risk variants at the *SORL1* locus were shown in turquoise trunk and those shared between AD and other contexts are highlighted in pink. Bound dfTF families at each dfCREs are listed below. The panel below zooms in one of the MG8-related dfCREs where GWAS variants and dfTFs from all contexts are located, potentially regulating *SORL1* expression and driving MG8 during AD progression. The plot is available from *cEpiNets* database https://cepinets.thresh-infra.xyz/.

We then leveraged these context-dependent TF–CRE–gene links to connect AD risk variants (AD GWAS^44^ nominal *P* < 1 x 10^-5^) within CREs to MG8 marker genes. Several MG8-enriched dfTFs identified by *cEpiNets* in AD or *SORL1* KO context recapitulated findings from motif enrichment analysis in Sun *et al.*’s report^8^, including CEBPD, POU2F1, and SPI1 (**Fig. 4b**). *SORL1* itself was among the predicted target genes, allowing *cEpiNets* to further resolve how regulatory elements at the *SORL1* locus are differentially engaged across disease and genetic perturbation contexts. In AD microglia, CEBPD showed increased binding at a dfCRE (chr11:121574474–121576278) containing the AD GWAS variants rs4420280, rs1620130, and rs75279208, whereas the same CRE showed increased SPI1 binding in *SORL1* KO. SPI1 also regulated *SORL1* via a distinct dfCRE (chr11:121588459-121591958) specifically in the *SORL1* KO context (**Fig. 4b**). This regulatory architecture extended beyond *SORL1*. These two dfTFs also regulated other AD risk genes such as *CASS4* and *ELMO1*, through distinct dfCREs (**Fig. 4b**), suggesting that MG8-associated regulators may coordinate multiple genetically implicated loci rather than act through a single AD gene. In addition, we identified novel MG8-enriched dfTFs programs shared between AD and *SORL1* KO that were not reported by Sun *et al*., including CEBP (e.g., CEBPA/B/E/G, DBP, HLF, NFIL3, TEF), ETS (e.g., EHF, ELF1/3/5, ELK1, ETV2, SPI1/B/C), FOX (FOXC1/I1/J1/O1/P1) families, and STAT2 (**Fig. S40**). They were predicted to target MG8 maker genes through AD genetic variants-containing CREs, including *SORL1, ASAH1*, *BNIP31, FGD4, MGAT1, PILRA,* and *PPFIBP2* (**Fig. S40**). Among them, the regulatory link, FOXP1–*SORL1*, is supported by prior literature^45^. Together, these findings suggest that regulation of the MG8 inflammatory state involves coordinated remodeling of CEBP-, ETS-, and FOX-centered programs across multiple genetically implicated AD loci, distinct from the FOS/JUN-dominated immediate-early programs prominent under acute inflammatory stimulation.

Given *SORL1* as an AD risk gene as well as its potential role in modulating AD-associated MG8 state, we zoomed into the regulation at the *SORL1* locus. We identified eight dfCREs annotated by *cEpiNets* harboring AD GWAS variants at the *SORL1* locus, three of which are shared between AD and other contexts (chr11:121481482-121484574, chr11:121574474-121576278, chr11:121588459-121591958) (**Fig. 4c, Data S6**). ETS, CEBP, FOS, and JUN families (**Fig. 4c**) dominated binding at these dfCREs, also identified as key TFs in response to acute inflammation above. However, none of the risk variants at these dfCREs are associated with *SORL1* expression in microglia^8^ (linear regression, *P* > 0.05), highlighting our context-resolved approach provides a complementary way to bridge the genetic variants and target genes compared with eQTLs, which may only capture baseline level effects^46^. We further analyzed the correlation between AD GWAS variants at the *SORL1* locus (*P* < 1 x 10^-5^) and the MG8 cell proportion. A total of eleven variants were significantly positively associated with MG8 proportion (linear regression, adjusted *P* < 0.05, **Data S7**). This included two variants (rs4420280 and rs1620130) located in the dfCRE at chr11:121574474-121576278 (Zoomed in plot, **Fig 4c**), a site previously identified as a microglial super-enhancer^47^. AD variant rs4420280 exhibited allelic imbalance in chromatin accessibility^48^, indicating its potential role in regulating the MG8 through *SORL1*. The dfTFs binding at the dfCRE are mainly from CEBP (e.g., CEBPA/B/D/E/G, HLF, TEF), ETS (e.g., EHF), and FOS families (**Fig. 4c**). Collectively, these data suggest that these AD variants at the *SORL1* locus (e.g., rs4420280, rs1620130) and associated TFs (e.g., CEBP, ETS, and FOS families) binding at this specific dfCRE (chr11:121574474-121576278) may regulate *SORL1* expression under inflammatory conditions, thereby driving MG8 and contributing to AD pathogenesis (**Fig. 4c**).

### Upregulated inflammatory TF binding shapes the epigenomic landscape towards an AD-like state

Persistent inflammatory states represent a prominent component of microglial dysregulation in AD^49^. Our analysis revealed both shared and divergent TF binding patterns between acute inflammatory state (IFN-***β*** stimulation) and potential chronic inflammatory state in AD microglia (**Fig. 3a**). TFs from FOS and JUN families were overall upregulated in both contexts, while ETS family (e.g., EHF, SPI1, SPIB, SPIC) exhibited decreased binding in acute inflammation yet increased binding in AD (**Fig. 3a**). Considering EHF may repress FOS and JUN responsive genes^42^, it suggests a dynamic repressive mechanism might be involved during the acute-chronic inflammatory transition. Given the emerging roles of transcriptional repressors in constraining inflammatory responses^50,51^, we thus asked whether rewiring of repressors-directed programs in pro-inflammatory microglia, including IFN-***β*** stimulation, *INPP5D* knockdown (KD), and HIV activation, could contribute to regulatory changes observed in AD. To test this, we examined whether dfCREs targeted by dfTFs in these contexts were preferentially located within 10 kb of dfCREs targeted by AD-associated dfTFs. We found that dfCREs targeted by inflammation-upregulated TFs with putative repressive activity, such as VEZF1 (IFN-***β*** and HIV activation), ZBED2 (IFN-***β***, HIV activation, and *INPP5D* KD), and ZBTB14 (IFN-***β*** and HIV activation), were significantly located near dfCREs targeted by AD dfTFs (multiple testing correction by the BH procedure, adjusted *P* < 0.05, **Fig. S41**). For instance, under IFN-***β*** stimulation, dfCREs with upregulated binding by ZBTB14 and ZBED2 were preferentially located near those targeted by AD-downregulated dfTFs (e.g., TFAP2 family, CTCFL, GLI1/2/3, and other zinc finger proteins) (**Fig. S41**), suggesting that inflammatory signaling engages repressive programs that may later dampen nearby TF binding, manifesting as AD-associated TF downregulation.

We next asked whether these spatially proximal regulatory elements targeted by various dfTFs also exhibited coordinated binding activity across individual samples. We then tested the correlation between the mean footprint scores of the dfCREs targeted by potential repressors in pro-inflammatory conditions and AD dfTFs across 238 samples with AD snATAC-seq data in microglia^10^ (see **Methods**). Two distinct repressor groups emerged (**Fig. 5a**, top panel). The first group included ZBED4, VEZF1, PATZ1, KLF15, ZBTB7B/C, ZBTB14, and NFKB2, whose binding score showed significantly positive correlations with AD-downregulated dfTF footprint scores and negative correlations with AD-upregulated dfTFs (*t*-test, multiple testing correction by the BH procedure, adjusted *P* < 0.05), suggesting potential homeostatic gate keepers. The second group includes ZBED2 and ZBTB33, showing the opposite pattern. These patterns suggested these potential repressors responding to pro-inflammation conditions may impact the binding of AD dfTFs nearby in microglia during AD pathogenesis.

**Fig. 5.**
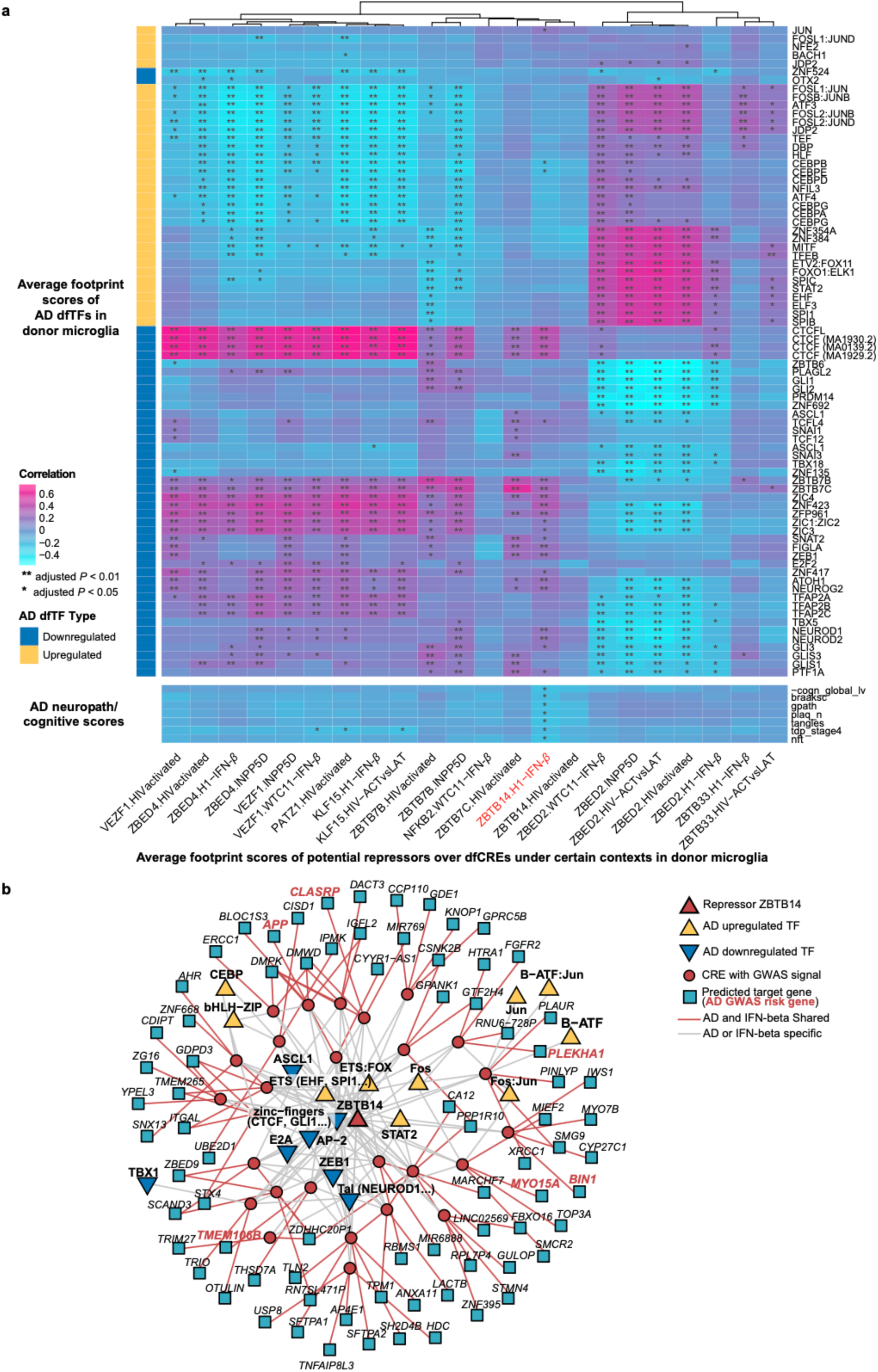
Upregulated bindings of inflammatory TFs shape the epigenomic landscape towards an AD-like state. **a.** Correlation among mean footprinting scores across dfCREs bound by the repressors under pro-inflammatory contexts and AD dfTFs based on microglia snATAC-seq data and AD cognitive scores across individuals. Rows are AD downregulated or upregulated dfTFs (top panel) or AD neuropath/cognitive scores (bottom panel), and each column represents a repressor in a pro-inflammatory context. AD upregulated and downregulated dfTFs are annotated with yellow and blue side bars, respectively. Braaksc, Braak score; gpath, global pathology score; plaq_n, amyloid plaque burden; nft, neurofibrillary tangle burden; tdp_stage4, four-stage measure of anatomical TDP-43 pathology distribution; cogn_global_lv, global cognition level. **b.** Repressor ZBTB14 (red triangle) co-bound with AD dfTFs at the same dfCREs with AD GWAS risk signals (red circles) and predicted target genes (turquoise squares). Yellow and blue triangles represent upregulated and downregulated dfTFs in AD, respectively. Red lines represent shared links between AD and IFN-***β*** stimulation.

Intriguingly, footprint scores of ZBTB14 from the first group are negatively correlated with AD neuropathology, including Braak score (braaksc), global pathology score (gpath), amyloid plaque burden (plaq_n), tangles, TDP-43 pathology (tdp_stage4), and neurofibrillary tangle burden (nft), while positively correlated with global cognition level (cogn_global_lv) (**Fig. 5a**, bottom panel), suggesting that ZBTB14 may mediate transcriptional repression, limit inflammatory remodeling in microglia, thereby reducing neuropathological burden and preserving cognitive function. Interestingly, ZBTB14 was also identified as an AD risk gene based on GWAS in African Americans^52^, and a negative regulator of SPI1^53^, which governs microglial neurotoxicity and neuroinflammatory response to amyloid plaques in AD^54^. We further interrogated the specific dfCREs cobound by ZBTB14 and AD dfTFs as well as their predicted targeted genes in AD with *cEpiNets*. Our results showed that in AD microglia, ZBTB14 and SPI1 were co-bound at dfCREs from AD-associated Cluster 44 (chr6:30907494-30908893, chr16:31072378-31075122) and Cluster 45 (chr7:12210196-12212328, chr7:17938864-17941508) with AD risk variants (*P* < 1×10^-5^) (**Fig. 5b, Data S8**). ZBTB14 also co-bound with other AD dfTFs (e.g., upregulated EHF, FOS, JUN, and CEBP families; downregulated CTCF, GLI1/2/3, and NEUROD1/2) at dfCREs harboring AD risk variants (**Fig. 5b, Data S8**). These circuits targeted genes included the AD risk genes *APP*, *BIN1*, *CLASRP*, *PLEKHA1*, and *TMEM106B* (**Fig. 5b, Data S8**). Specifically, ZBTB14 co-bound with SPI1 at a dfCRE (chr7:12210196-12212328), containing AD GWAS signals (SNPs rs7781670, rs1019309, and rs1019307, P < 1×10^-7^) and predicted to target *TMEM106B,* a regulator of lysosomal function implicated in neurodegenerative disease^55^. These results indicate that ZBTB14 active in acute inflammation may modulate how microglial epigenome transit toward a chronic inflammatory state in AD by modulating AD dfTF binding, particularly at CREs containing AD risk variants.

### *ZBTB14* is downregulated by AD brain prevalent proinflammatory cytokines in combination with differentially expressed genes

To investigate the response of ZBTB14 and its predicted regulatory targets under AD-relevant inflammatory conditions, we established an *in vitro* microglial inflammatory model using human microglial HMC3 cells stimulated with human IFNγ (100 ng/mL) and/or TNFα (50 ng/mL), which are prevalent in AD brains^56^, for 6 hours (**Fig. 6a; Methods**). We quantified the expression of *ZBTB14*, predicted *ZBTB14* target genes identified from the *cEpiNets* network (**Data S9**), and the inflammatory and AD-related marker genes by reverse transcription quantitative PCR (RT-qPCR). While cytokine treatments did not noticeably alter cell density or morphology (**Fig. S42a**), combined IFNγ and TNFα stimulation induced a robust inflammatory response, evidenced by a 2.95-fold increase in *PD-L1* expression^57^ (*P* < 0.0001) (**Fig. 6b**). Consistently, *IL6* and the NF-κB subunit *RELA* were significantly upregulated by 2.1-fold (*P* < 0.0001) and 2.5-fold (*P* = 0.0003), respectively **(Fig. 6c** and **6d**), confirming activation of inflammatory signaling by IFNγ and TNFα.

**Figure 6:**
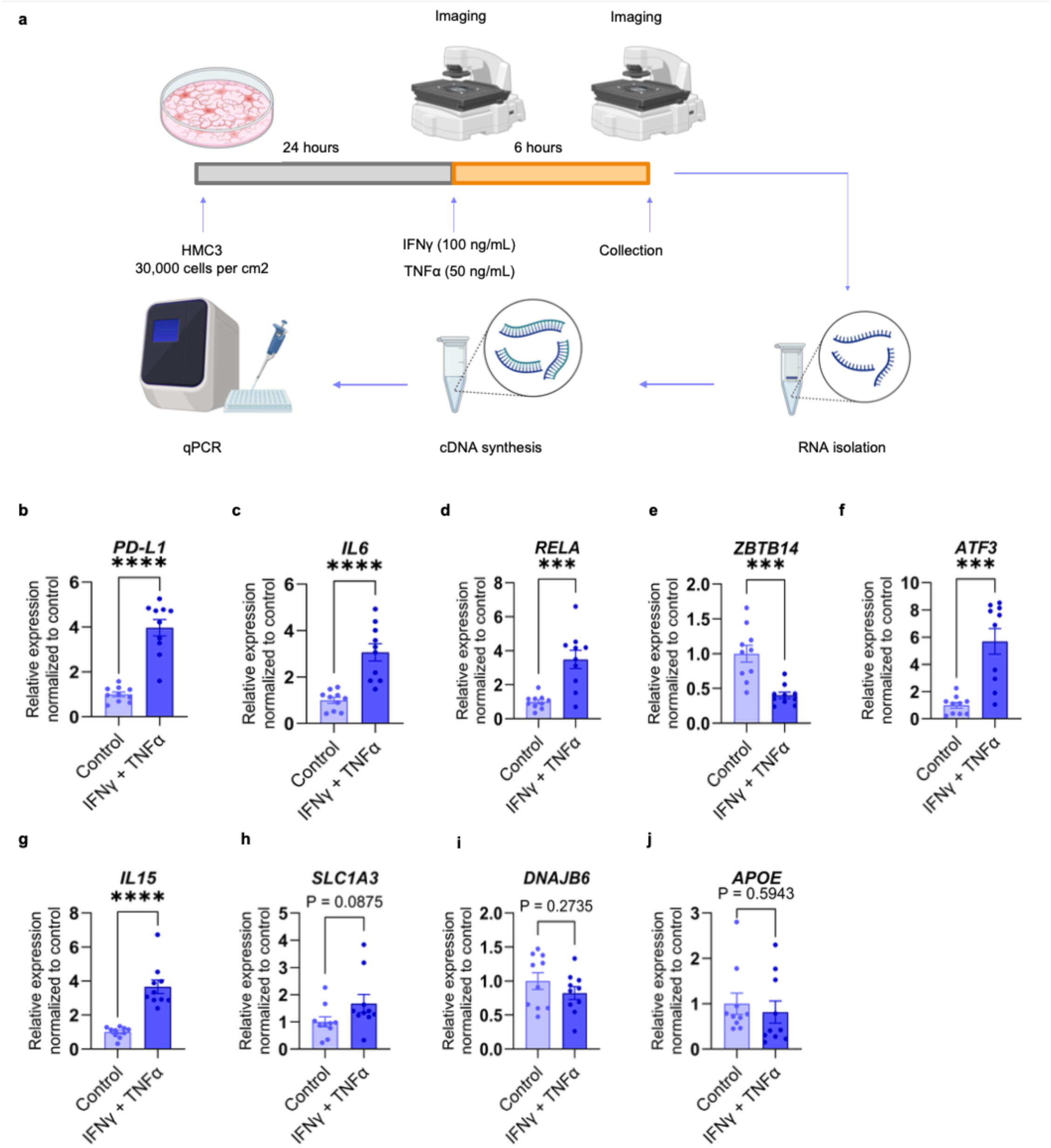
The relative gene expression of selected genes differentially expressed in connection with *ZBTB14*. **a.** Experimental design for the exposure of HMC3 cells to proinflammatory cytokines human IFNγ (100 ng/mL) and TNFα (50 ng/mL). Relative gene expressions of **b.** *PD-L1,* a positive control direct downstream gene of the treated cytokine and **c-d.** *IL6,* and NF-κB subunit *RELA*, genes involved in AD with a strong connection with inflammation, and **e.** *ZBTB14*, the gene encoding the target TF for this study. Relative expressions of **f.** *ATF3,* a gene involved in inflammation and associated with *ZBTB14,* **g.** *IL15*, a gene involved in AD with a strong connection with inflammation and associated with *ZBTB14***. h-i.** Relative expressions of *SLC1A3* and *DNAJB6,* genes involved in AD and associated with *ZBTB14,* and **j.** *APOE,* a gene having a strong connection with AD. Data plotted on graph as average ± standard deviation, each dot represents one biological replication from two technical replicates. n = 10 biological replicates per group \*\*\**P* < 0.001, \*\*\*\**P* < 0.0001 in student’s two-tailed t-test.

Under the same conditions, *ZBTB14* expression was reduced by approximately 60% (*P* = 0.0002) **(Fig. 6e**). Correspondingly, two predicted ZBTB14 target genes, *ATF3*, a stress-responsive transcription factor^58^, and *IL15*, a cytokine implicated in neuroinflammation^59^, were significantly induced by 4.7-fold (*P* = 0.0001) and 2.65-fold (*P* < 0.0001), respectively (**Fig. 6f** and **6g**). In contrast, other predicted targets including *SLC1A3* and *DNAJB6,* as well as *APOE* showed no significant changes (**Fig. 6h–6j**), suggesting selective activation of the ZBTB14 regulatory program.

Interestingly, IFNγ treatment alone was sufficient to induce *PD-L1* expression but did not significantly alter *IL6*, *RELA*, *ZBTB14, SLC1A3, or APOE* expression (**Fig. S42b-g**), indicating that repression of ZBTB14 is not a general consequence of an inflammatory stimulus. Instead, the loss of ZBTB14 and induction of its downstream regulatory program appear to require a more robust inflammatory context, such as under our combined treatment. Together with the negative association between ZBTB14 footprint activity and AD neuropathology observed in human postmortem microglia (**Fig. 5a**), these findings support a model in which ZBTB14 functions as a context-dependent transcriptional repressor whose loss permits activation of inflammatory regulatory programs during chronic inflammatory remodeling in AD microglia.

## Discussion

We present *cEpiNets*, including a comprehensive epigenomic atlas for microglia and a computational framework that deciphers context-dependent regulation via TF-CRE-gene triplets. Using *cEpiNets*, we found that dfTFs of the ETS, FOS, and Jun families showed upregulated binding profiles in AD and co-bound at dfCREs exhibiting AD-associated differential activity. These dfTFs, along with the majority of other AD-upregulated ones, display persistent differential binding across other AD-related contexts (e.g., IFN-***β*** stimulation, AD risk variant perturbations, and *APOE4* carriers), albeit through context-dependent CREs. FOS and JUN also respond to early inflammation in other cell types such as neurons^60^, suggesting their broad roles across cell types. Using *SORL1* KO as a proof-of-concept, we showed that EHF, FOS, and JUN families might modulate the effects of AD GWAS risk variants on gene expression, governing the inflammatory microglial state and influencing AD pathogenesis. We then leveraged *cEpiNets* to provide evidence that inflammation-associated repressors (e.g., ZBTB14, VEZF1, and ZBED2) may be associated with the binding of AD dfTFs at dfCREs harboring AD GWAS risk signals. Together with *in vitro* microglia models mimicking a sustained microglia inflammatory state, our results suggested a novel mechanism of deactivating ZBTB14 and inducing its potential target genes in AD microglia.

From a computational perspective, *cEpiNets* provides several advantages for reconstructing context-dependent regulatory programs, particularly in low-abundance cell types such as microglia. First, by integrating curated bulk ATAC-seq datasets across perturbations, *cEpiNets* alleviates the sparsity and limited coverage of single-cell chromatin profiles (**Fig. 2a**), and it leverages TF footprinting to provide additional evidence of differential TF occupancy than other motif-based methods, such as *chromVar*^61^. Second, the framework translates differential TF occupancy into TF–CRE regulatory links and leverages graph embeddings that capture shared TF/cofactor regulatory profiles to predict CRE–gene relationships. Explicitly modeling CREs provides a critical intermediate layer for variant-to-function mapping, enabling non-coding disease-associated variants to be connected to their potential target genes and the *trans*-regulatory TF programs that may modulate their effects. Our approach complements conventional distance-, eQTL-, and correlation-based linking strategies^62–65^ by explicitly modeling the *trans*-regulatory TF programs connecting CREs and genes. Moreover, the learned embedding space could provide a flexible basis for extending CRE–gene prediction to new contexts lacking direct chromatin-interaction data, for example by reweighting embedding dimensions according to context-specific regulatory signals. Finally, given the limited generalizability of current perturbation-response prediction frameworks^66,67^, *cEpiNets* could provide a mechanistically informed prior that extends TF–gene models by explicitly incorporating the CREs through which regulatory effects are propagated.

Beyond linking disease-associated variants to target genes, cEpiNets revealed context-dependent *trans*-regulatory programs that may shape inflammatory responses in microglia. In particular, our cross-context analysis highlighted transcriptional repressors as potential modulators of inflammatory remodeling, with ZBTB14 emerging as a candidate regulator. The ZBTB14-associated network includes multiple genes implicated in neuroinflammation and AD, including *IL15*^59^, *ATF3*^68^, *SLC1A3*^69^, and *DNAJB6*^70^. Consistent with the network predictions, combined IFNγ/TNFα stimulation mimicking AD brain environment reduced *ZBTB14* expression while inducing the predicted target genes *IL15* and *ATF3*. In contrast, *SLC1A3* and *DNAJB6* did not respond under the same conditions, suggesting that individual branches of the ZBTB14 regulatory program are selectively engaged depending on the inflammatory context or additional regulatory inputs. Likewise, IFNγ alone was sufficient to induce *PD-L1* but did not alter *ZBTB14* or its downstream targets, indicating that repression of the ZBTB14 regulatory program is not a general consequence of inflammatory signaling but instead requires specific combinations or intensities of inflammatory stimuli that may associated with disease conditions. These observations are consistent with the central premise of *cEpiNets* that transcriptional regulatory programs are highly context dependent, with only subsets of predicted TF–CRE–gene interactions becoming active under particular biological conditions. Together with the inverse association between ZBTB14 footprint activity and AD neuropathology in human microglia, these findings support a model in which ZBTB14 participates in context-dependent regulation of inflammatory remodeling during AD progression.

To our knowledge, this is the first study to integrate TF footprinting and machine learning to provide functional epigenomic annotation of AD risk signals in microglia. Several limitations remain. First, many of the perturbation datasets were generated from human iPSC- or ESC-derived microglia, which may not fully recapitulate the regulatory complexity of human microglia *in vivo*. Second, our inflammatory perturbations primarily rely on IFN-*β* stimulation as a model of acute inflammation and therefore capture only a subset of the diverse inflammatory environments relevant to AD. Third, TF occupancy is inferred from ATAC-seq footprinting rather than directly measured by TF-specific assays such as ChIP-seq. Consequently, TFs with highly similar DNA-binding motifs, particularly members of the same TF family, can be difficult to distinguish. Finally, specific regulatory circuits identified by *cEpiNets*, including the ZBTB14 program, remain to be causally validated. Future studies incorporating TF-specific binding, perturbation, and chromatin-interaction data will help refine these networks and establish their regulatory mechanisms.

## Methods

### Datasets collection and ethical approval

The sample information of collected datasets were provided (**Table S1)**. The snATAC-seq data of human microglia was obtained from Morabito et al.^31^. The samples were isolated from the prefrontal cortex (PFC) of the postmortem human brain from individuals with late-stage AD (n = 12) and age-matched cognitively healthy controls (n = 8; 74–90+ years old). Other bulk ATAC-seq data of human iPSC- or ESC-derived microglia were from publicly available datasets under contexts of APOE isoforms^25^, AD risk gene mutants^26,27^, IFN-***β*** stimulation^71^, high-level lipid droplets^24^, co-cultured with cerebral organoid or xenotransplanted into mouse brain^29^, and HIV activated or HIV latent^30^. For the snRNA-seq and snATAC-seq datasets, the original studies were responsible for the ethical approval for sample collection.

### Preprocessing of bulk ATAC-seq datasets

We downloaded raw Fastq sequencing files and then assessed for quality, adapter content, and duplication rates with FastQC (V0.12.1) (https://www.bioinformatics.babraham.ac.uk/projects/fastqc/). We cut adapters and trimmed low quality reads with fastp (0.23.4) (--cut_mean_quality 20 -g -D --length_required 100)^72^, and aligned the clean reads to the human genome GRCh38-3.0.0 using Bowtie2 (V2.5.1)^73^ with default parameters. For each context, we merged clean reads of the biological replicate samples using the *view* and *merge* function of samtools (V1.6)^74^. We identified accessible regions through peak calling for merged alignments with MACS2 (V2.2.7.1)^75^ (--nomodel --shift 100 --extsize 200 --broad --qvalue 0.01 --broad-cutoff 0.01). We then merged peaks from the same dataset using the merge function of bedtools (V2.31.1)^76^, and defined them as CREs.

### Preprocessing of snATAC-seq datasets and cell type annotation

We downloaded the raw Fastq sequencing files for each sample and then mapped the reads onto the human reference genome (GRCh38) with cellranger-atac count (V2.1.0)^77^ to get the fragment files. We used ArchR (V1.0.2)^78^ to process the fragment files. We removed doublets for each sample using the *filterDoublets* function. The cells with transcription start site (TSS) enrichment > 6 and number of fragments between 1,000 and 100,000 were retained. We performed Iterative LSI dimension reduction and clustering based on a 500 bp tile matrix with parameters *iterations = 4, resolution = 0.2, varFeatures = 15,000*. UMAP was used to visualize cell embedding. We calculated gene scores with ArchR and annotated cell types based on marker genes from the original paper. After cell-type annotation, we conducted peak calling using the pseudobulk strategy of MACS2 (q-value < 0.01)^75^. The sequencing reads of microglia in each individual were extracted with the *filterbarcodes* function of sinto (V0.10.0) (https://github.com/timoast/sinto), and with the microglia peaks together were used for the following footprinting. We finally merged microglial snATAC-seq peaks and those from bulk ATAC-seq together into a set of union peaks with bedtools.

### Transcription factor (TF) footprinting and differential binding

We predicted TF binding score for each TF under each context using TOBIAS pipeline (V0.13.3)^33^. The human genome GRCh38 was used as a reference genome and a total of 878 TF motifs were downloaded from JASPAR CORE 2024^79^. Briefly, we utilized the TOBIAS ATACorrect tool to correct an inherent insertion bias from Tn5 transposase of ATAC-seq data. Then we performed TOBIAS ScoreBigwig to calculate the footprint score at each binding site across accessible regions, defined by the union peak set above. The footprint scores were integrated with the information of TF binding motifs to predict specific TF binding across the genome using TOBIAS BINDetect. We also used TOBIAS BINDetect to calculate differential binding of TFs between the control and treated groups for each context pair. According to TOBIAS, the TFs with top 5% -log10(p-value) or top 5% differential binding scores in either direction were identified as differential footprinting TFs (dfTFs). We confirmed that all the dfTFs were expressed with matched RNA-seq or snRNA-seq datasets.

### Predicting (differential) CRE activity based on TF footprint scores and identification of differential-footprinting CREs (dfCREs)

For each context pair, we calculated the number of alignments from Bam files with *bedtools multicov* and reads per million mapped reads (RPM) for each peak of the union set to measure observed CRE activity. Log2 fold change of RPM between control and treated groups was calculated to represent observed differential CRE activity. We used a linear regression model and a random forest model to predict CRE activity and differential CRE activity based on TF footprint score, respectively, with the function of *lm* in R (V4.4.0) and an R package *ranger*^80^. For the prediction in each context pair, we used 70% of all the CREs or dfCREs as training dataset and the remaining 30% as testing dataset. For each dfTF, we defined its target dfCREs from the union peak set according to the direction and magnitude of differential footprinting. For upregulated dfTFs, we selected CREs with differential footprint scores in the top 10% among positive values; for downregulated dfTFs, we selected CREs in the bottom 10% among negative values.

### Graph embedding based on dfTF-dfCRE linking

We projected TFs and CREs into a shared embedding space using node2vec^35^ implemented in PecanPy (SparseOTF mode)^81^. In this framework, each node is represented by a numeric vector, and distances between vectors reflect regulatory proximity.

We first generated protein embeddings from the STRING protein–protein interaction (PPI) network^82^ (walk_length = 5), and computed pairwise Euclidean distances between TFs. For each context, we then integrated the TF–TF PPI network with context-specific dfTF–dfCRE links to construct a weighted graph capturing both protein interactions and regulatory relationships. Only TF–TF edges involving TFs present in the dfTF–dfCRE network were retained. All dfTF–dfCRE edges were assigned weight 1. For each TF_i_, we defined TF_i_–TF_j_ edge weights as e^-20^ ^×^ ^distanceᵢ,ⱼ^ and scaled relative to the total TF–CRE edge weights using a parameter (*TF_TRANS_PROB_RATIO_TF_vs_CRE*) that controls the probability of random walks transitioning from a TF to another TF versus to a CRE (set to 1 in this study). Then we generated random walks for each context (num_walks = 10, walk_length = 10) based on each weighted network, and finally learned node embeddings from the aggregated walks from all contexts (vector_size = 128, window = 5, min_count = 1, sg = 1, workers = 4).

To test whether embedding learned could be used to predict dfTF-dfCRE linking in unseen context, we used a leave-one context-out cross-validation method. We treated dfTF-dfCREs of the target context as true signals each time and used embedding learned from the rest of the contexts as features to predict the true signals. In each target context, we split dfTF-dfCREs into two separate datasets based on upregulation or downregulation. For each dfTF, we treated the dfTF and its target dfCREs as positive TF–CRE pairs. We generated an equal number of negative pairs by randomly sampling CREs from the bottom 50% of the CRE sets ranked by differential footprint score in the direction corresponding to that dfTF. Then, for each TF-CRE pair, we calculated the difference vector in the embedded space between TF and CRE as features. We used 70% of pairs as the training dataset and the remaining 30% as the testing dataset to predict true dfTF-dfCREs with logistic regression (*glm* function in R). We also tested whether embedding from a single context had equal power compared to embeddings from all the rest of contexts to predict dfTF-dfCREs in the same target context. We sampled to generate the training and testing sets ten times for each target context and compared results with AUPRCs. We shuffled the remaining embedded vectors randomly and conducted the same prediction process for comparison.

### CRE-gene linking prediction

To dissect and compare epigenomic regulatory networks in AD microglia with those from non-AD contexts, we trained embeddings with only dfTF-CREs from non-AD contexts. We next mapped CREs overlapping gene transcription start sites (TSSs) as promoters and considered CREs within 1 Mb of a TSS as candidate enhancers for the corresponding gene. For each candidate CRE–gene pair, we calculated the difference between the embedding vectors of the candidate enhancer and promoter CRE and used these differences as features in a logistic regression model to predict CRE–gene regulatory links. Training labels were derived from ABC scores^32^ reported in the original microglial Hi-C study^11^: CRE–gene pairs with the top 5% of ABC scores were defined as positive links, whereas an equal number of negative links were randomly sampled from the bottom 50% of ABC scores and matched to positive links by genomic-distance bins. To avoid ambiguity in promoter assignment, model training and evaluation were restricted to genes whose TSS mapped uniquely to a single CRE. Predictive performance was evaluated using leave-one-chromosome-out cross-validation. Within each held-out test set, we benchmarked the embedding-based score against alternative measures of CRE–gene association, including genomic distance, marginal eQTL association P values, posterior inclusion probabilities (PIPs) from eQTL fine-mapping^37^, and co-accessibility scores derived from ATAC-seq data^8^. Finally, we trained the logistic regression model using data from all chromosomes and applied it to candidate enhancer–gene pairs to generate predicted CRE–gene regulatory scores based on their embedding-vector differences.

We finally used the CRE-gene links derived from embedding-based prediction, meeting both criteria: (1) with top 10% of predicted probability and (2) allowing only top 3 predicted target genes per CRE. We additionally combined predicted CRE-genes by linking each CRE to its nearest gene within 100 kb upstream of the TSS or downstream of the transcription terminal site (TTS) using UROPA^83^ (--feature_anchor start end --distance 100000 100000). All the predicted links from both embedding- and nearest gene-based methods were used for the following analyses.

### dfCRE modules

To identify the modules of dfCRE based on their regulation of TFs across contexts, we first generated a binary matrix representing context-dependent regulation with dfCREs in rows and dfTF-context combinations in each column. We assigned “1” to the cells where dfTF-CRE pairs were present under the specific context, and we assigned “0” to the rest of the matrix. We then clustered dfCREs into 100 modules by the R package flexclust^84^ with a k-centroids algorithm (k=100, family=kccaFamily("ejaccard")) following a previous report^65^. We annotated the genomic location and biological function of dfCRE modules based on the biological processes of gene ontology enrichment for nearby genes using the R package rGREAT^85^. For each module, we showed statistics on the number of dfCREs and the proportion of dfCREs located within 5kb from TSS based on genomic annotation. The heatmap of dfCRE modules and the following correlation analyses were visualized by the R package ComplexHeatmap^86^.

### dfCRE modules enriched for AD differential signals from another AD snATAC-seq dataset

We compared our dfCREs with AD-associated differential signals from a previous brain snATAC-seq report by Xiong *et al.*^10^ The snATAC-seq peak matrix was obtained from the ArchR object, and we performed FindAllMarkers to quantify differential signals of all CREs between AD (n = 44) and control individuals (n = 48) in PFC (min.pct = 0.1, logfc.threshold = 0, and test.use = MAST). We controlled the covariates of age, sex, postmortem interval, and number of fragments from the original report. The generated average log2FC was used as CRE ranking, and our dfCREs from each module was used as CRE sets for GSEA^41^ enrichment with the fgsea R package^87^.

### dfCRE enrichment of TF EHF and Fos:Jun cobinder in dfCRE modules

We calculated the number of bound dfCREs by EHF, FOS:JUN, and their overlaps in AD and *SORL1* KO contexts, separately. We performed Fisher’s exact test (two-sided) with fisher.test in R to test the enrichment of bound dfCREs across dfCRE modules. For testing the enrichment of co-bound dfCREs by EHF and FOS:JUN, dfCREs of each module were used as background, and the odds ratio and FDR (using Benjamini-Hochberg, BH procedure) were calculated.

### dfTF-CRE-gene level enrichment in marker genes of microglia states

We obtained the Seurat object of microglia snRNA-seq data reported by Sun *et al*.^8^ The “RNA” assay of the object was used to identify marker genes for each microglial state (MG0-12), with the FindAllMarkers function (min.pct = 0.1, logfc.threshold = 0, return.thresh = 1, test.use = MAST). The logFC of gene expression of cells in each microglial state against the rest cells was used as the ranking for GSEA enrichment analysis. For each context, target genes linked to each TF (e.g, TF-gene regulons) through TF-CRE-gene linking by *cEpiNets* were used as gene sets in GSEA analysis for each microglial state. We defined TFs with a significantly enriched regulon (GSEA adjusted *P* < 0.05) as driver TFs, further filtered target genes based on leading-edge analysis from fGSEA, and finally presented the whole TF-CRE-gene links as potential driving circuits for a specific microglia state. We visualized the circuits with the igraph R package^88^. We also obtained the Seurat object of Human Microglia Atlas (HuMicA) and conducted the same GSEA enrichment test for each cluster.

### Correlation between *SORL1* risk variants and MG8 cell proportion

We obtained the MG8 cell proportion of individuals from the Seurat object reported by Sun *et al*.^8^, and tested its association with each AD GWAS variant (*P* < 1✕10^-5^) located at CREs linked to *SORL1* with a linear regression model considering age, sex, and AD status in the original report. We carried out multiple test corrections with the BH procedure, and used the adjusted *P* < 0.05 as the cutoff.

### *Trans*-regulations at *SORL1* locus

We utilized networks of dfTF-dfCRE linked to *SORL1* across contexts to annotate AD GWAS signals^44^. The track browser for visualizing was generated from IGV^89^.

### Correlation between inflammatory repressors, AD dfTFs, and AD pathological/cognitive scores

To evaluate whether dfCREs bound by TFs under inflammatory conditions were preferentially located near dfCREs bound by AD-associated dfTFs, we performed a permutation-based enrichment analysis. For each AD dfTF, we randomly sampled matched numbers of dfCREs from the full set of dfCREs (30 permutations) and quantified the number of sampled elements located within 10 kb of dfCREs targeted by inflammation-associated repressors. This procedure generated a null distribution of expected overlap counts, from which we estimated the mean and variance. Observed overlap counts were then compared against this null distribution to assess enrichment. *P* values were adjusted for multiple testing using the BH procedure.

To assess donor-level associations, we calculated, for each individual, the mean footprint scores of dfCREs targeted by each repressor under the specified inflammatory conditions, as well as those targeted by AD-associated dfTFs, using the microglial snATAC-seq data from 238 individuals^10^ processed with ArchR. Footprint scores were adjusted for sequencing depth by regressing out ReadsInTSS. We then computed correlations between residualized footprint scores and AD neuropathological or cognitive measures in the original report. Corresponding *P* values were corrected for multiple testing using the BH procedure.

We obtained the same dfCREs targeted by each repressor and dfTFs in AD microglia and predicted target genes. These dfCREs were mapped to AD GWAS signals^44^. The networks of dfCREs harboring AD GWAS signals (*P* < 1✕10^-5^), linked dfTFs, and target genes were visualized with the igraph R package^88^. For the graph presentation, we combined dfTFs from the same family together in the network. Details of dfTFs-dfCREs-target genes can be found in **Data S8**.

### Prioritizatizing candidate targets of ZBTB14

To prioritize ZBTB14 target genes for experimental validation, we integrated three lines of evidence: (1) the genes predicted as ZBTB14 targets by *cEpiNets*; (2) their expression across donors was negatively associated with ZBTB14 footprint scores defined above; and (3) positively associated with early- or late-AD status across donors with *zlm* function in the MAST^90^, adjusting for age, sex, batch, brain region, mitochondrial and ribosomal gene proportions, and postmortem interval; Finally, we examined their expression changes following IFN-β stimulation using data from Yang *et al.*^24^ as an additional support for inflammation responsiveness. We classified those candidate genes into two tiers based on whether or not direct literature evidence supports their function in AD and showed the results in **Data S9**.

### Cell culture

The human microglial clone 3 cell line (HMC3^91,92^, ATCC, CRL-3304) was cultured in DMEM, high glucose (Gibco, 11-965-092) supplemented with 1% Antibiotic-Antimycotic (Gibco, 15-240-062) and 2% fetal bovine serum (Gibco, 10-438-026). The HMC3 were plated for experiment 24 hours prior to cytokine treatment at the density of 30,000 cells per cm^2^. At the start of the experiment, the media was refreshed with control media, or media supplemented with human IFNγ (100 ng/mL) and human TNFα (50 ng/mL). The cells were captured with Evos M7000 microscope (Thermofisher) on a 10x objective. After 6 hours of cytokine treatment, the cells were washed with DPBS and imaged.

### Reverse transcription quantitative PCR

After imaging of cells, the cells were collected for RNA isolation with RNeasy Mini kit (Qiagen) and reverse transcribed by cDNA synthesis (iScript. Bio-rad) using the MiniAmp Thermal cycler (ThermoFisher). The relative gene expression of selected genes was determined with ViiA7 RT-qPCR instrument (ThermoFisher) using SYBR green (Fisher Scientific) reagent and beta-2-microglobulin (B2M) as a reference gene. Total 10 biological replicates were generated with two technical replicates for RT-qPCR. 0.5 ng/μL cDNA was loaded per reaction. Reagents and resources are provided (**Table S3).**

## Data availability

The *cEpiNets* object is available in the web interface (https://cepinets.thresh-infra.xyz/). The accession codes and/or links for the original bulk ATAC-seq and snATAC-seq datasets used to generate the *cEpiNets* are as follows: Yang *et al.* (GSE173316); Rheinberger *et al.* (GSE205807); Liu *et al.* (GSE153658); Zhao *et al.* (GSE263804); Murphy *et al.* (GSE271384); Haney *et al.* (GSE254205); Han *et al.* (GSE226690); Morabito *et al.* (GSE174367). The snATAC-seq data used for mapping AD epigenomic signals and the AD neuropathological or cognitive measures are from the report by Xiong *et al.* (https://www.synapse.org/#!Synapse:syn52293417). The AD snRNA-seq data used for analyses of microglial state enrichment and inflammation associated *SORL1* variants identification is from the report by Sun *et al.* (https://www.synapse.org/#!Synapse:syn52293417).

## Code availability

All codes for this study were deposited in Github (https://github.com/HouGroup-CompBio-GxE/cEpiNets).

## Acknowledgements

The authors would like to thank all the suggestions from Drs. Matheus Victor, Pawel Przytycki, and Jubao Duan. The project was supported by the NIH grants U19-AG068753 to L.F and L.H. and R01-AG083941, R01-AG082362 to J.TCW.

## Author Contributions Statement

L.H. and T.F conceived the study. T.F carried out the computational analysis under the direction of L.H, with the help from N.S and J.Z. M.K. performed experimental validation under the supervision of J.TCW. Jiayong.T built the database and website with the help from T.F. T.F. and J.TCW. wrote the manuscript with input from L.H, M.K, N.S., and L.F.

