## Supplementary Figures and Tables for "Context-dependent regulatory networks connect Alzheimer’s disease genetics to microglial inflammatory responses"

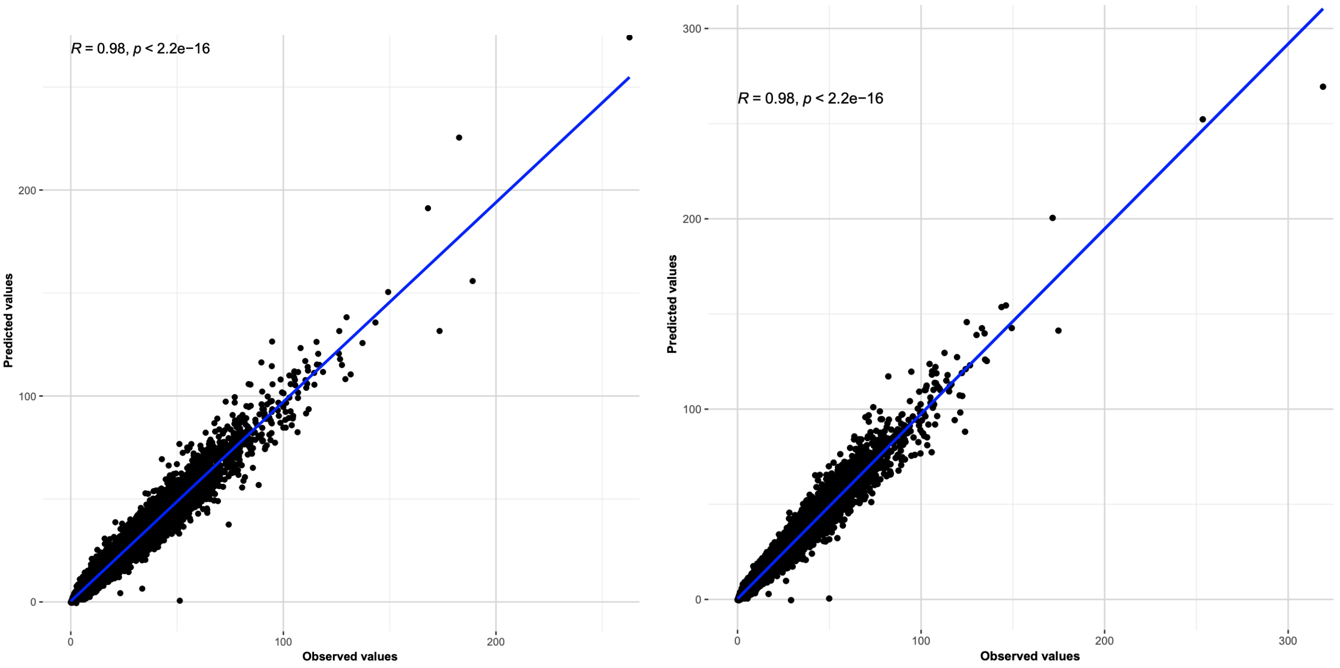


**Fig S1. Multiple linear regression between predicted (x axis) and observed (y axis) CRE activity in the AD context pair.** The left panel is the control group and the right panel is the AD group. Each dot is a CRE. The blue line is the fitted curve.

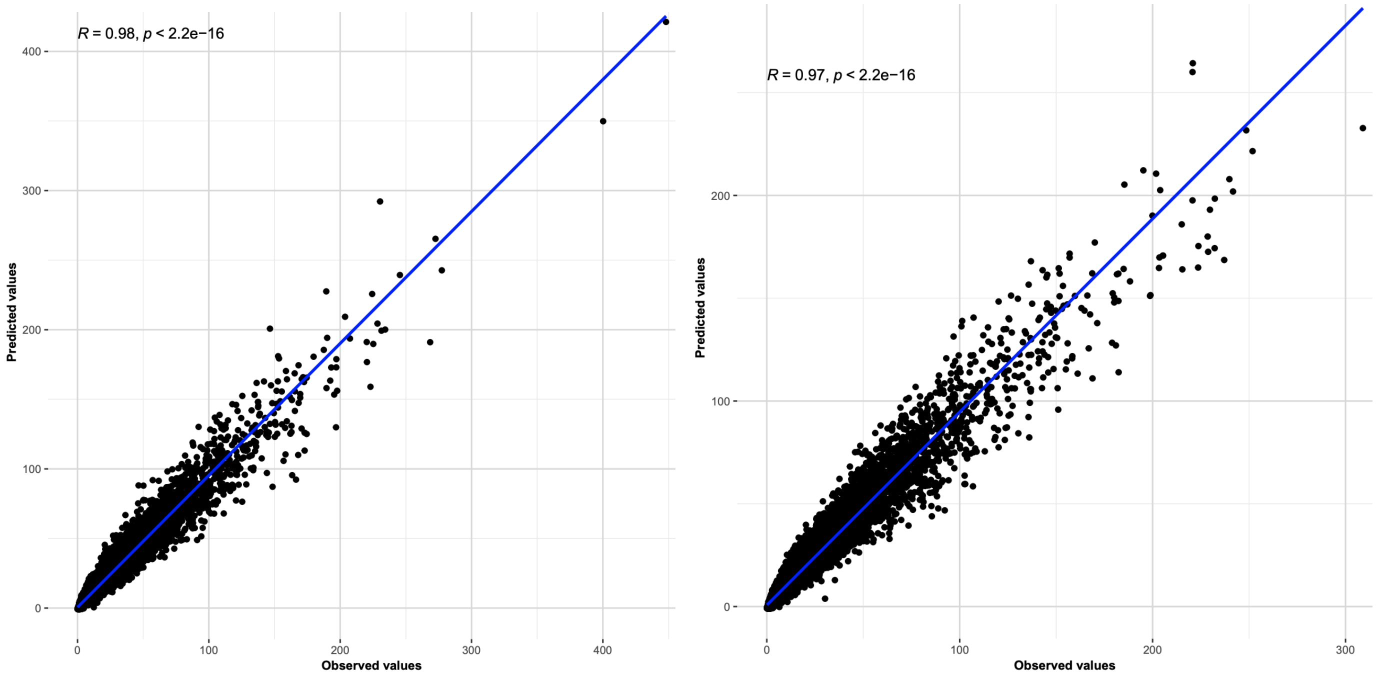


**Fig S2. Multiple linear regression between predicted (x-axis) and observed (y-axis) CRE activity in the High-lipid droplets (LD) context pair.** The left panel is the LD-low group and the right panel is the LD-high group. Each dot is a CRE. The blue line is the fitted curve.

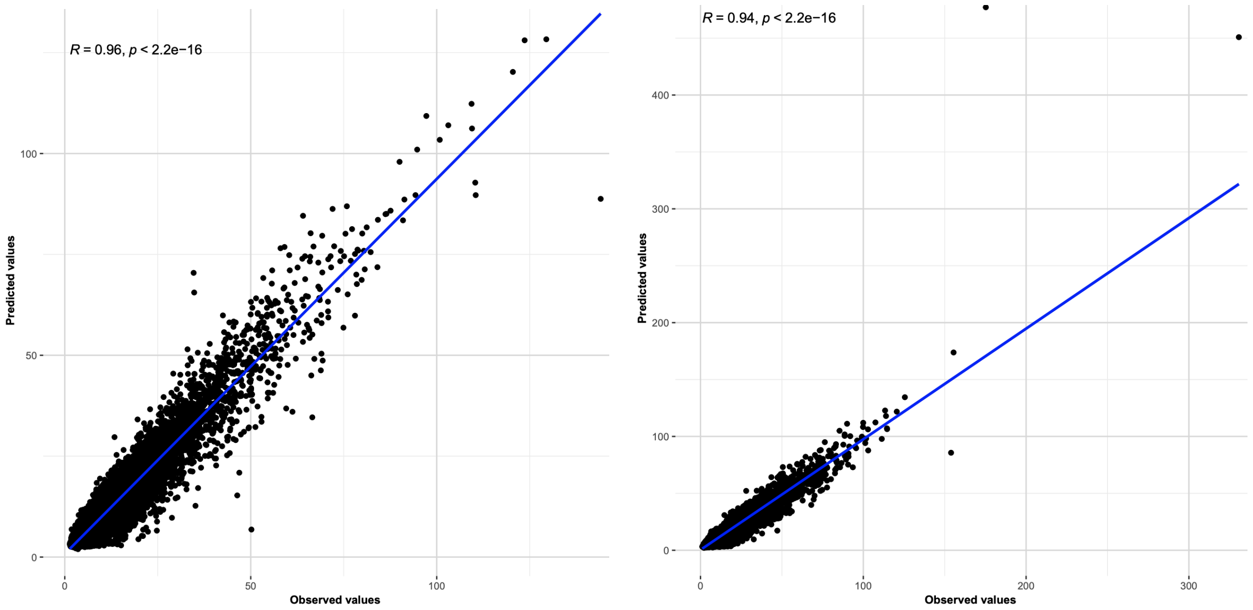


**Fig S3. Multiple linear regression between predicted (x-axis) and observed (y-axis) CRE activity in the *CD33* variant context pair.** The left panel is the wild-type group and the right panel is the *CD33*-mutant group. Each dot is a CRE. The blue line is the fitted curve.

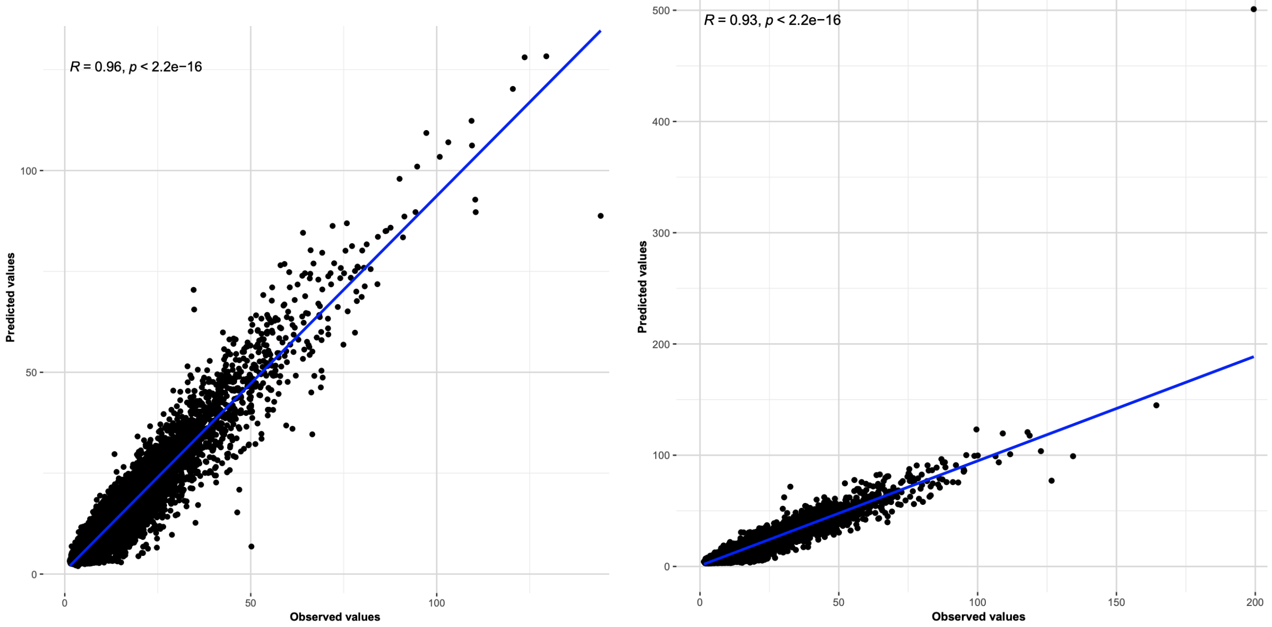


**Fig S4. Multiple linear regression between predicted (x-axis) and observed (y-axis) CRE activity in the *INPP5D* variant context pairs.** The left panel is the wild-type group and the right panel is the *INPP5D*-mutant group. Each dot is a CRE. The blue line is the fitted curve.

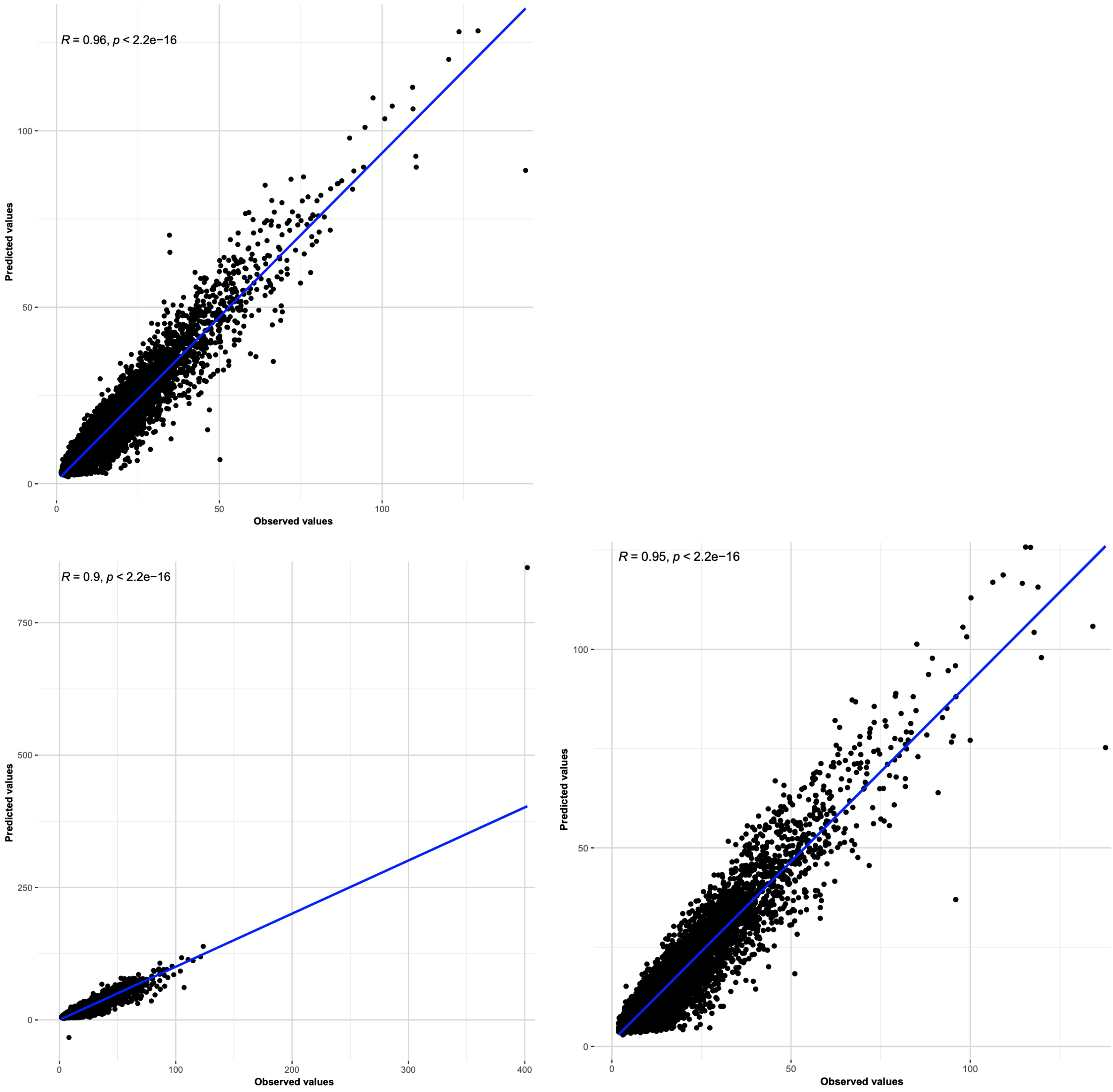


**Fig S5. Multiple linear regression between predicted (x-axis) and observed (y-axis) CRE activity in *SORL1* variant or *SORL1* KO context pairs.** The top-left panel is the wild-type group and the bottom panels are the *SORL1A528T* and *SORL1* KO groups separately. Each dot is a CRE. The blue line is the fitted curve.

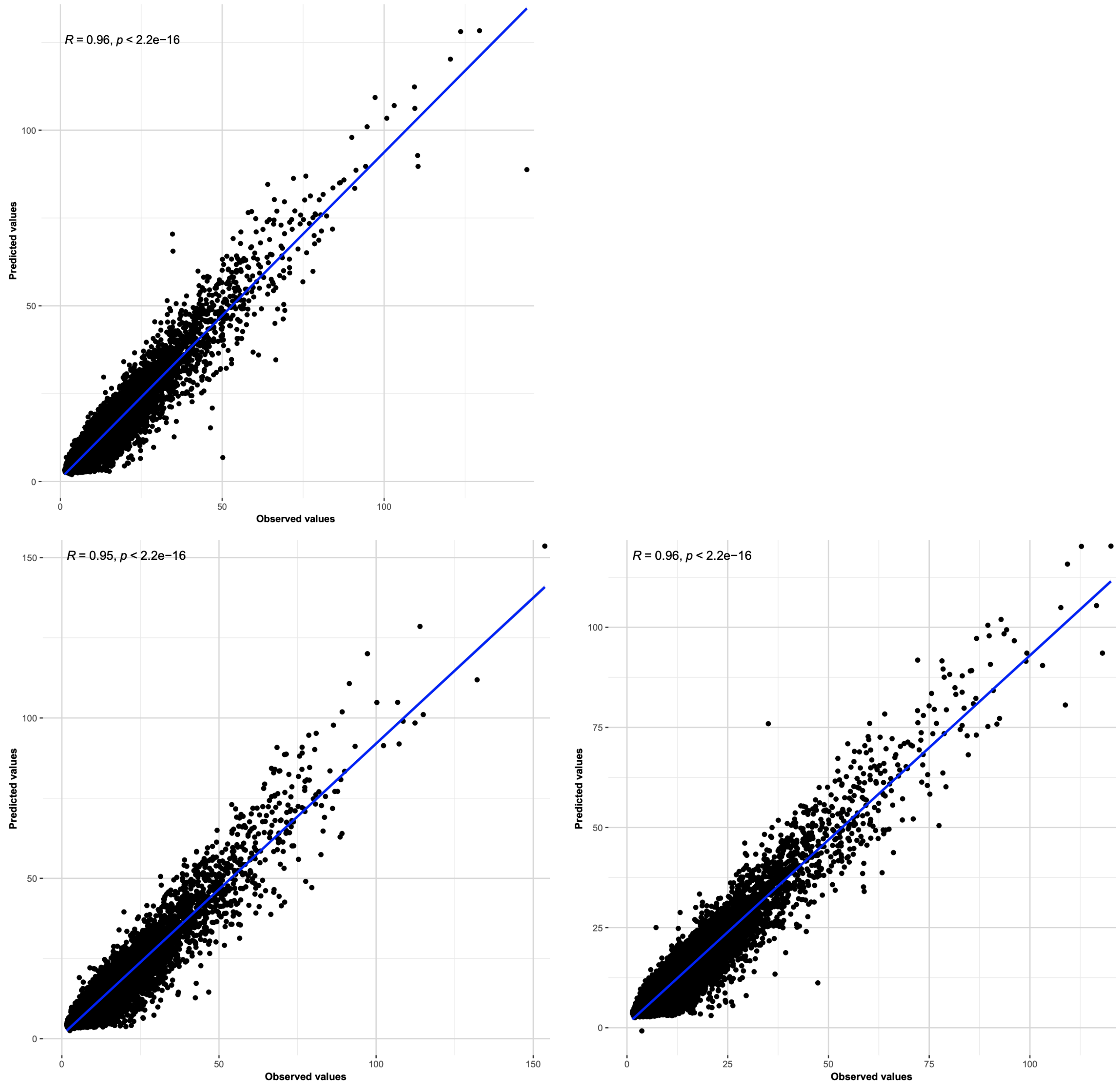


**Fig S6. Multiple linear regression between predicted (x-axis) and observed (y-axis) CRE activity in *TREM2* variant or *TREM2* KO context pairs.** The top-left panel is the wild-type group and the bottom panels are the *TREM2R47H* and *TREM2* KO groups separately. Each dot is a CRE. The blue line is the fitted curve.

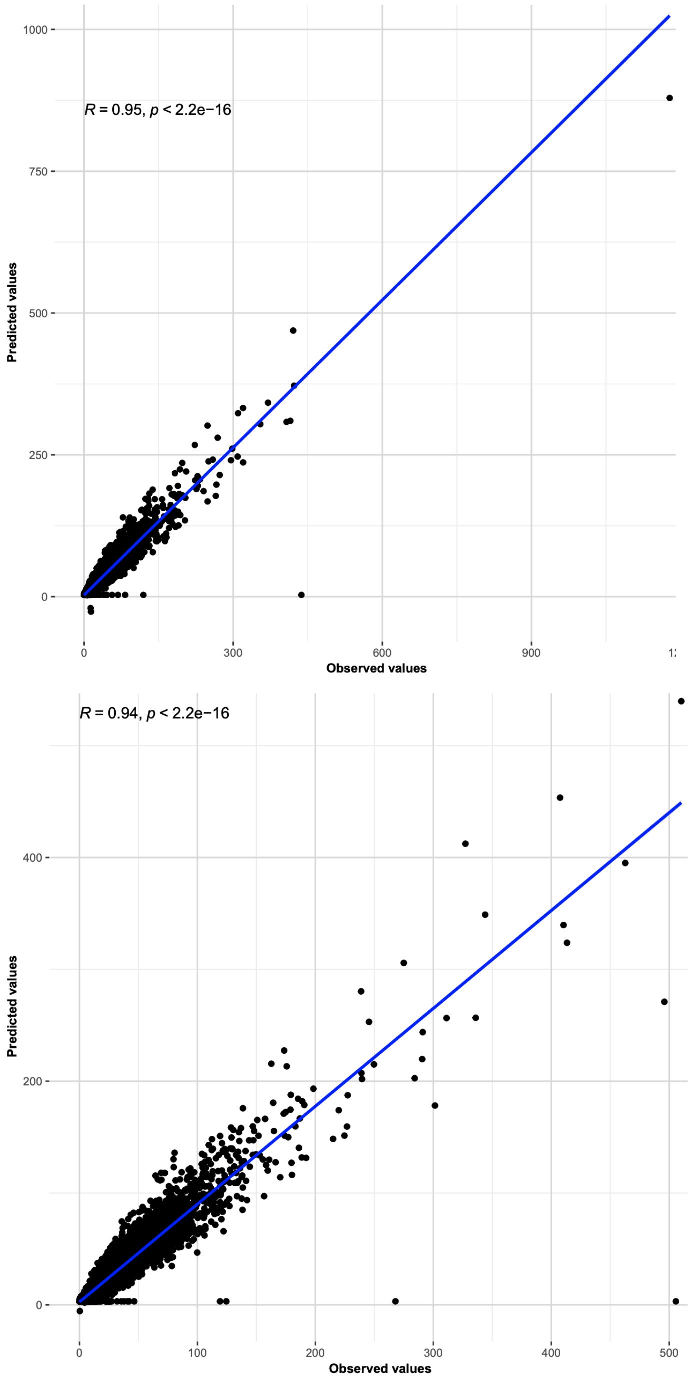


**Fig S7. Multiple linear regression between predicted (x-axis) and observed (y-axis) CRE activity in APOE isoform context pairs.** The top-left panel is the *APOE3* (control) group and the top-right one is the *APOE2* group. The bottom-left panel is the *APOE4* group and the bottom-right panel is the *APOE* KO group. Each dot is a CRE. The blue line is the fitted curve.

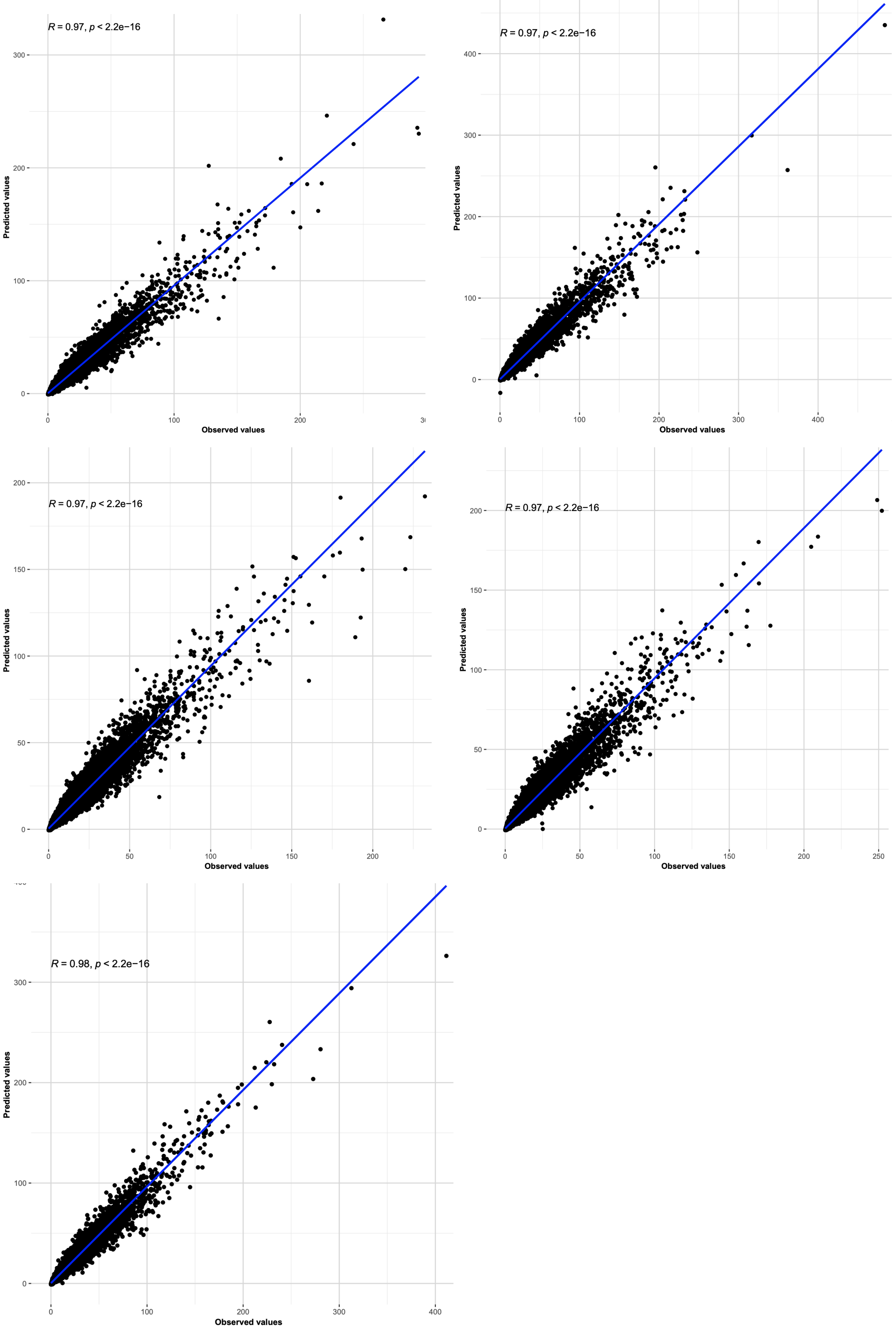


**Fig S8. Multiple linear regression between predicted (x-axis) and observed (y-axis) CRE activity in co-culture or xenotransplantation context pairs.** In the order from top-left to bottom-left, the figure shows iPSC-derived microglia, co-culture with organoids, 7-day post-xenotransplantation, 12-day post-xenotransplantation, and 8-week post-xenotransplantation. Each dot is a CRE. The blue line is the fitted curve.

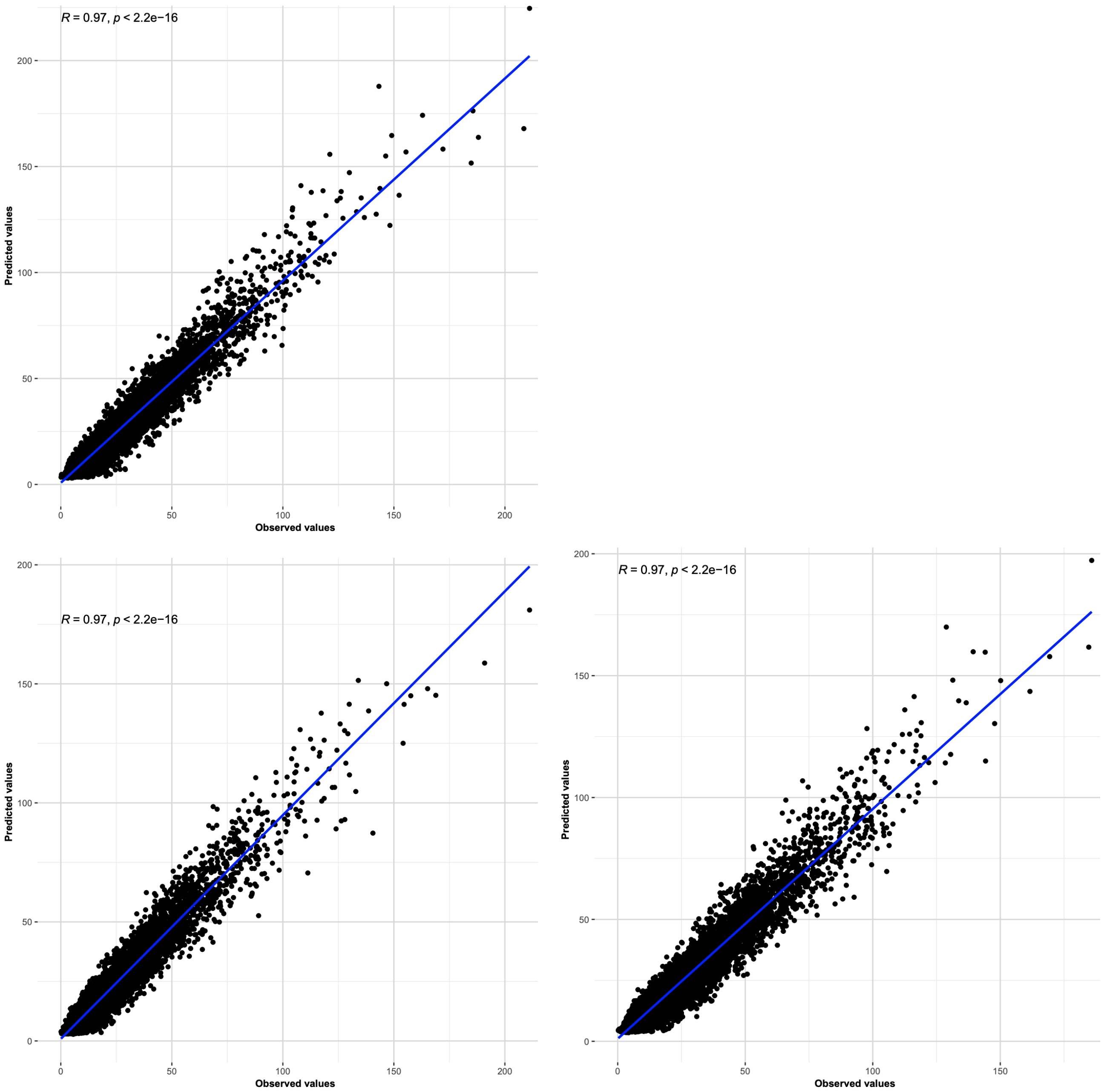


**Fig S9. Multiple linear regression between predicted (x-axis) and observed (y-axis) CRE activity in HIV infected context pairs.** The top left panel is the HIV uninfected (control) group. The bottom left panel is the HIV latent group and the bottom right is the HIV activated group. Each dot is a CRE. The blue line is the fitted curve.

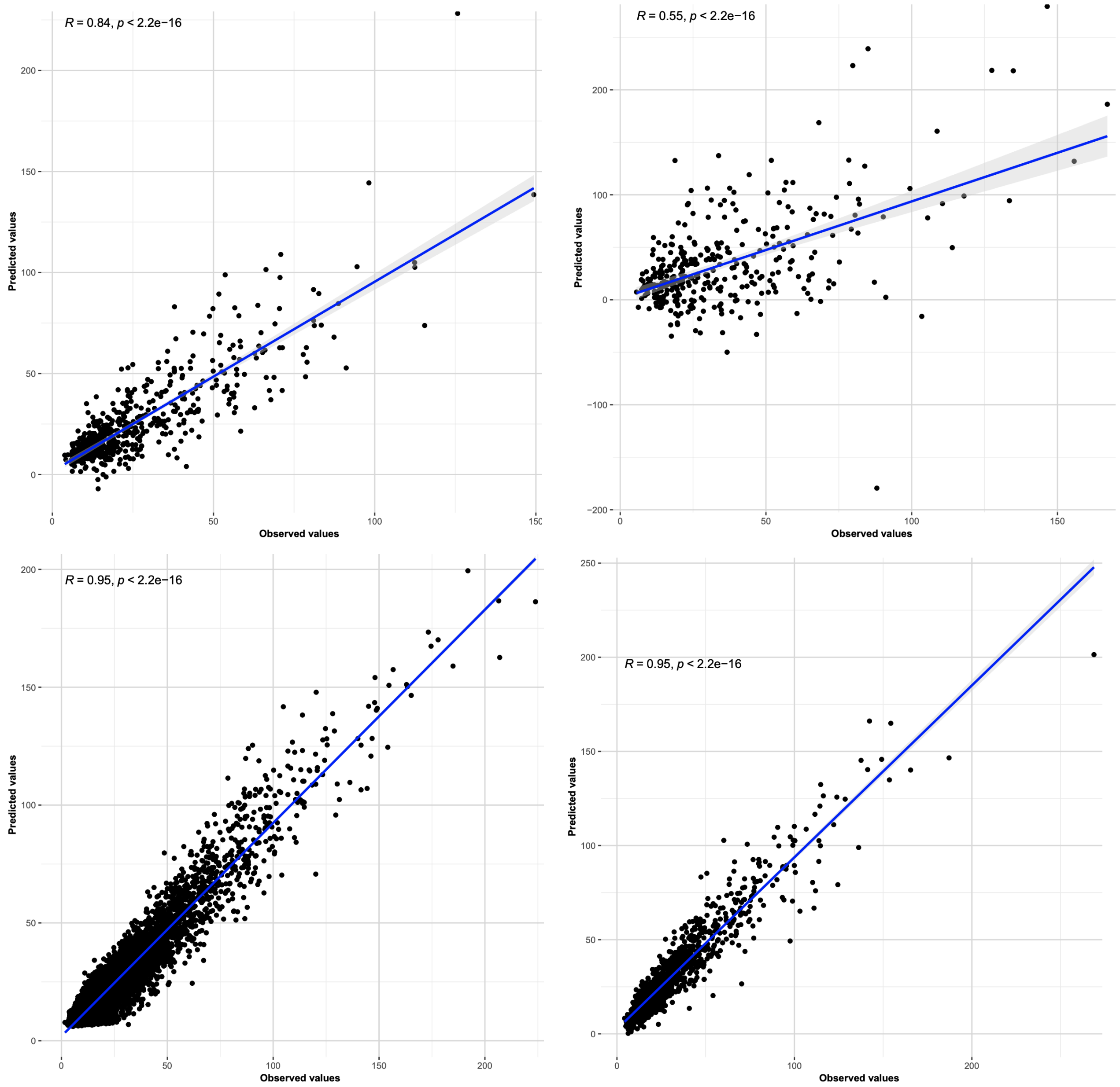


**Fig S10. Multiple linear regression between predicted (x-axis) and observed (y-axis) CRE activity in H1 (top) or WTC11 cell line (bottom) under IFN-*β* context pairs.** The top panels are the H1 (left) and H1-IFN-*β* (right) groups. The bottom-left panels are the WTC11 and WTC11-IFN-*β* group. Each dot is a CRE. The blue line is the fitted curve.

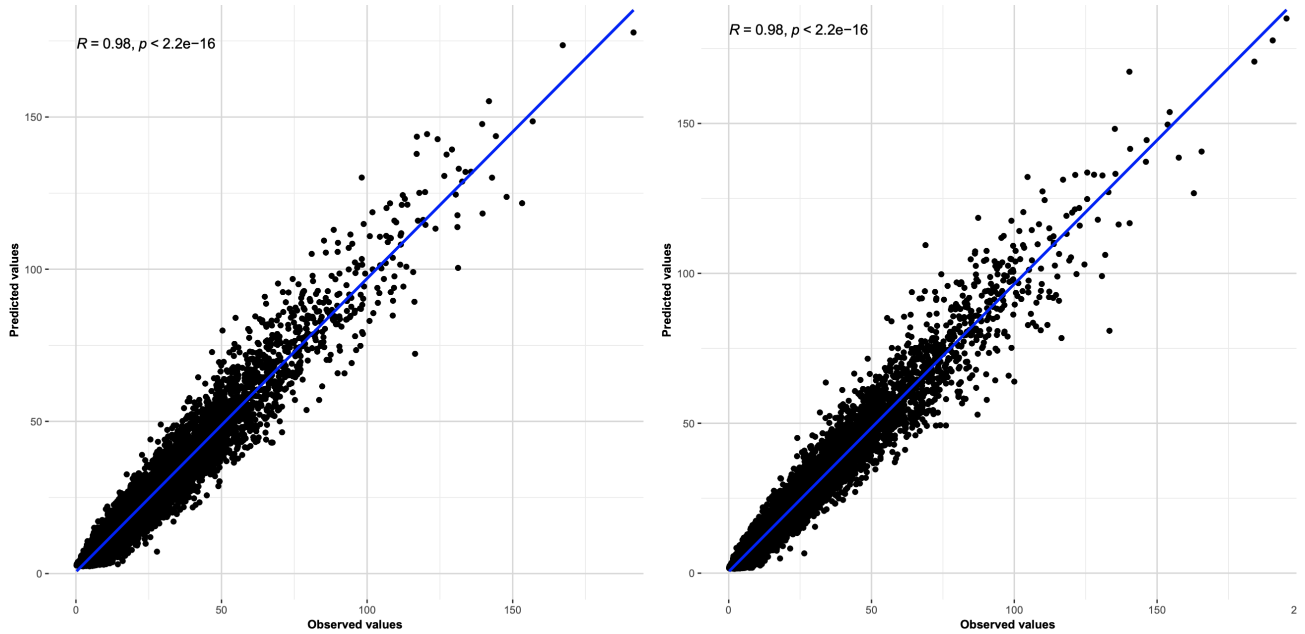


**Fig S11. Multiple linear regression between predicted (x-axis) and observed (y-axis) CRE activity in the CRISPR-edited *CLU* context pair.** The left panel is the control group and the right panel is the *CLU*-mutant (CRISPR-Cas9-edited) group. Each dot is a CRE. The blue line is the fitted curve.

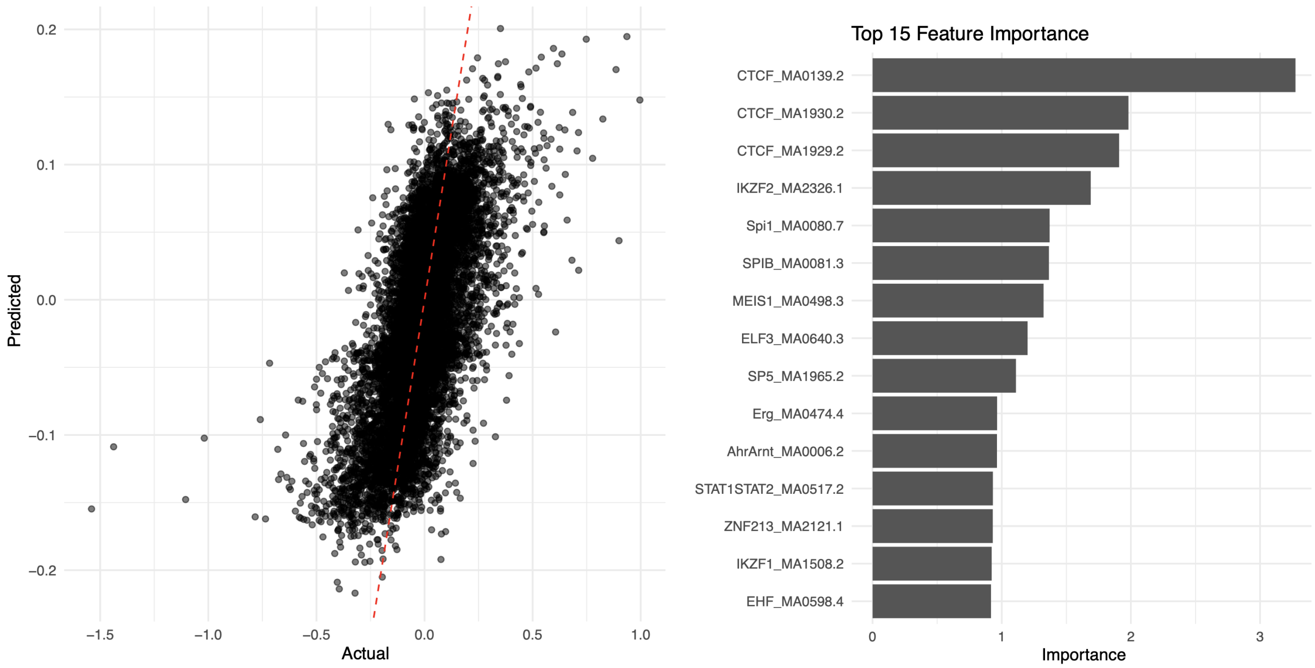


**Fig S12. Random forest model for prediction of dfCRE activity in AD context pair (left) and top 15 TFs important in prediction (right).** In the left panel, the x-axis is the actual dfCRE activity value and the y-axis is the predicted value. Each dot is a dfCRE. The red dashed line is the fitted curve. In the right panel, the x-axis is the importance from the random forest model and each row represents a TF.

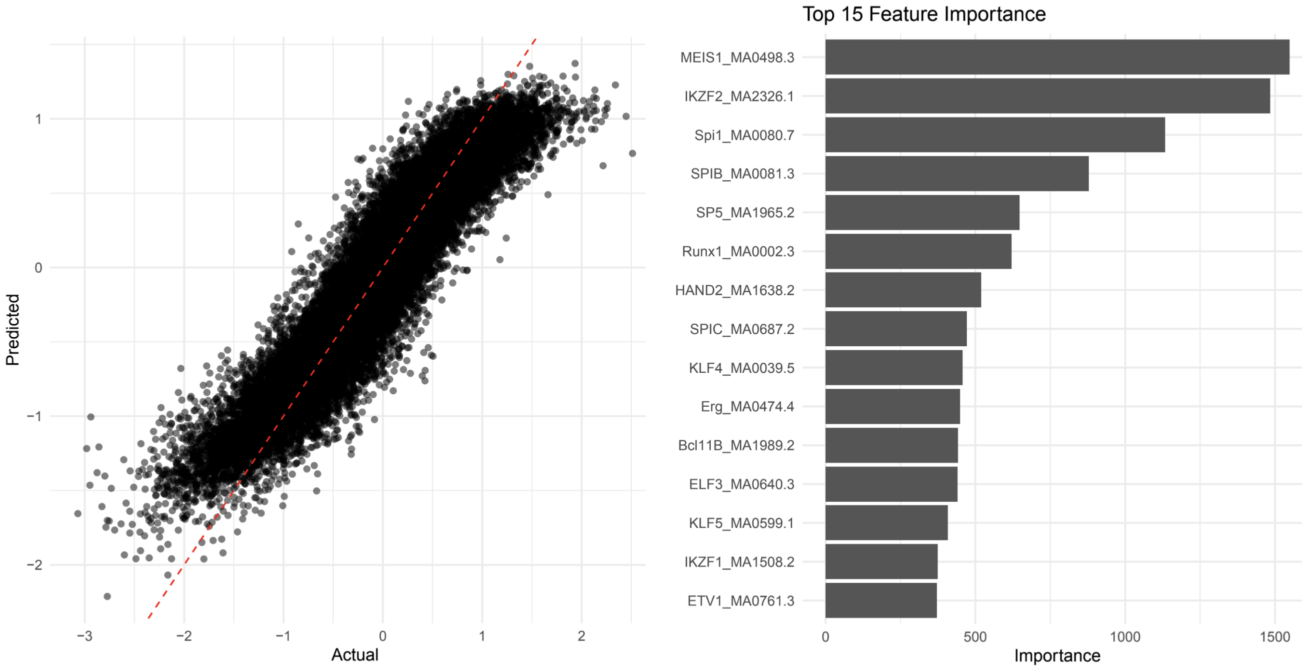


**Fig S13. Random forest model for prediction of dfCRE activity in high-LD context pair (left) and top 15 TFs important in prediction (right).** In the left panel, the x-axis is the actual dfCRE activity value and the y-axis is the predicted value. Each dot is a dfCRE. The red dashed line is the fitted curve. In the right panel, the x-axis is the importance from the random forest model and each row represents a

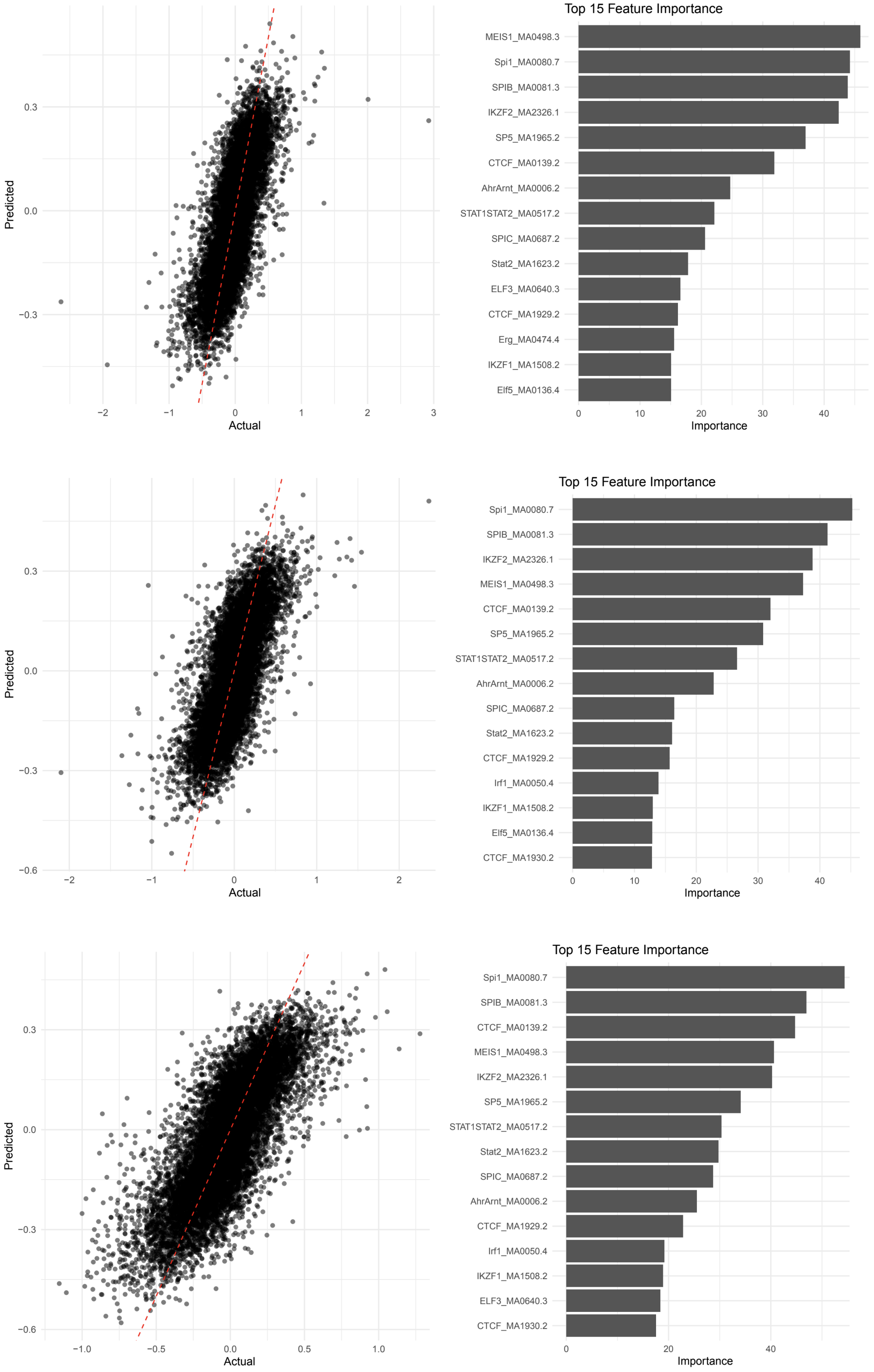


**Fig S14. Random forest model for prediction of dfCRE activity and top 15 important TFs in context pair of *APOE4* vs *APOE2* (above), *APOE4* vs *APOE3* (middle), or *APOE* KO (below).** For each row on the left, the x-axis is the actual dfCRE activity value and the y-axis is the predicted value.

Each dot is a dfCRE. The red dashed line is the fitted curve. In the right panels of each row, the x-axis is the importance from the random forest model and each row represents a TF.

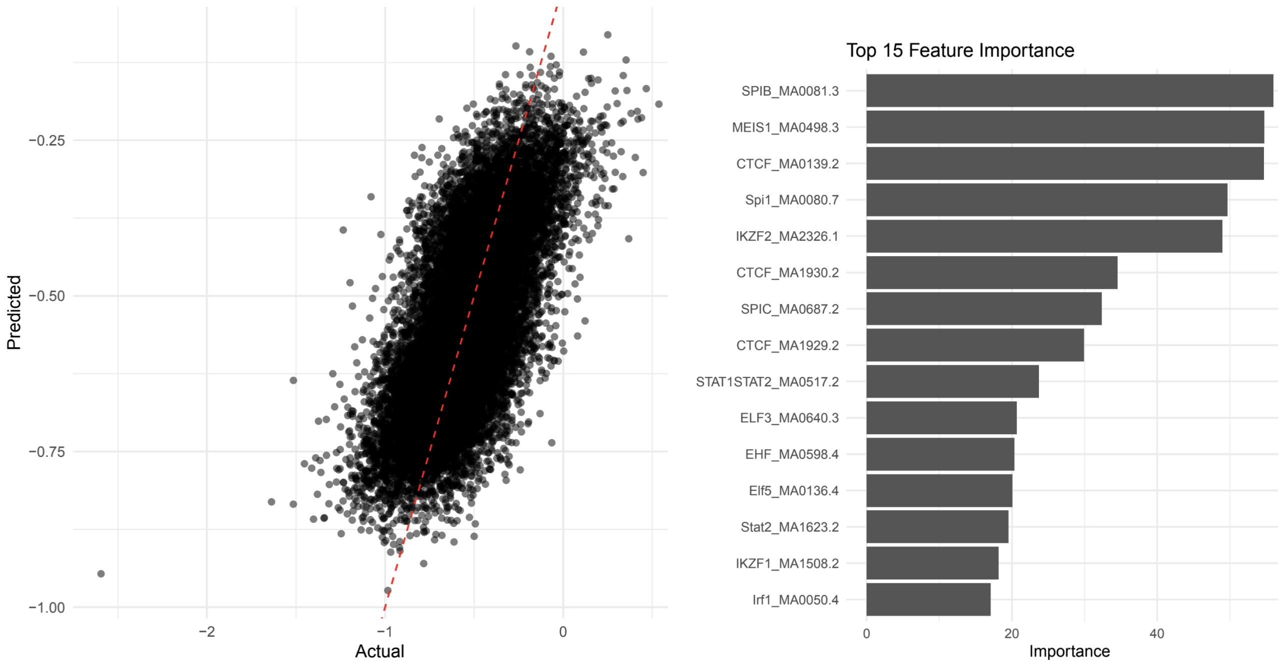


**Fig S15. Random forest model for prediction of dfCRE activity in *CD33* with GWAS variant context pair (left) and top 15 TFs important in prediction (right).** In the left panel, the x axis is the actual dfCRE activity value and y axis is the predicted value. Each dot is a dfCRE. The red dashed line is the fitted curve. In the right panel, the axis is the importance from random forest model and y axis is a TF.

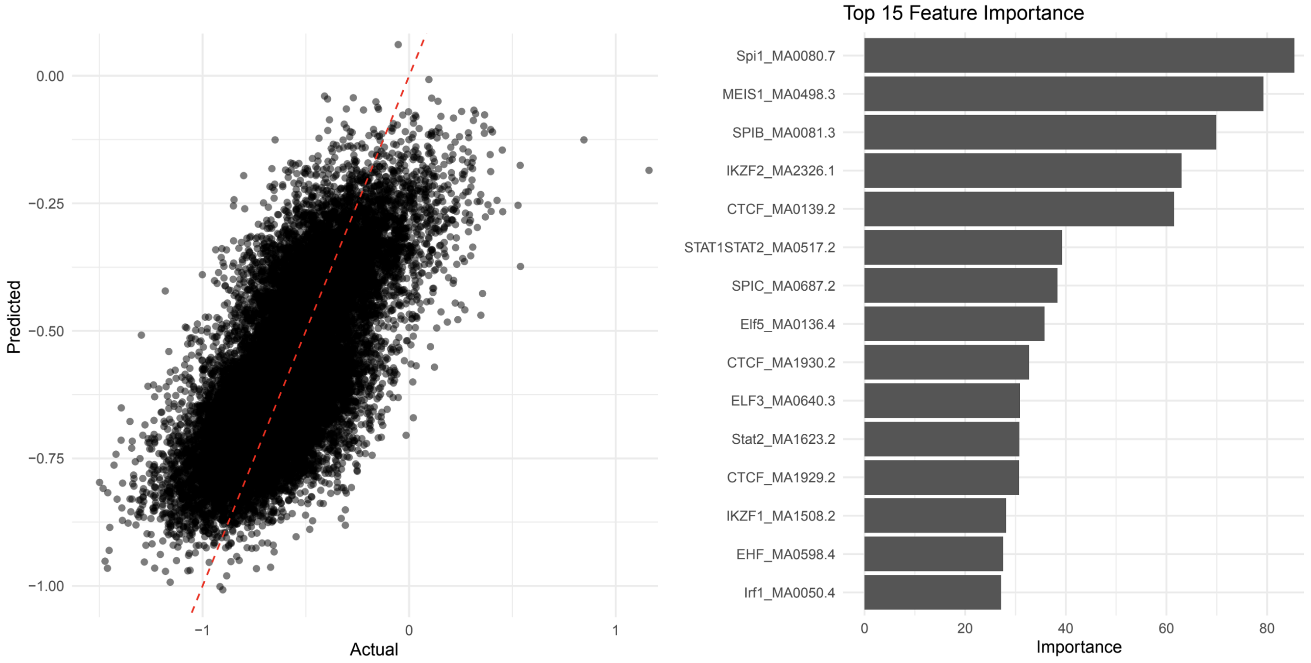


**Fig S16. Random forest model for prediction of dfCRE activity in *INPP5D* with GWAS variant context pair (left) and top 15 TFs important in prediction (right).** In the left panel, the x-axis is the actual dfCRE activity value and the y-axis is the predicted value. Each dot is a dfCRE. The red dashed line is the fitted curve. In the right panel, the x-axis is the importance from the random forest model and each row represents a TF.

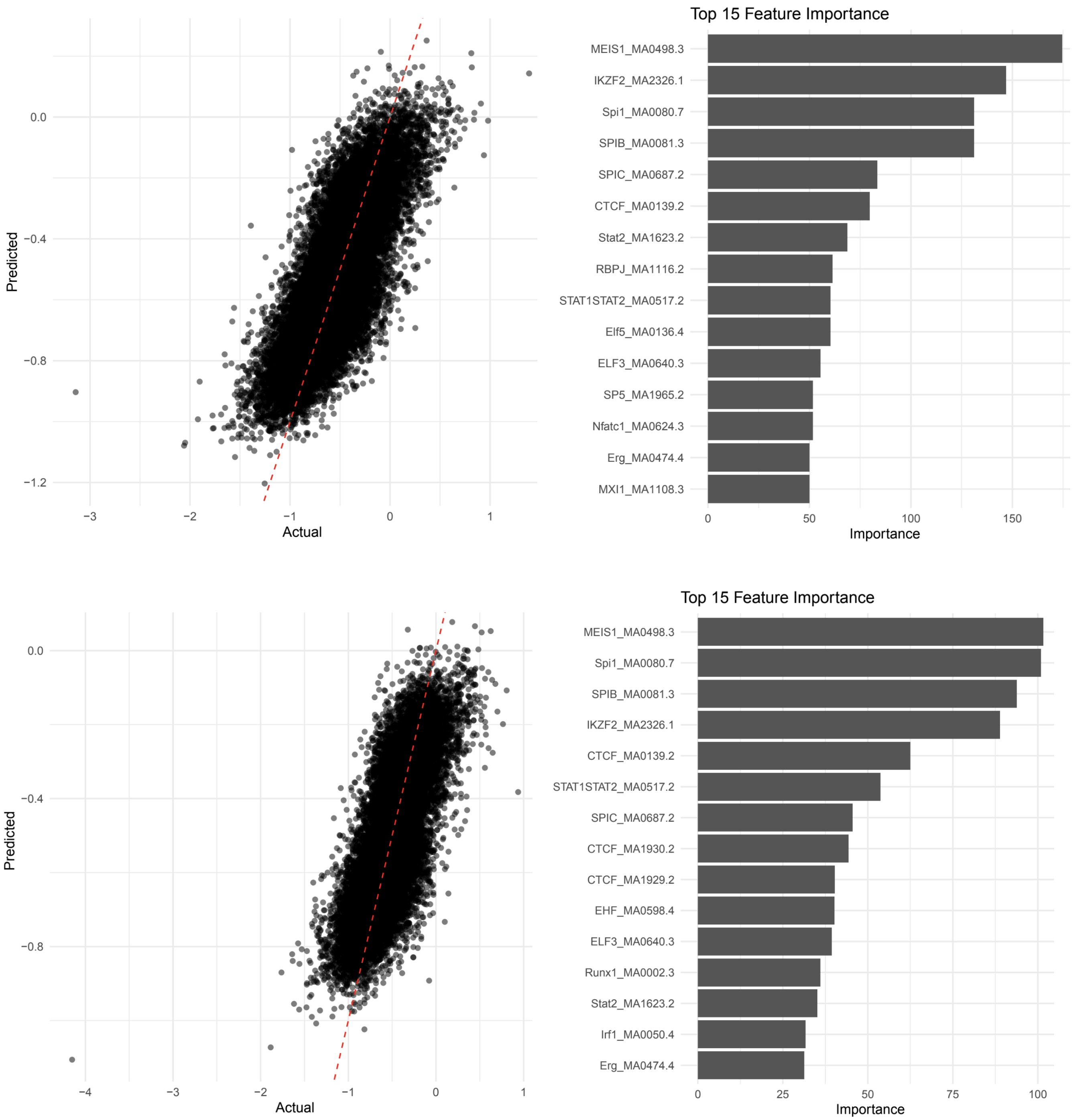


**Fig S17. Random forest model for prediction of dfCRE activity and top 15 important TFs in *SORL1*$^{A528T}$ (above) or *SORL1* KO (below) context pair.** In the left panel of each row, the x-axis is the actual dfCRE activity value and the y-axis is the predicted value. Each dot is a dfCRE. The red dashed line is the fitted curve. In the right panel of each row, the x-axis is the importance from the random forest model and each row represents a TF.

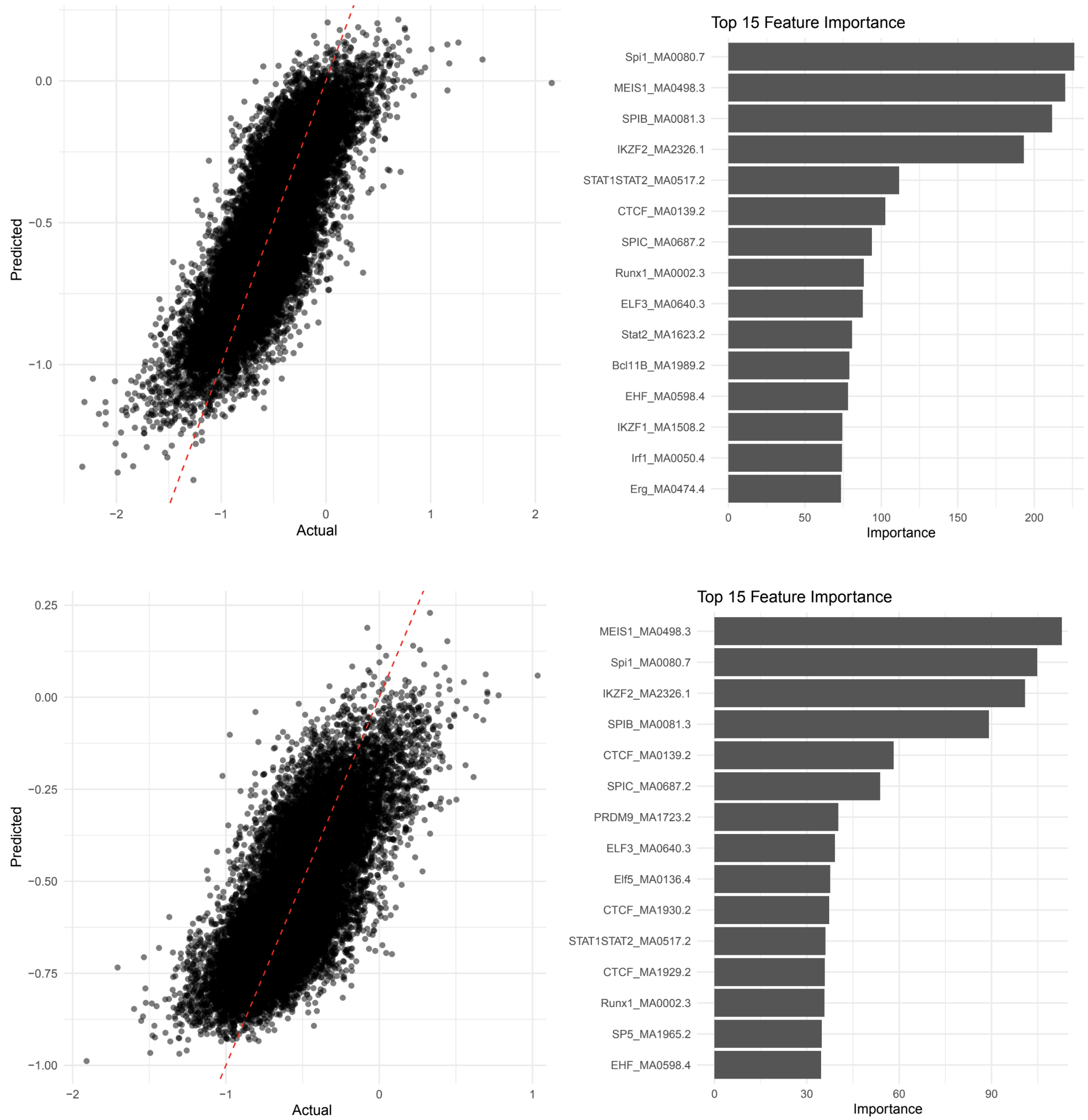


**Fig S18. Random forest model for prediction of dfCRE activity and top 15 important TFs in *TREM2*$^{R47H}$ (above) or *TREM2* KO (below) context pair.** In the left panel of each row, the x-axis is the actual dfCRE activity value and the y-axis is the predicted value. Each dot is a dfCRE. The red dashed line is the fitted curve. In the right panel of each row, the x-axis is the importance from the random forest model and each row represents a TF.

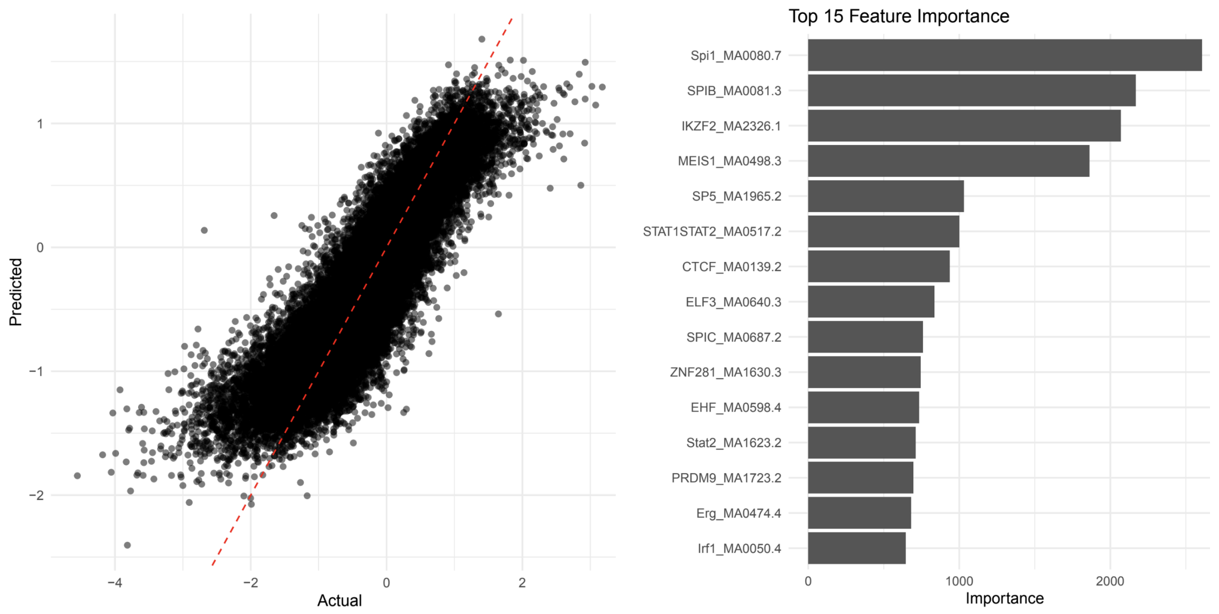


**Fig S19. Random forest model for prediction of dfCRE activity in co-culture with organoids context pair (left) and top 15 TFs important in prediction (right).** In the left panel, the x-axis is the actual dfCRE activity value and the y-axis is the predicted value. Each dot is a dfCRE. The red dashed line is the fitted curve. In the right panel, the x-axis is the importance from the random forest model and each row represents a TF.

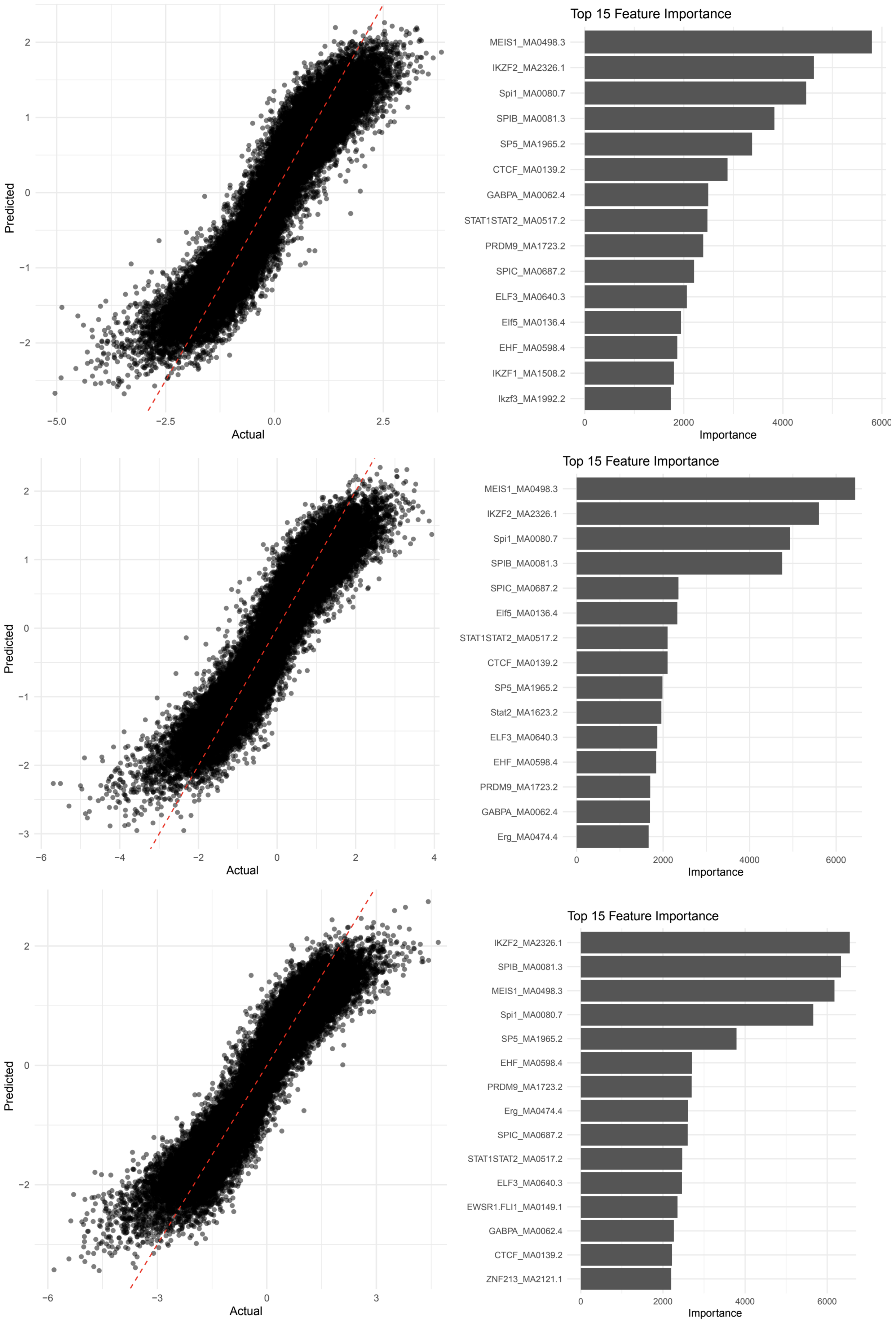


**Fig S20. Random forest model for prediction of dfCRE activity and top 15 important TFs in context pairs of xenotransplantation of 7 days (above), 12 days (middle), or 8 weeks (below).** In the left panel of each row, the x-axis is the actual dfCRE activity value and the y-axis is the predicted value. Each dot is a dfCRE. The red dashed line is the fitted curve. In the right panel of each row, the x-axis is the importance from the random forest model and each row represents a TF..

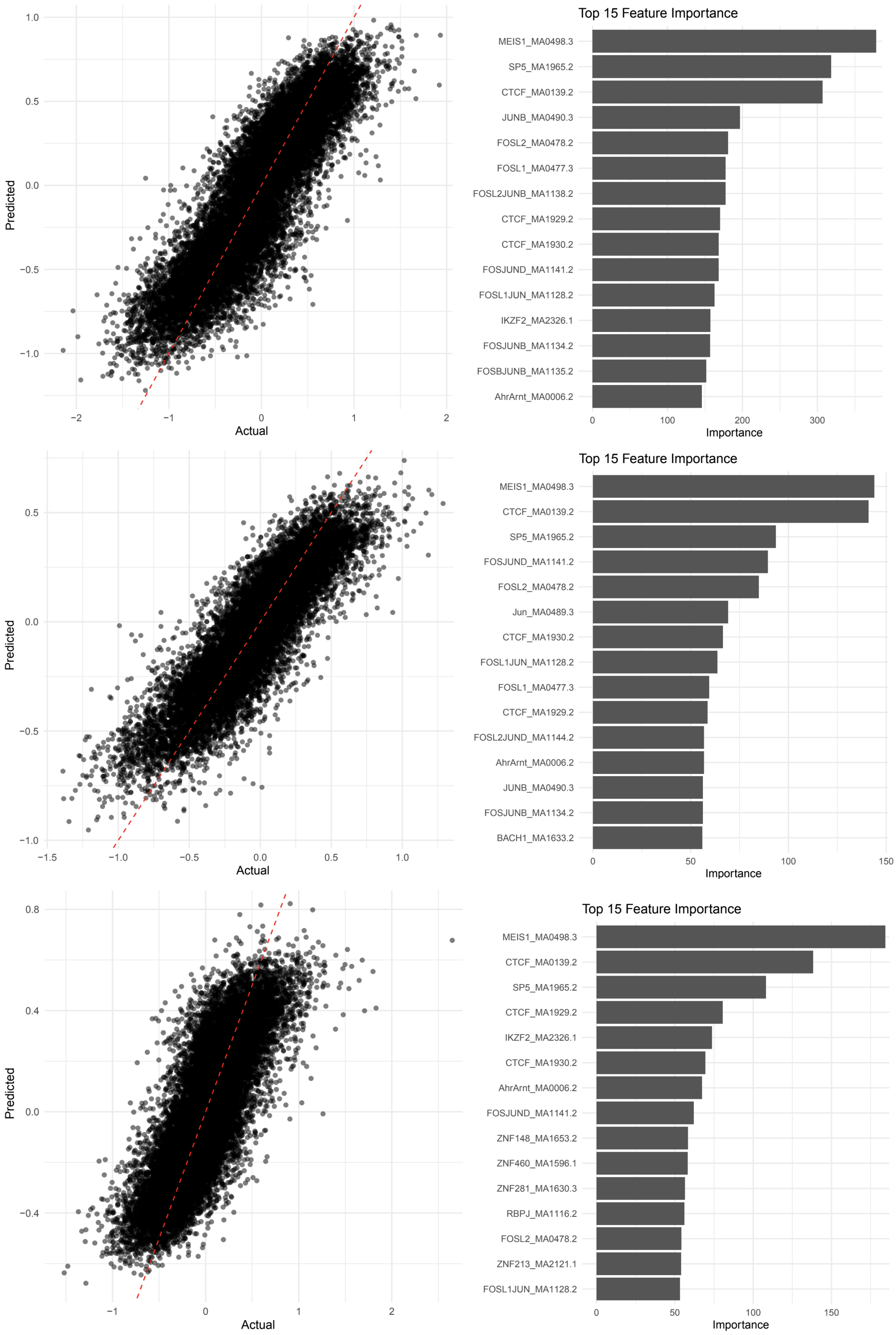


**Fig S21. Random forest model for prediction of dfCRE activity and top 15 important TFs in context pairs of HIV activated vs. HIV latent (above), uninfected (middle), or HIV latent vs. uninfected (below).** In the left panel of each row, the x-axis is the actual dfCRE activity value and the y-axis is the predicted value. Each dot is a dfCRE. The red dashed line is the fitted curve. In the right panel of each row, the x-axis is the importance from the random forest model and each row represents a TF.

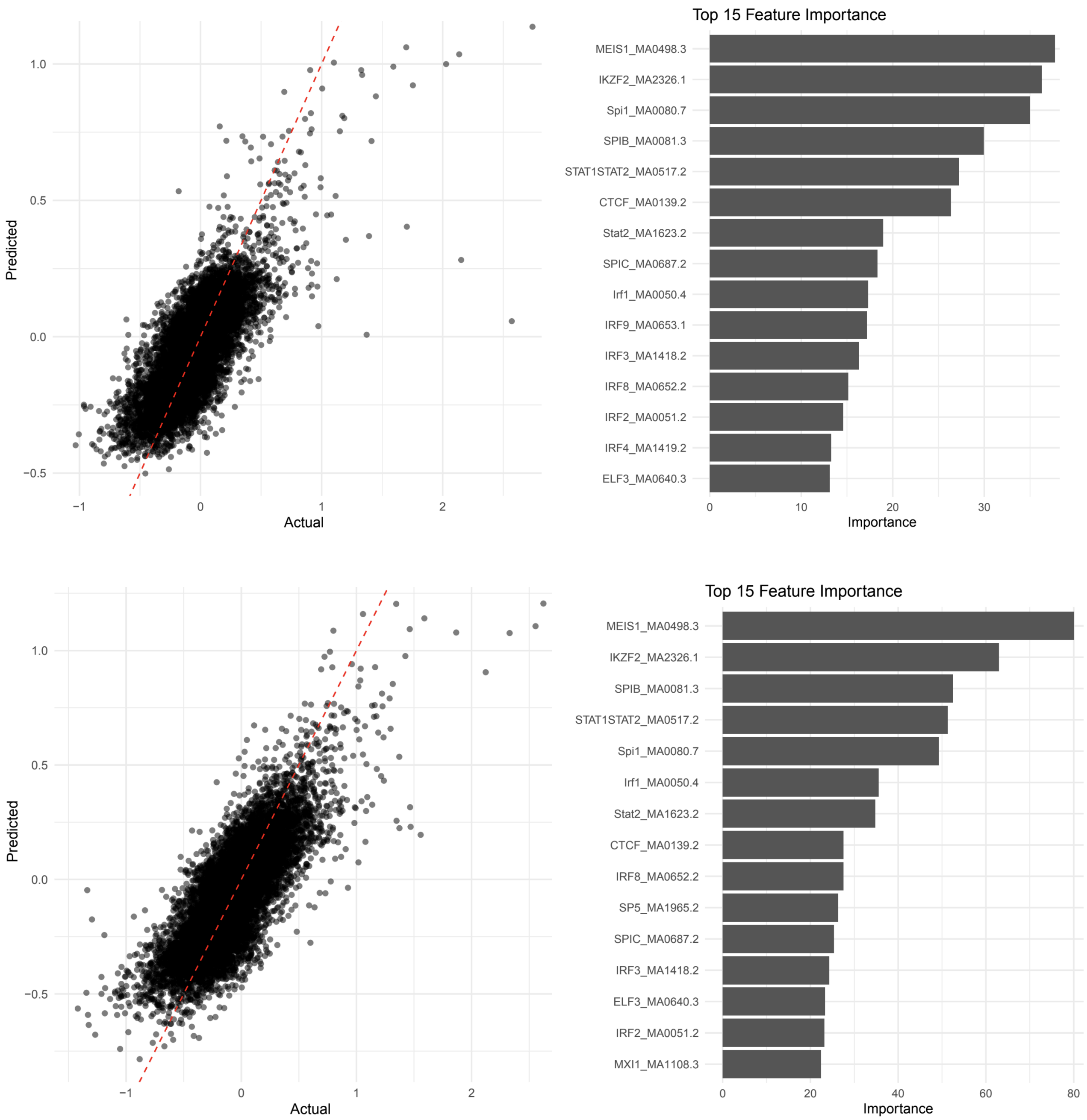


**Fig S22. Random forest model for prediction of dfCRE activity and top 15 important TFs in ESC H1-derived microglia (above) or iPSC WTC11-derived (below) under IFN-$\beta$ stimulation.** In the left panel of each row, the x-axis is the actual dfCRE activity value and the y-axis is the predicted value. Each dot is a dfCRE. The red dashed line is the fitted curve. In the right panel of each row, the x-axis is the importance from the random forest model and each row represents a TF.

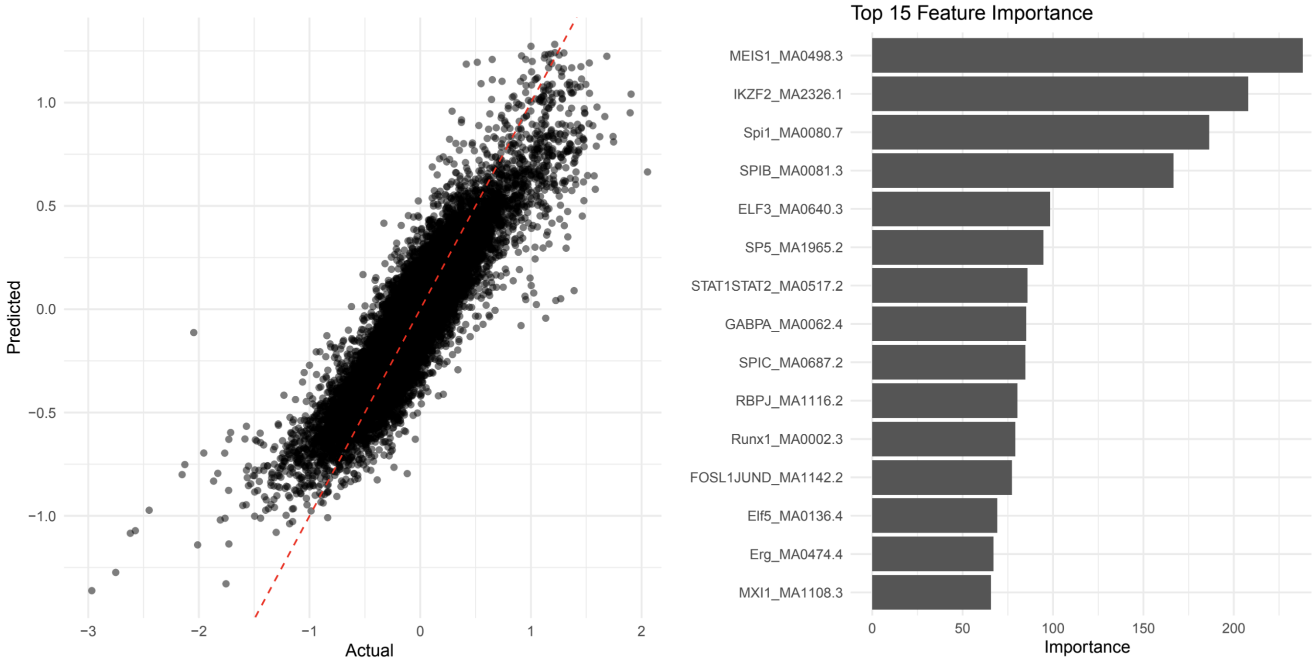


**Fig S23. Random forest model for prediction of dfCRE activity in CRISPR-edited *CLU* context pair (left) and top 15 TFs important in prediction (right).** In the left panel, the x-axis is the actual dfCRE activity value and the y-axis is the predicted value. Each dot is a dfCRE. The red dashed line is the fitted curve. In the right panel, the x-axis is the importance from the random forest model and each row represents a TF.

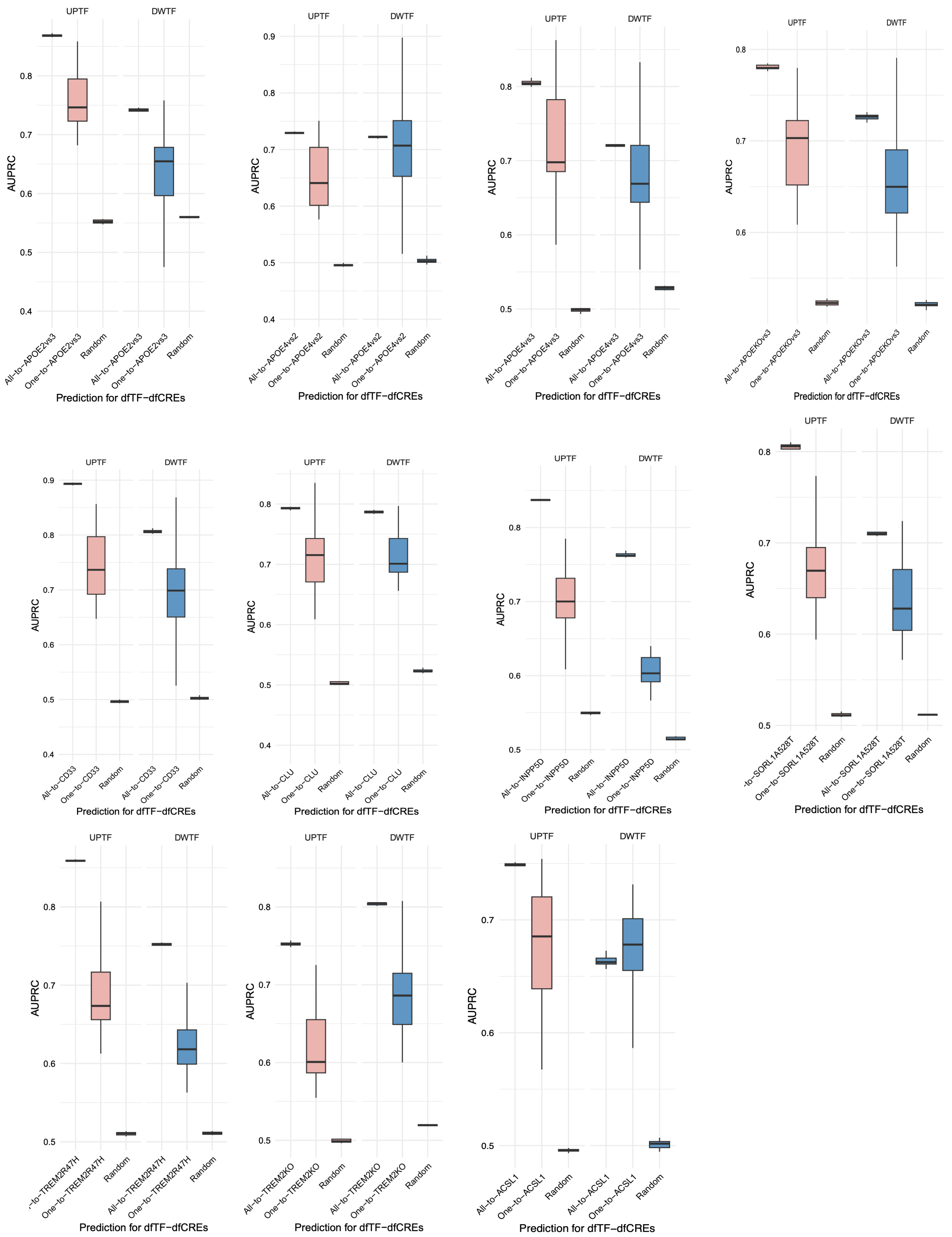


**Fig S24. Embedding-based prediction for dfTF-dfCRE linkings in the target contexts of genetic perturbations (*APOE4* vs. *APOE2*, *APOE4* vs. *APOE3*, *APOE* KO, *CD33*, *CLU*, *INPP5D*, *SORL1*$^{A528T}$, *TREM2*$^{R47H}$, *TREM2* KO) and high-LD context.** For each panel, the x-axis is the predicted context and the y-axis is the AUPRC for assessing prediction. Each box is from ten iterations of prediction for upregulated dfTF-dfCREs (pink) or downregulated ones (blue). The box starts in the first quantile (25%) and ends in the third (75%). The line inside the box represents the median.

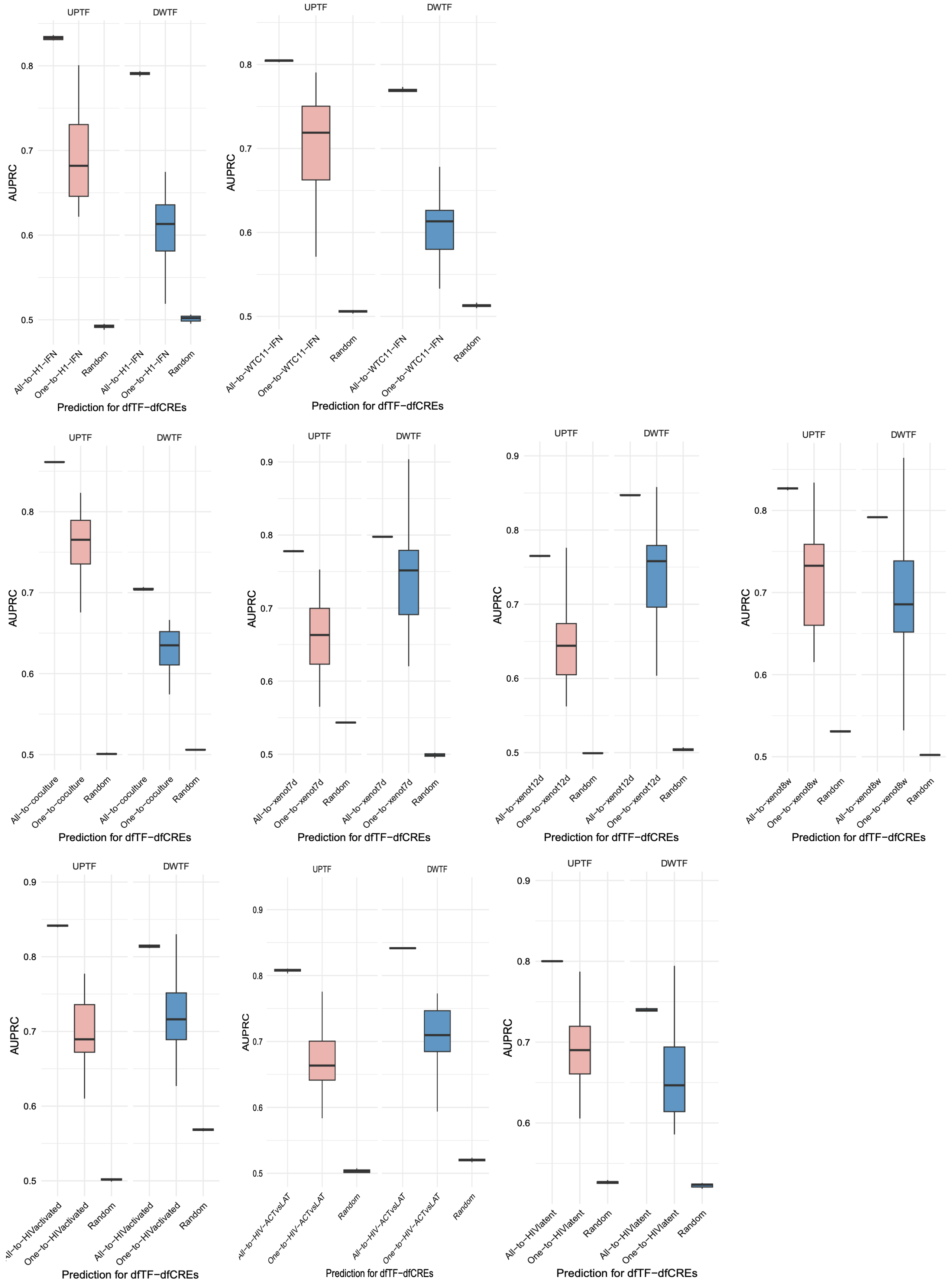


**Fig S25. Embedding-based prediction for dfTF-dfCRE linkings in the target context of IFN-β (in iPSC H1- or WTC11-derived microglia), co-culture with organoids, xenotransplantation (after 7 days, 12 days, and 8 weeks), and HIV infection (HIV-activated vs. HIV-uninfected, HIV-activated vs. HIV-latent, HIV-latent vs. HIV-uninfected).** For each panel, the x-axis is the predicted context and the y-axis is the AUPRC for assessing prediction. Each box is from ten iterations of prediction for upregulated dfTF-dfCREs (pink) or downregulated ones (blue). The box starts in the first quantile (25%) and ends in the third (75%). The line inside the box represents the median.

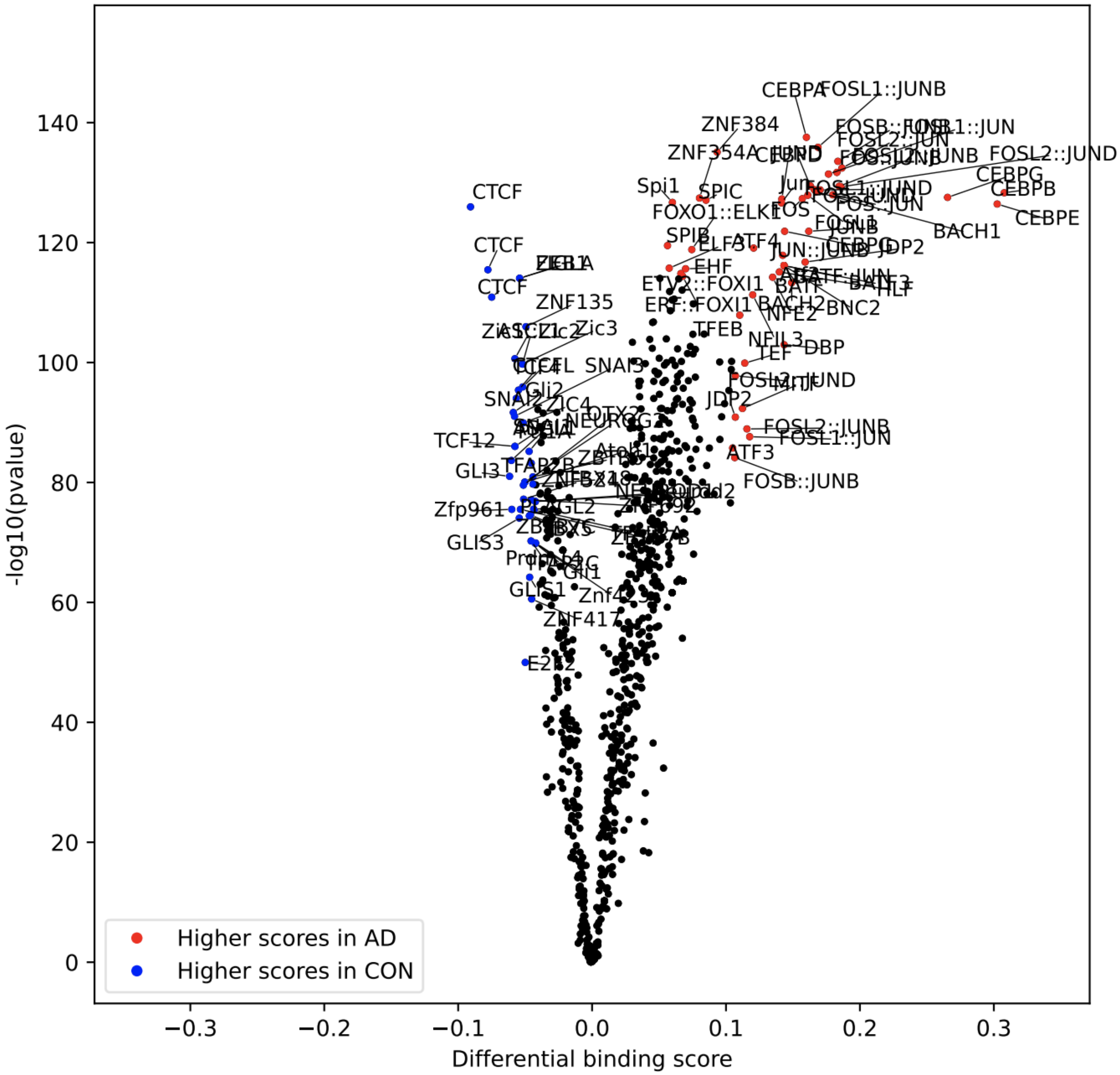


**Fig S26. Differential TF footprint score in the context of AD.** The x-axis is differential footprint score and the y axis is -$\log_{10}$(p value). Each dot is a TF motif. Red represents significantly upregulated TFs in the comparison and blue represents significantly downregulated ones.

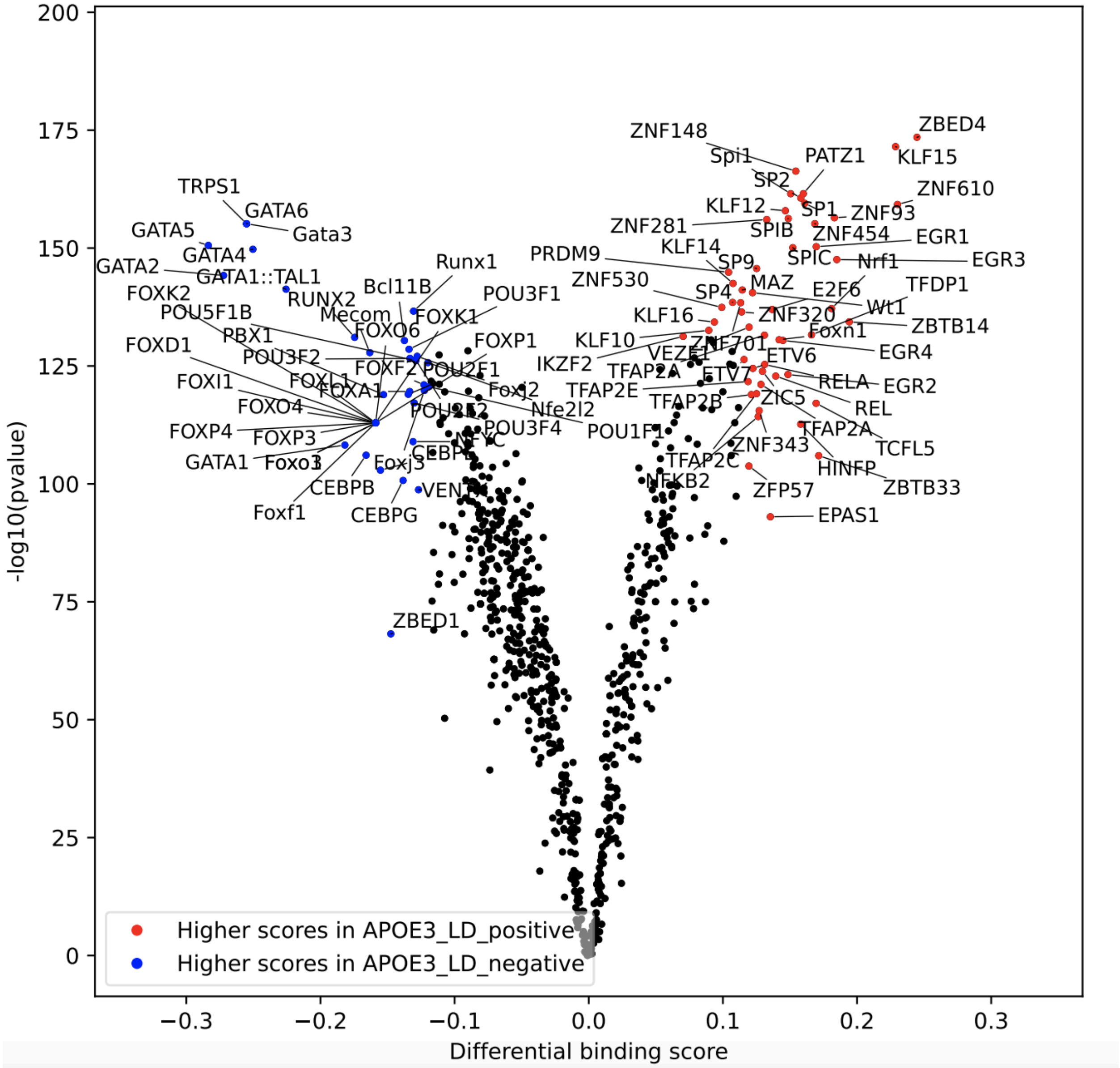


**Fig S27. Differential TF footprint score in the high-LD context.** The x axis is differential footprint score and y axis is -$\log_{10}$(p value). Each dot is a TF motif. Red represents significantly upregulated TFs in the comparison and blue represents significantly downregulated ones.

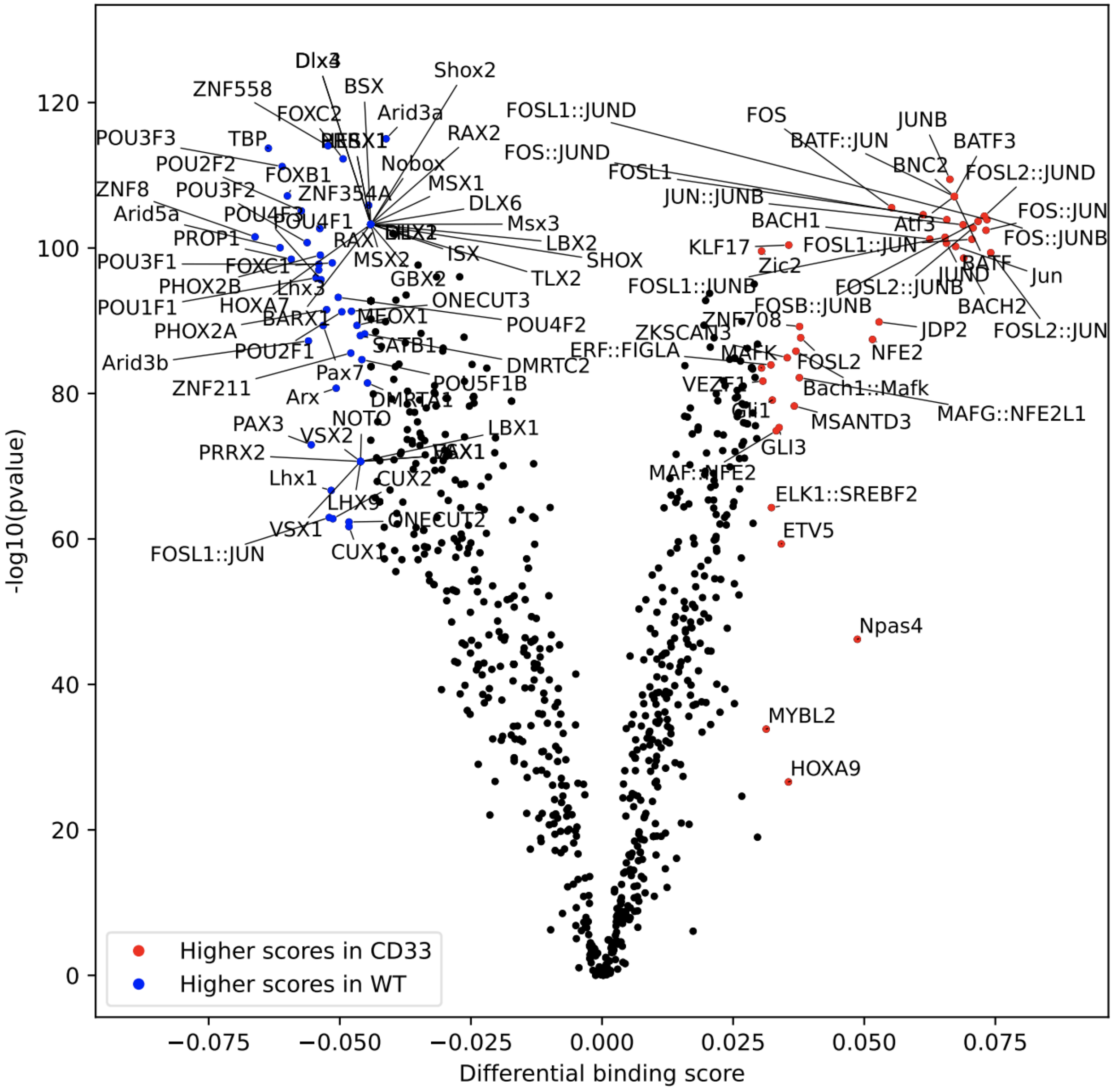


**Fig S28. Differential TF footprint score in the context of *CD33* with GWAS variant.** The x axis is differential footprint score and y axis is -log$_{10}$(p value). Each dot is a TF motif. Red represents significantly upregulated TFs in the comparison and blue represents significantly downregulated ones.

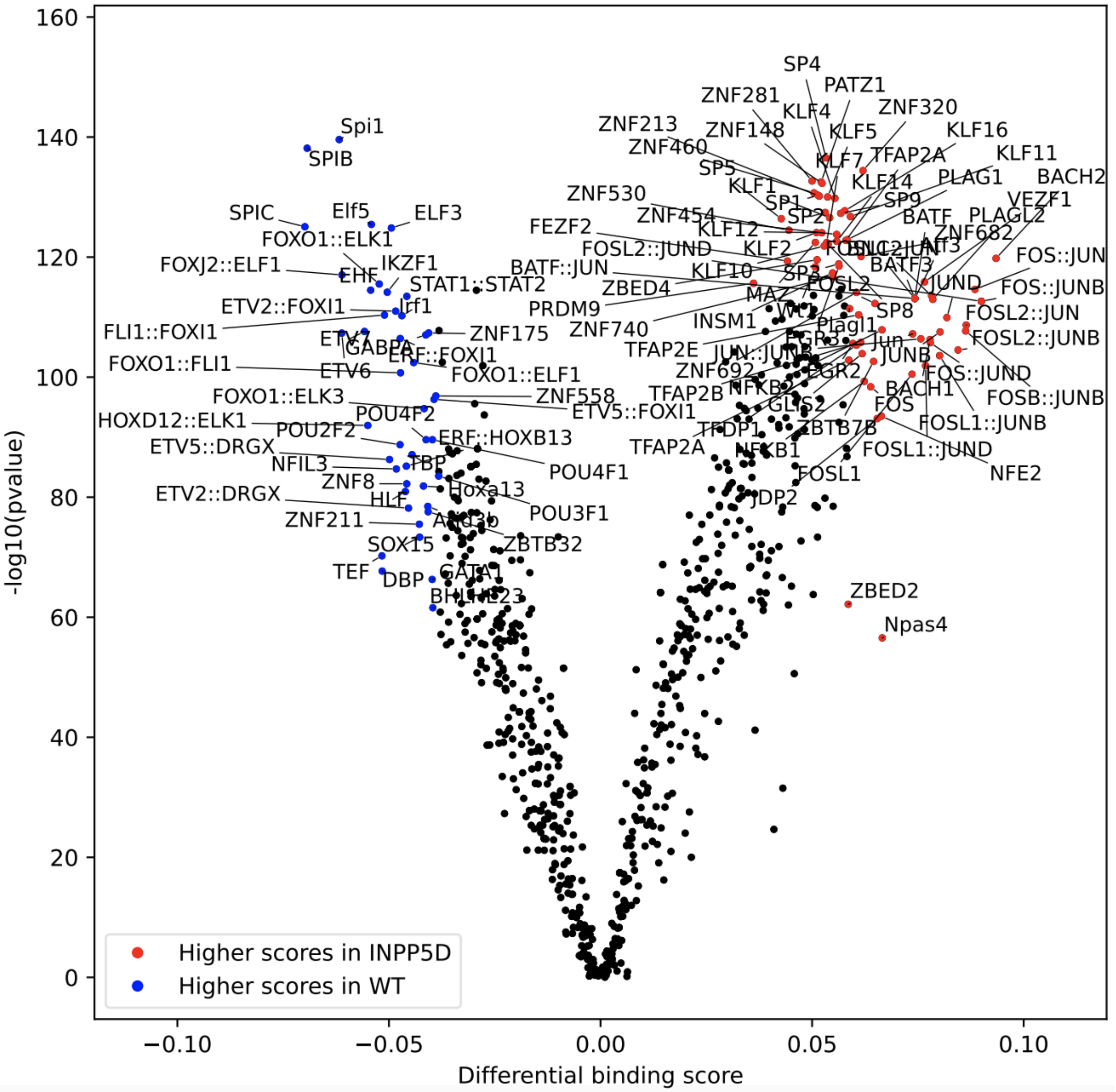


**Fig S29. Differential TF footprint score in the context of *INPP5D* with GWAS variant.** The x axis is differential footprint score and y axis is -log$_{10}$(p value). Each dot is a TF motif. Red represents significantly upregulated TFs in the comparison and blue represents significantly downregulated ones.

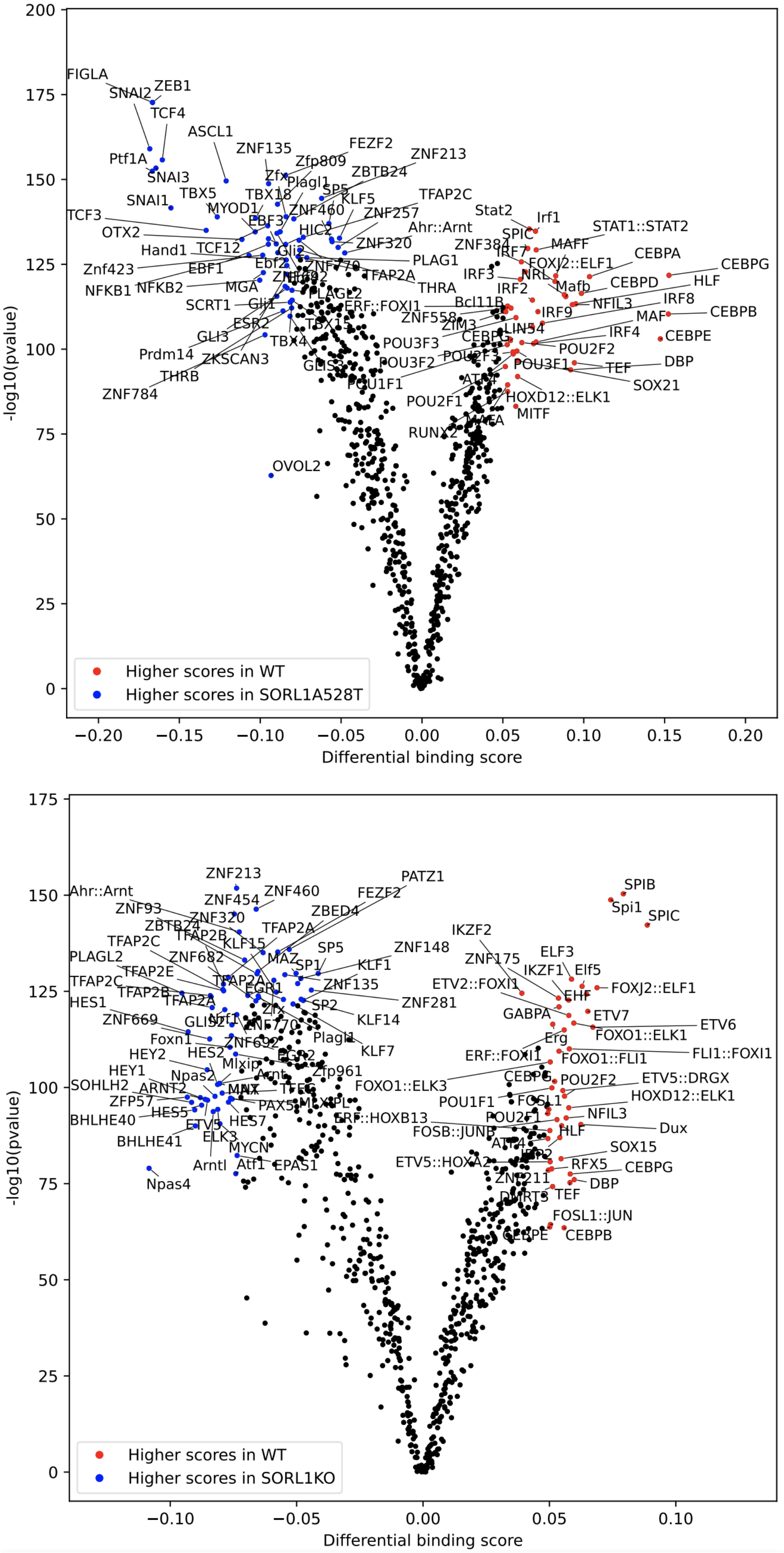


**Fig S30. Differential TF footprint score in the context of *SORL1*$^{A528T}$ (above) or *SORL1* KO (below) vs. control.** The x axis is differential footprint score and y axis is -log$_{10}$(p value). Each dot is a TF motif. Red represents significantly upregulated TFs in the comparison and blue represents significantly downregulated ones.

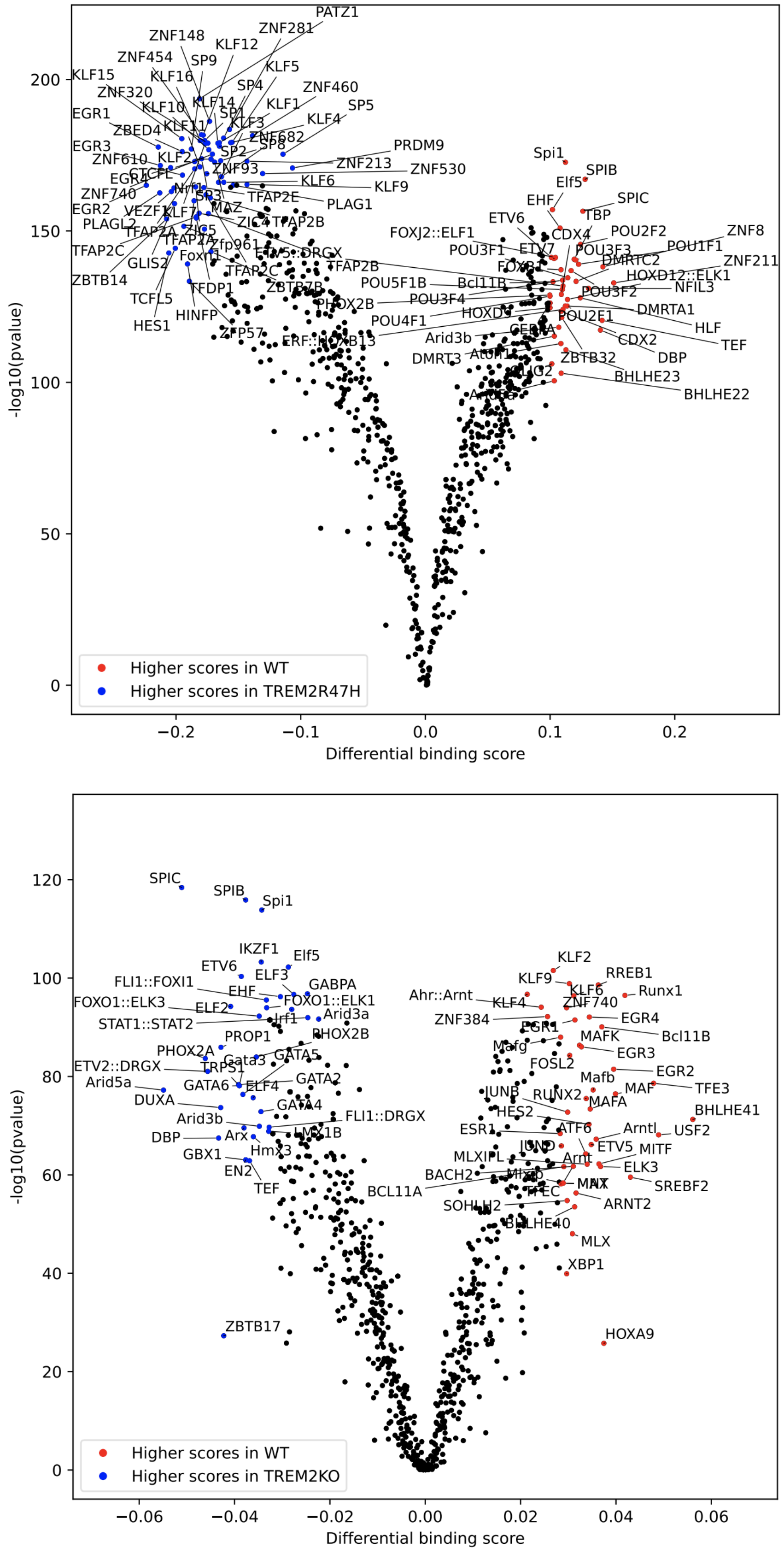


**Fig S31. Differential TF footprint score in the context of *TREM2*$^{R47H}$ (above) or *TREM2* KO (below) vs. control.** The x axis is differential footprint score and y axis is -log$_{10}$(p value). Each dot is a TF motif. Red represents significantly upregulated TFs in the comparison and blue represents significantly downregulated ones.

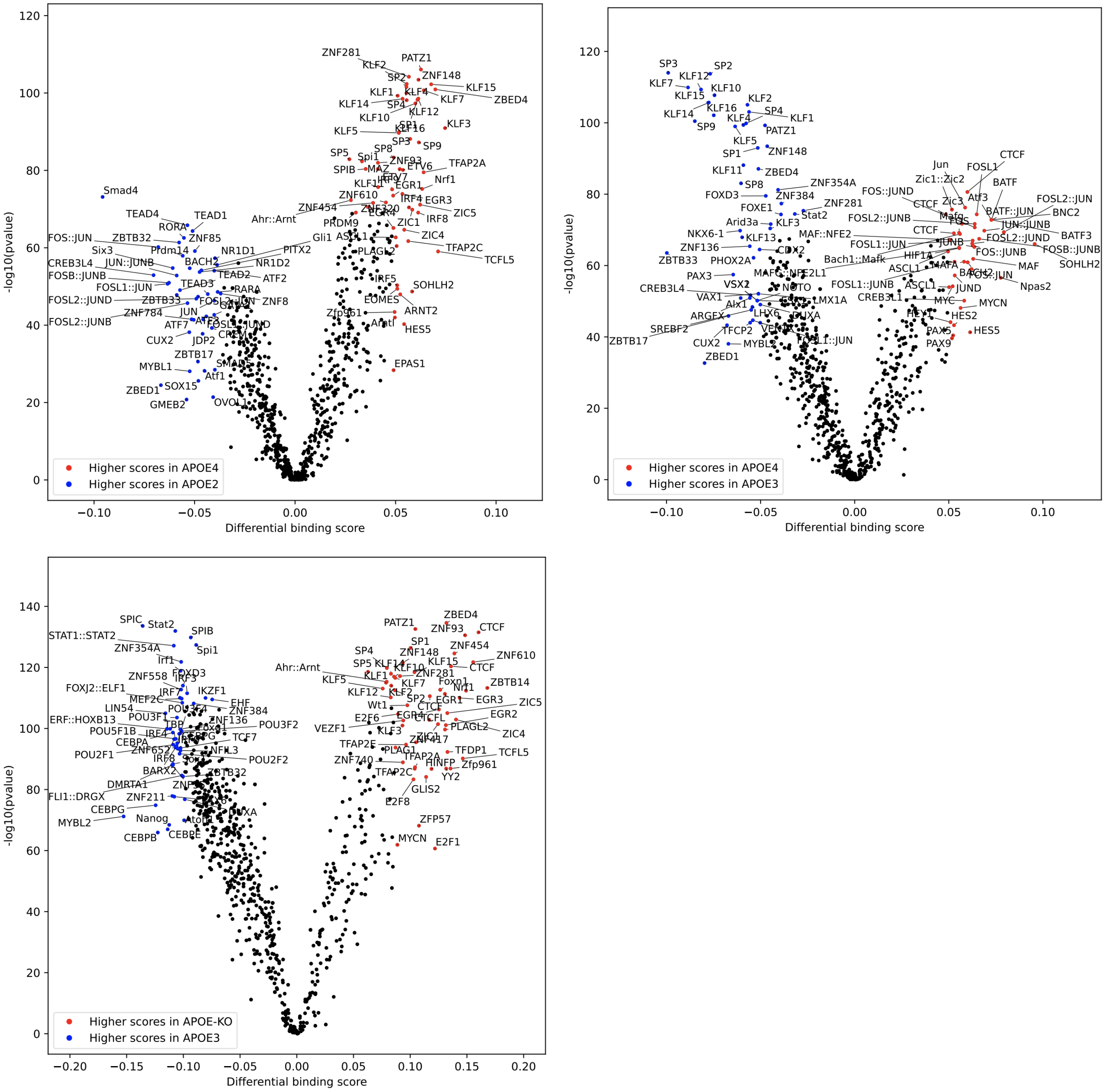


**Fig S32. Differential TF footprint score in the context of *APOE4* vs. *APOE2* (upper left), *APOE4* vs. *APOE3* (upper right), or *APOE* KO vs. *APOE3* (below).** The x-axis represents the differential footprint score and the y-axis represents -log$_{10}$($P$). Each dot is a TF motif. Red represents significantly upregulated TFs in the comparison, and blue represents significantly downregulated ones.

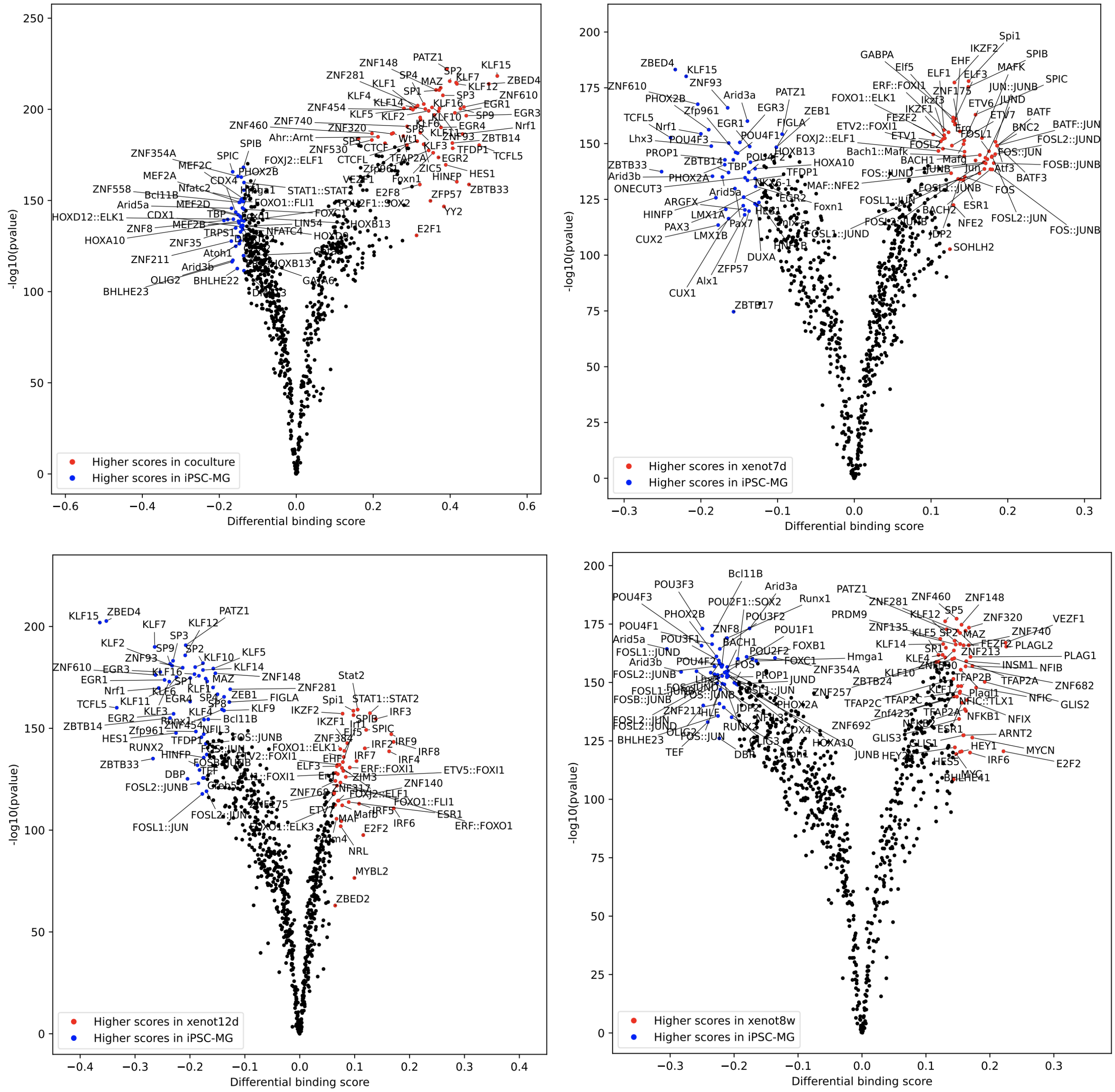


**Fig S33. Differential TF footprint score in the contexts of co-culture with organoids (upper left), xenotransplantation of 7 days (upper right), 12 days (left lower), or 8 weeks (right lower).** The x axis is differential footprint score and y axis is -$\log_{10}$(p value). Each dot is a TF motif. Red represents significantly upregulated TFs in the comparison and blue represents significantly downregulated ones.

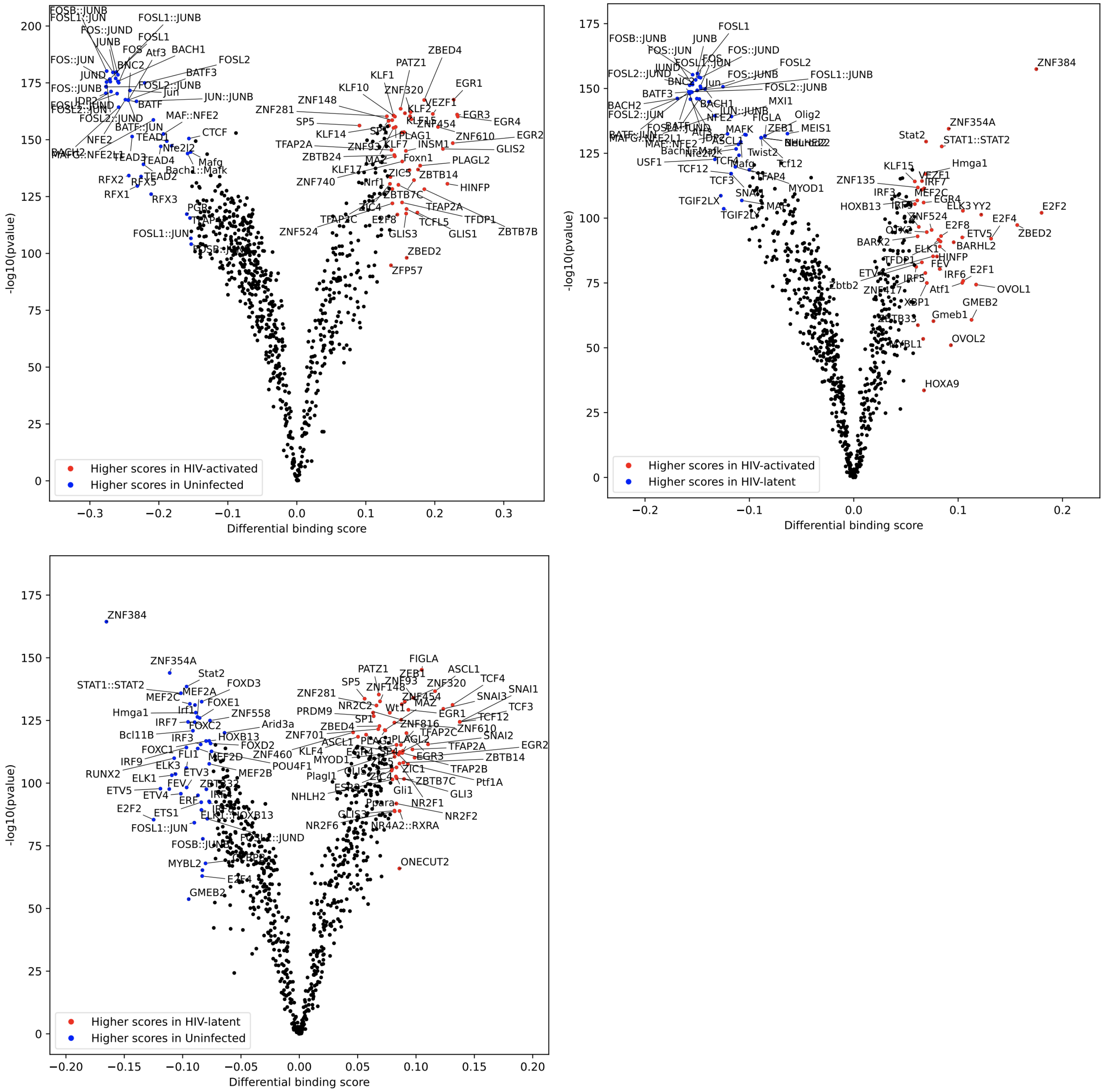


**Fig S34. Differential TF footprint score in the context of HIV activated vs. uninfected (upper left), HIV activated vs. HIV latent (upper right), or HIV latent vs. uninfected (below).** The x axis is differential footprint score and y axis is -$\log_{10}$(p value). Each dot is a TF motif. Red represents significantly upregulated TFs in the comparison and blue represents significantly downregulated ones.

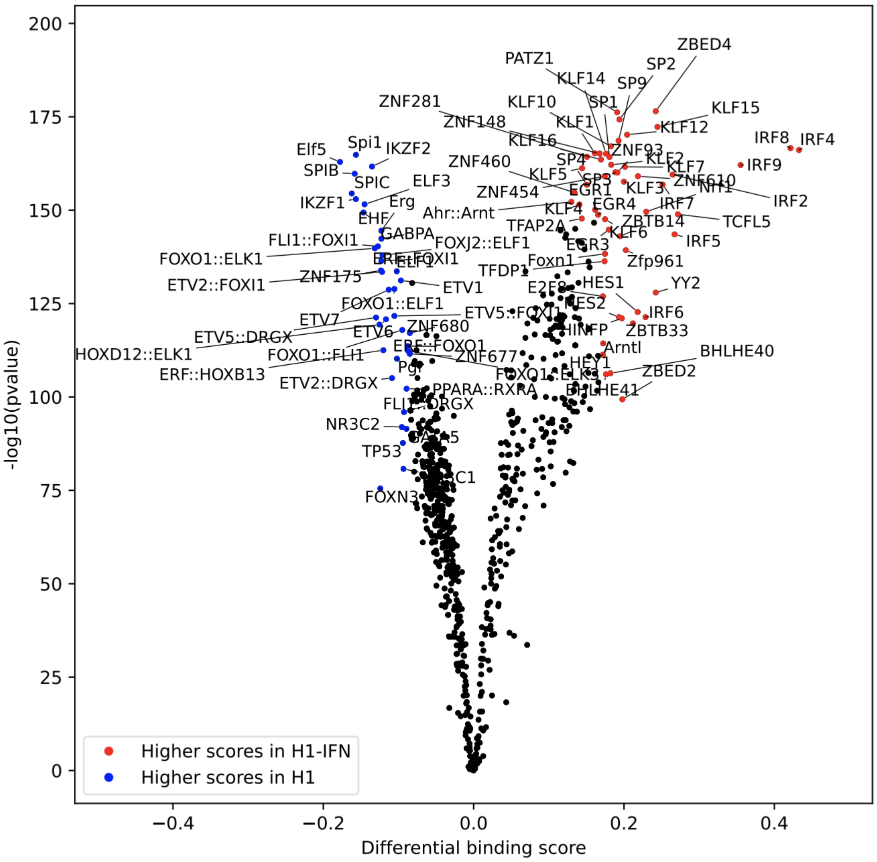


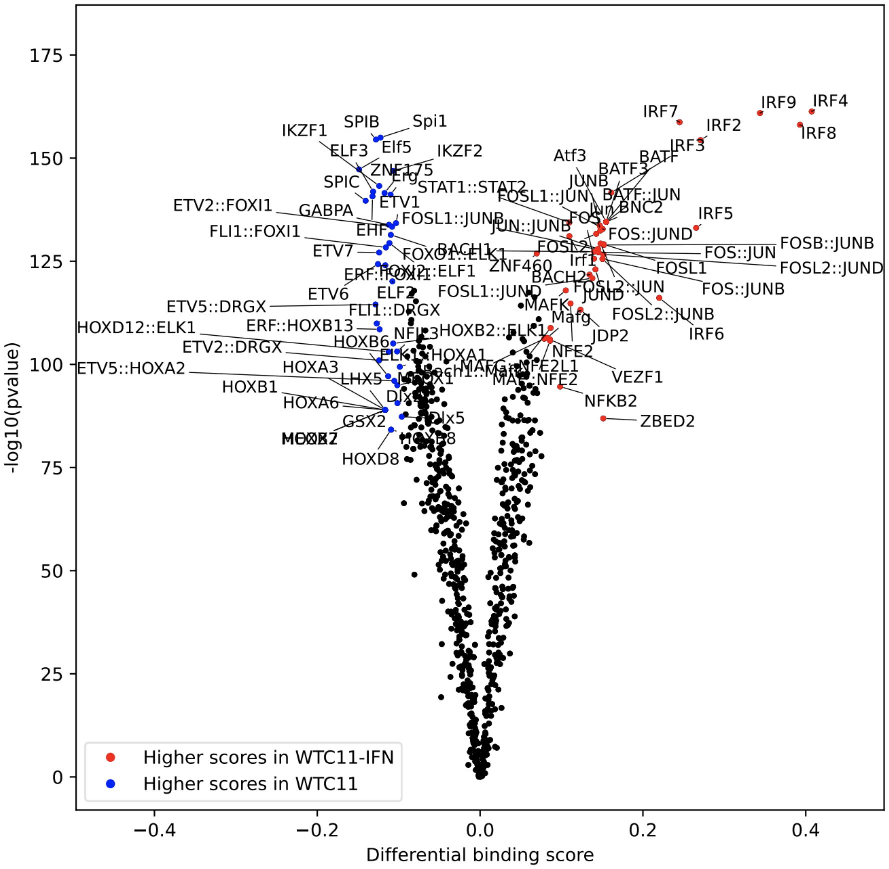


**Fig S35. Differential TF footprint score in ESC H1-derived microglia (above) or iPSC WTC11-derived microglia (below) under IFN-$\beta$ stimulation.** The x axis is differential footprint score and y axis is -$\log_{10}$(p value). Each dot is a TF motif. Red represents significantly upregulated TFs in the comparison and blue represents significantly downregulated ones.

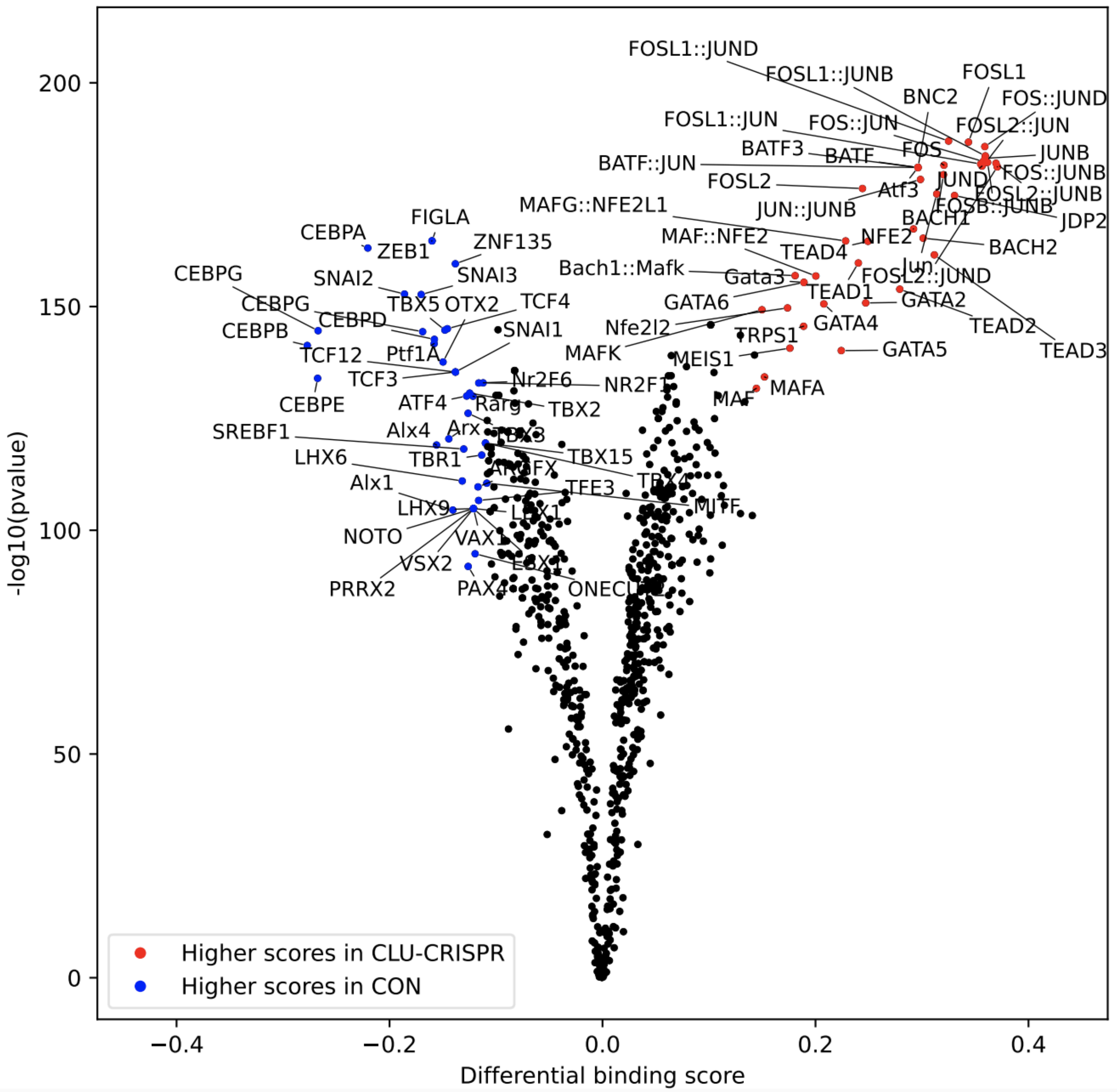


**Fig S36. Differential TF footprint score in the context of CRISPR-edited *CLU*.** The x axis is differential footprint score and y axis is -$\log_{10}$(p value). Each dot is a TF motif. Red represents significantly upregulated TFs in the comparison and blue represents significantly downregulated ones.

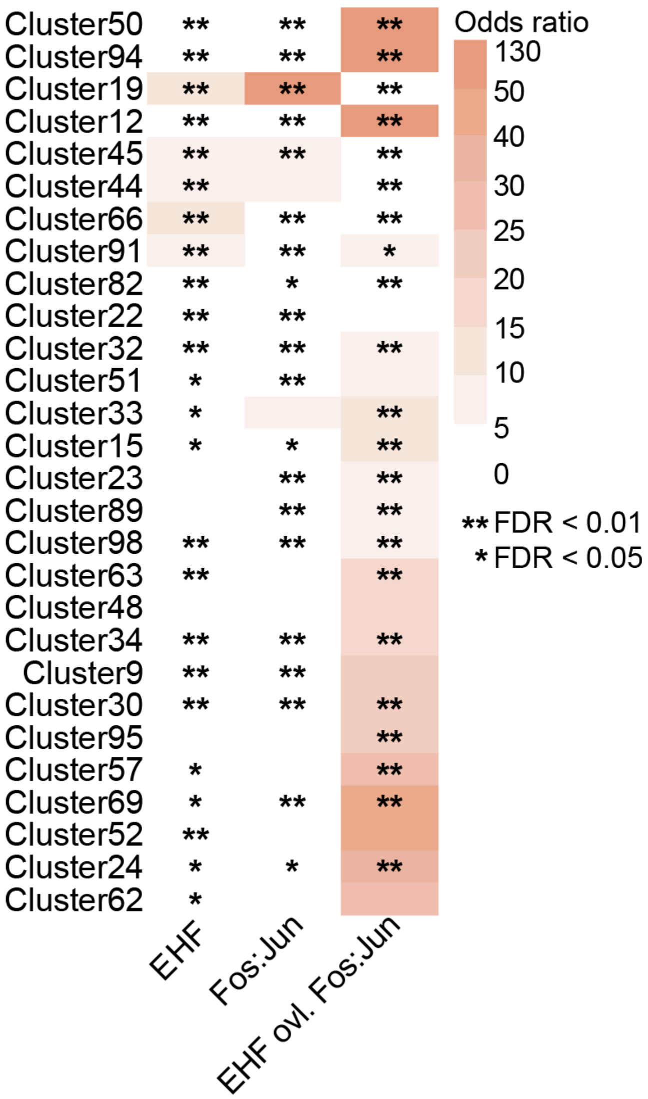


**Fig S37. Enrichment of bound dfCREs of EHF, FOS:JUN, and their co-occupancy in dfCRE modules of AD microglia.** Fisher's exact test was used to determine the statistical significance level, shown with *.

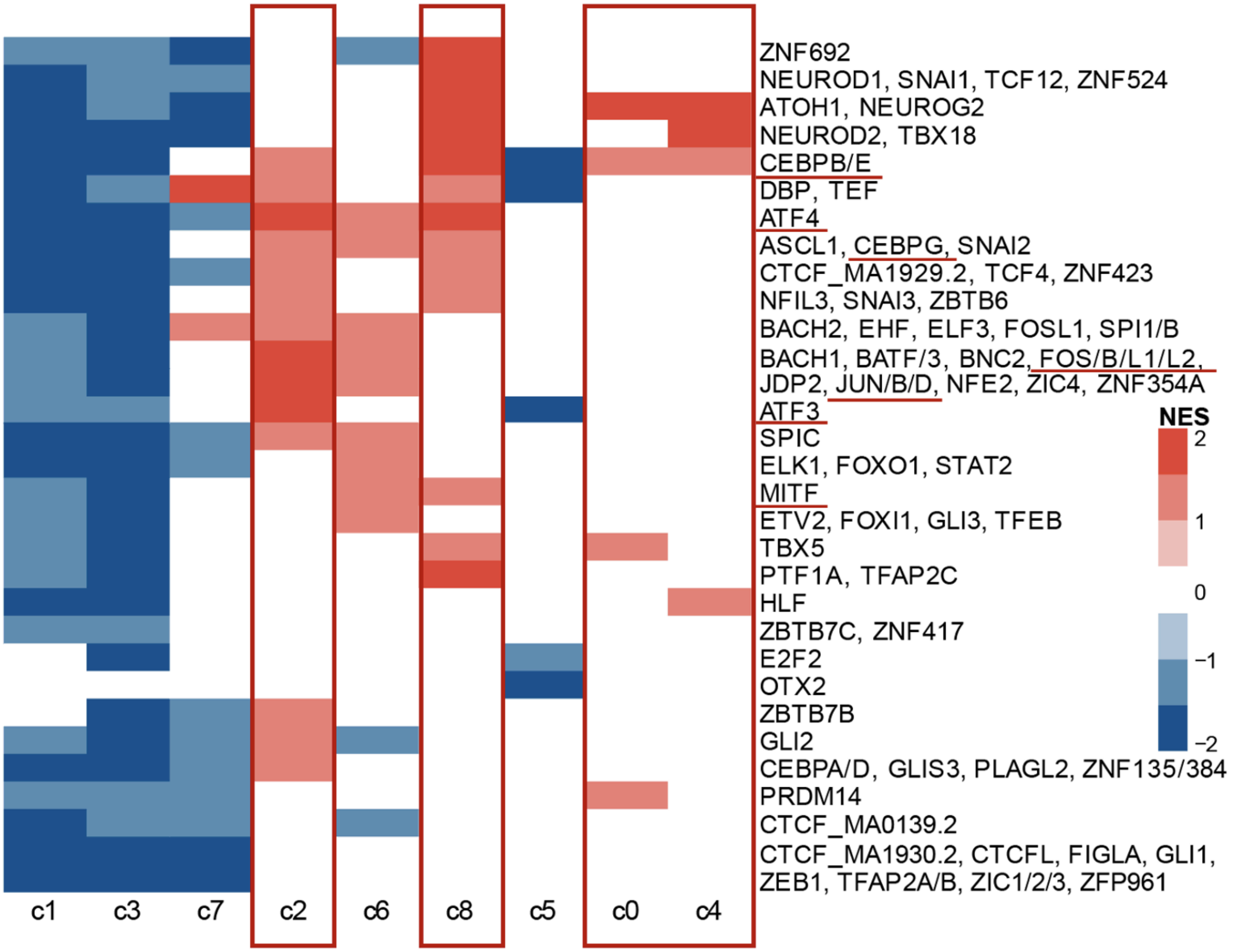


**Fig S38. GSEA enrichment of regulatory circuits of AD microglia in *HuMicA* clusters.** The x axis is a cluster defined by *HuMicA* and the y-axis shows TFs. Red represents positive NES values from GSEA analysis while blue represents negative NES values. The *HuMicA* c0, c2, c4, and c8 were highlighted in a red box and dfTFs enriched in Sun *et al*.'s MG8 were highlighted with red lines.

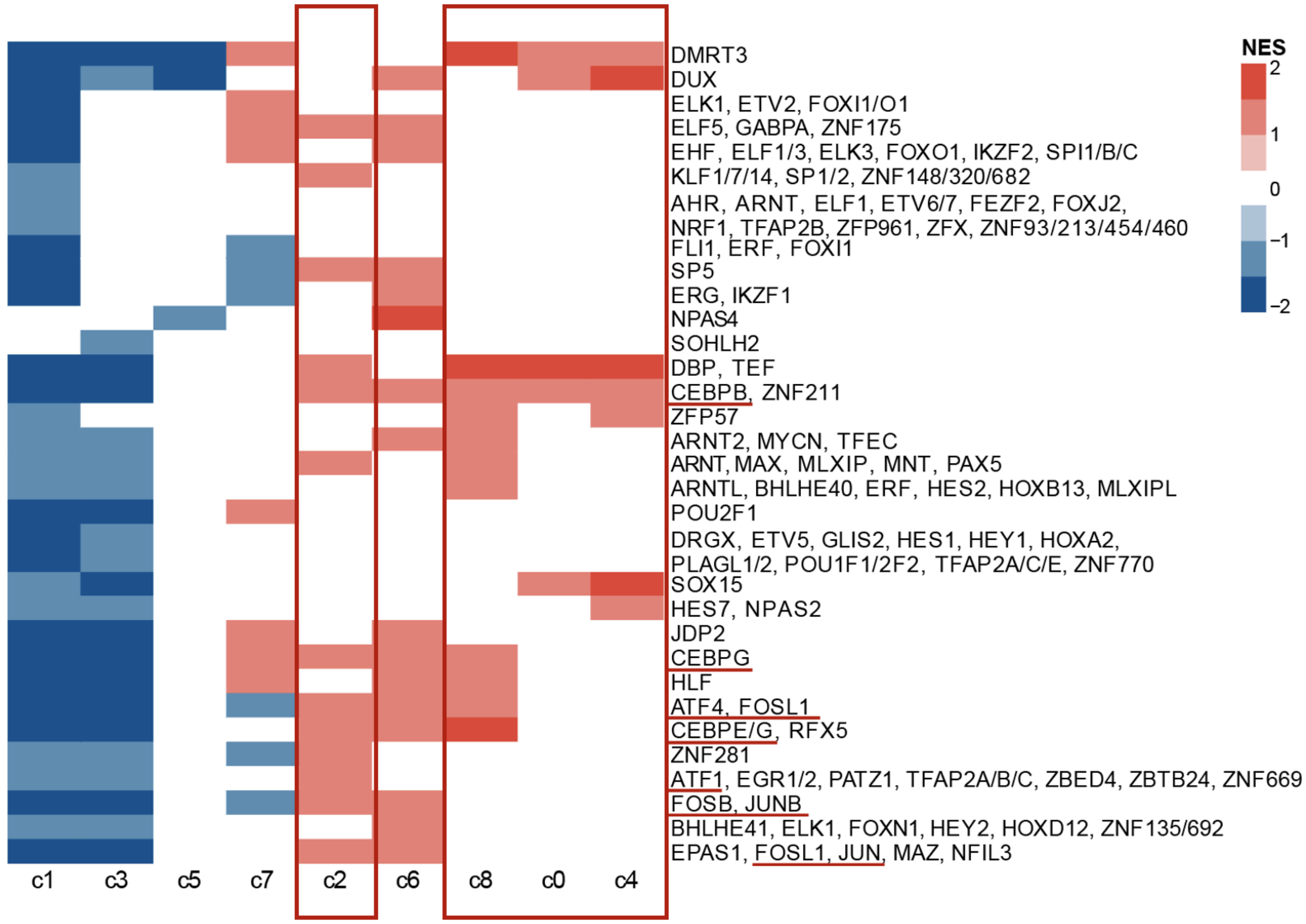


**Fig S39. GSEA enrichment of regulatory circuits of *SORL1* KO in *HuMicA* clusters.** The x-axis is a cluster defined by *HuMicA* and the y-axis shows TFs. Red represents positive NES value from GSEA analysis while blue represents negative NES value. The *HuMicA* c0, c2, c4, and c8 were highlighted in a red box and dfTFs enriched in Sun *et al*.'s MG8 were highlighted with red lines.

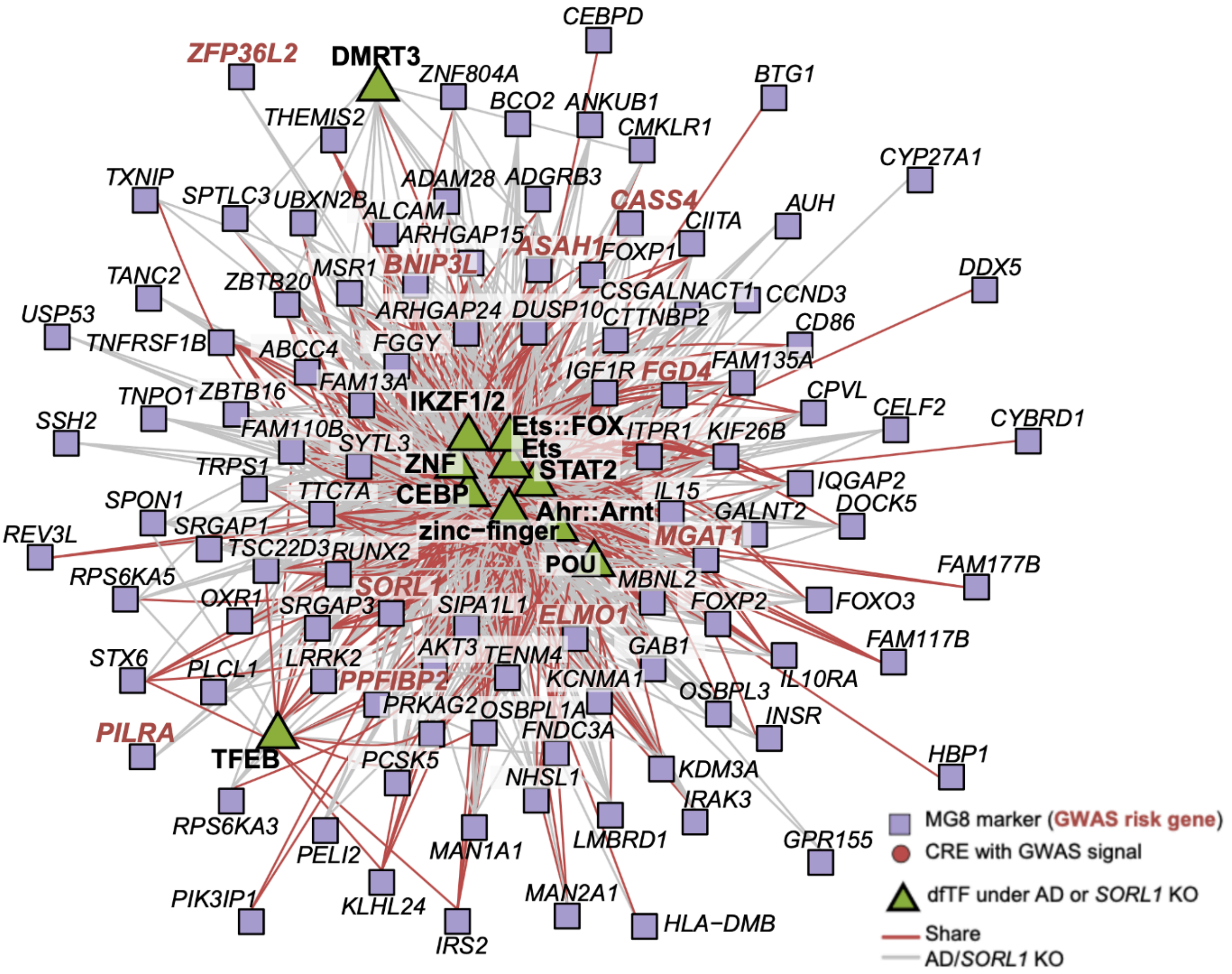


**Fig S40. Novel MG8-enriched regulatory circuits identified in the context of AD or *SORL1* KO.** Purple squares represent MG8 markers. Gene symbols in red bold fonts represent markers also as AD risk genes. Green triangles represent the dfTF families derived from AD or *SORL1* KO microglia. Red lines represent shared links between AD and *SORL1* KO, and grey lines mean specific links in one of the two contexts.

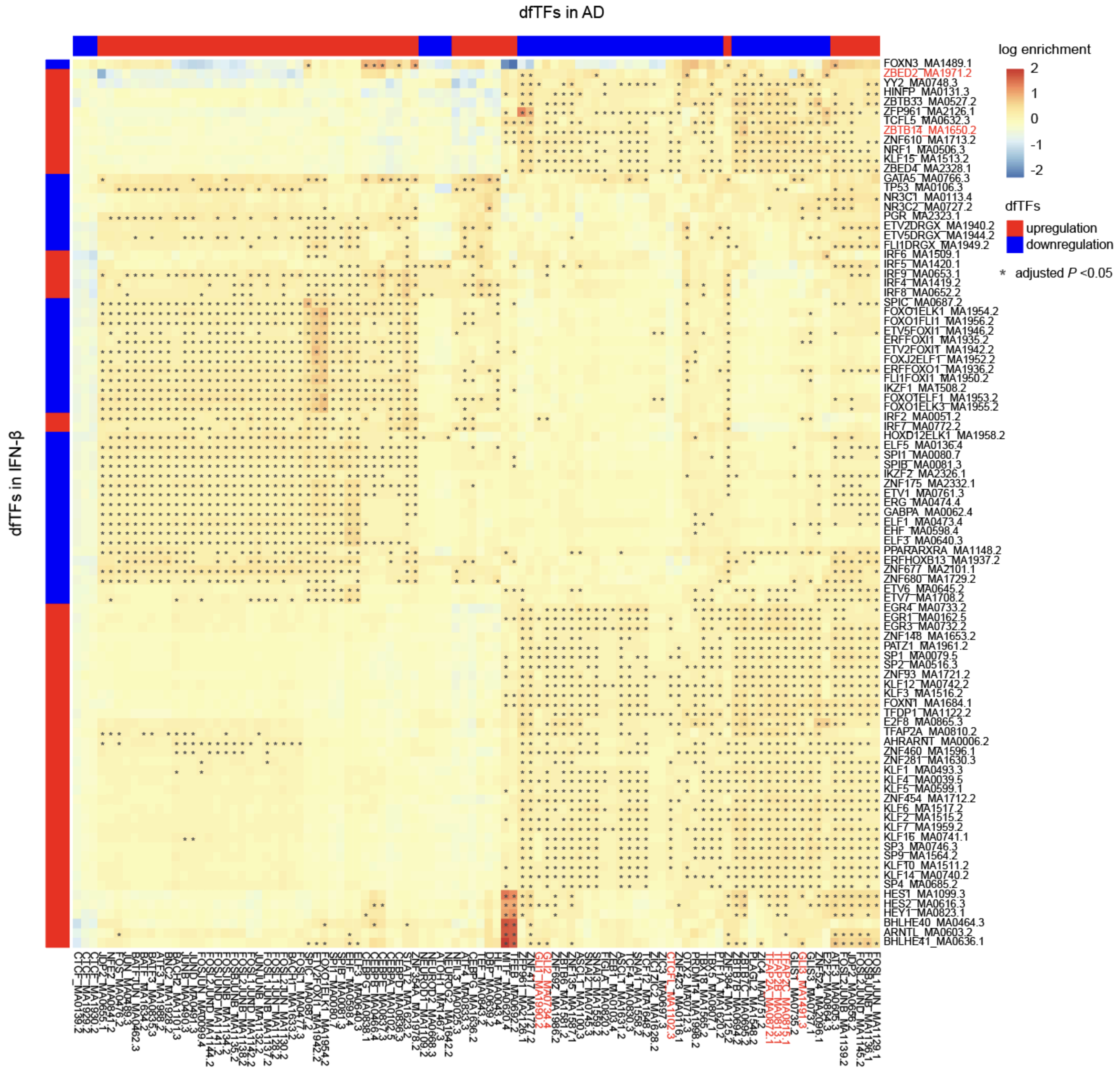


**Fig S41. dfCREs targeted by inflammation-upregulated TFs with putative repressor activity were significantly located near dfCREs targeted by AD dfTFs.** The heatmap shows the enrichment of dfCREs targeted by dfTFs in IFN-***β*** condition localized nearby dfCREs targeted by TFs in AD condition. Each row and column represents a dfTF in IFN-***β*** and AD contexts, respectively. TFs mentioned in the main text are in red.

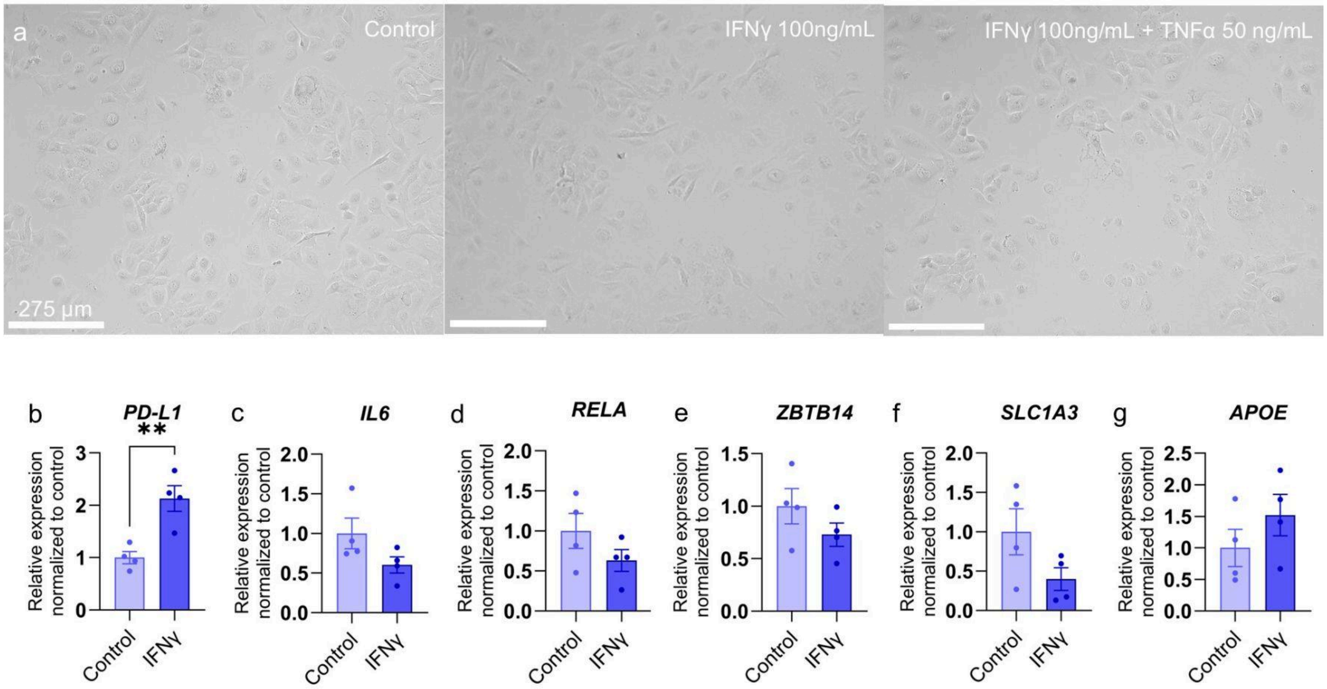


**Fig S42: The relative gene expression of selected genes differentially expressed in connection with *ZBTB14*. a.** Brightfield images of HMC3 cells after 6 hours of treatment with IFNγ (100 ng/mL), and co-treatment with IFNγ (100 ng/mL) and TNFα (50 ng/mL). Scale bar = 275 µm. **b.** Relative gene expression of *PD-L1,* an inflammation response gene (positive control), and **c.** *IL6*, **d.** *RELA*, **e.** *ZBTB14*, **f.** *SLC1A3,* a gene associated with *ZBTB14*, and **g.** *APOE*. Data plotted on the graph as average ± standard deviation; each dot represents one biological replicate from two technical replicates. n = 4 biological replicates per group $^{**}P < 0.01$ in student's two-sided t-test.

### Supplementary Tables

**Table S1. Summary of the public datasets used in this study (the AD dataset was from single-nucleus ATAC-seq, others were from bulk ATAC-seq)**

| Dataset | Cell type (derived cell line) | Data source | Perturbations | Controlled conditions |
|---|---|---|---|---|
| Morabito S., *Nat. Genet.*, 2021 | Human microglia from prefrontal cortex tissue | GSE174367 | AD (N = 12) | Healthy (N = 8) |
| Haney M.S., *Nature*, 2024 | iMG (EC11) | GSE254205 | High lipid droplet (LD) in the genetic background of *APOE3/3* (N = 4) after fibrillar Aβ treatment | Low LD in the genetic background of *APOE3/3* (N = 4) after fibrillar Aβ treatment |
| Kitty B. Murphy, *Nat. Commun.*, 2024 | iMG (BIONi010-C) | GSE271384 | *APOE2/0* (N = 5) | *APOE3/0* (N = 5) |
| | | | *APOE4/0* (N = 5) | *APOE3/0* (N = 5) |
| | | | *APOE4/0* (N = 5) | *APOE2/0* (N = 5) |
| | | | *APOE*-KO (N = 5) | *APOE3/0* (N = 5) |
| Zhao X., Mol. *Neurodegener.*, 2025 | iMG (CD04, CD09 for control; CD05, CD07 for CRISPR/Cas9-editing) | GSE263804 | CRISPR/Cas9-edited to homozygous C/C at AD GWAS site rs1532278 of *CLU* (N = 2) | CRISPR/Cas9-edited to homozygous T/T at AD GWAS site rs1532278 of *CLU* (N=38) |
| Liu T., *J. Exp. Med.*, 2020 | hMGL (H9) | GSE153658 | *CD33* SNP rs3865444 (chr 19 C/A minor) (N = 3) | *CD33* (chr19 WT) (N = 3) |
| | | | *INPP5D* SNP rs35349669 (chr 2 C/T minor) (N = 3) | *INPP5D* (chr 2 WT) (N = 3) |
| | | | *TREM2* KO (N = 3) | *TREM2* WT (N = 3) |
| | | | $TREM2^{R47H}$ rs75932628 (chr6) (N = 3) | *TREM2* (chr6 WT) (N = 3) |
| | | | *SORL1* KO (N = 3) | *SORL1* WT (N = 3) |
| | | | $SORL1^{A528T}$ rs2298813 (chr11) (N = 3) | *SORL1* (chr11 WT) (N = 3) |
| Yang X., *Nat. Genet.*, 2023 | iMG (WTC11), hMGL (H1) | GSE173316 | IFN-***β*** on WTC11 (N = 2) | Resting WTC11 (N = 2) |
| | | | IFN-***β*** on H1 (N = 2) | Resting H1 (N = 2) |
| Han C.Z., *Immunity*, 2023 | iMG (EC11) | GSE226690 | Co-cultured with cerebral organoid (N = 2) | iMG EC11 (N = 5) |
| | | | 7 days post-xenotransplantation (xenot) into mouse (N = 3) | iMG EC11 (N = 5) |
| | | | 12-day xenot (N = 4) | iMG EC11 (N = 5) |
| | | | 8-week xenot (N = 3) | iMG EC11 (N = 5) |
| Rheinberger M., *Cell. Rep.*, 2023 | iMG (C20) | GSE205807 | HIV activated (N = 2) | Uninfected (N=2) |
| | | | HIV activated (N = 2) | HIV latent (N = 2) |
| | | | HIV latent (N = 2) | Uninfected (N=2) |

**AD* Alzheimer's disease; *iMG* human iPSC-derived microglia; *hMGL* human ESC-derived microglia-like cell; *IFN-**β*** interferon-***β***; *HIV* human immunodeficiency virus

**Table S2. Significant enrichment of co-bound dfCREs by EHF and FOS:JUN under *SORL1* KO in dfCRE modules (two-sided Fisher's exact test)**

| dfCRE cluster | *P* value | FDR | Odds ratio |
|---|---|---|---|
| Cluster12 | 2.19E-09 | 1.06E-08 | 6.94587755 |
| Cluster13 | 0.02614899 | 0.04739504 | 76.3878251 |
| Cluster15 | 0.00334091 | 0.00645909 | 13.7268936 |
| Cluster19 | 1.03E-12 | 7.43E-12 | 2.516379 |
| Cluster24 | 0.03638509 | 0.06206869 | 7.36645078 |
| Cluster33 | 3.30E-19 | 3.19E-18 | 2.76640487 |
| Cluster35 | 0.04970685 | 0.08008326 | 26.5598055 |
| Cluster38 | 0.00239709 | 0.00534736 | 34.0991478 |
| Cluster44 | 3.44E-11 | 2.00E-10 | 2.34736662 |
| Cluster45 | 4.68E-06 | 1.70E-05 | 1.69420851 |
| Cluster48 | 1.38E-05 | 4.00E-05 | 15.9989361 |
| Cluster50 | 3.63E-21 | 1.05E-19 | 26.2888332 |
| Cluster57 | 5.97E-20 | 8.65E-19 | 8.6743464 |
| Cluster63 | 5.84E-06 | 1.88E-05 | 5.83869666 |
| Cluster65 | 0.00033324 | 0.00080533 | 12.816127 |
| Cluster66 | 0.00276262 | 0.00572258 | 2.33957978 |
| Cluster91 | 1.40E-06 | 5.78E-06 | 7.5974642 |
| Cluster98 | 2.57E-05 | 6.78E-05 | 5.20257599 |

**Table S3 Reagents and resources:**

| Resource | Provider | Catalogue number |
|---|---|---|
| **Cell lines** | | |
| HMC3 | ATCC | CRL-3304 |
| **Reagents** | | |
| DMEM, high glucose | Gibco | 11-965-092 |
| Antibiotic-Antimycotic | Gibco | 15-240-062 |
| Fetal bovine serum | Gibco | 10-438-026 |
| Human IFNγ | ThermoFisher | AF-300-02 |
| Human TNFα | ThermoFisher | AF-300-01A |
| DPBS | ThermoFisher | 14190235 |
| RNeasy Mini kit | Qiagen | 74106 |
| iScript™ cDNA Synthesis Kit | Bio-rad | 1708891 |
| Power SYBR™ Green PCR Master Mix | Fisher Scientific | 43-687-06 |
| **Primers** | **Forward** | **Reverse** |
| *APOE* | GAGCAGGCCCAGCAGATAC | CTCCATGTCTTCCACCAGGG |
| *ATF3* | CGGAGCCTGGAGCAAAATGA | GGATGGCAAACCTCAGCTCT |
| *B2M* | AGATGAGTATGCCTGCCGTG | GCGGCATCTTCAAACCTCCA |
| *DNAJB6* | GAGCTGTGAGGAGATTCGGG | TTGTTGGAATGGGTCCGAGG |
| *IL6* | AGACAGCCACTCACCTCTTCAG | TTCTGCCAGTGCCTCTTTGCTG |
| *IL15* | ATTGTGGATGGATGGCTGCT | CTGCACTGAAACAGCCCAAA |
| *PD-L1* | TGCAGGGCATTCCAGAAAGA | ACCGTGACAGTAAATGCGTTC |
| *RELA* | GCTGCATCCACAGTTTCCAG | TCCCCACGCTGCTCTTCTAT |
| *SLC1A3* | ACATGAAGGAACAGGGGCAG | CACGGGGGCATACCACATTA |
| *ZBTB14* | ATGGAGTTCTTCCGGTCTGG | TCTCCTTCCAGGCGTTGTTC |
| **Instruments** | **Provider** | |
| Evos M7000 | ThermoFisher | |
| MiniAmp Thermal cycler | ThermoFisher | |
| ViiA7 qPCR | ThermoFisher | |
| **Softwares** | | |
| QuantStudio | ThermoFisher | |
